# Scalable Generation of Universal hiPSC-Derived Vascular Progenitor Cells for Safe and Sustained Revascularization in Chronic Limb-Threatening Ischemia

**DOI:** 10.64898/2026.01.16.699995

**Authors:** Joshua Heuslein, Haiyan Cao, Sen Chen, William Schachterle, Min-Su Kim, Bryan Sutermaster, Dmitriy Podolskiy, Alla Amcheslavsky, Richa Hanamsagar, Shankar Swaminathan, Pratik Lalit, Joseph Laning, Yongting Wang, Erin Kimbrel, Nutan Prasain

## Abstract

**Background:** Chronic limb-threatening ischemia (CLTI) is the most severe form of peripheral artery disease and can result in debilitating tissue damage, limb loss, and mortality if left untreated. Despite surgical bypass and endovascular interventions, there is high unmet need to develop novel therapies that can restore durable blood flow and rescue limb function in patients whose disease is not amenable to surgical bypass and endovascular procedures. Human induced pluripotent stem cell (hiPSC)-derived vascular progenitor cells (VPC) hold promise for addressing this unmet need, yet their clinical adoption will require a scalable and consistently high-quality cell product that can be used safely in a large number of CLTI patients.

**Methods:** Here, we report a robust, scalable GMP-adaptable platform for generating universally immuno-compatible VPC from human leukocyte antigen (HLA) class I/II-edited hiPSCs with extensive characterization of phenotypic and functional attributes critical to address key translational gaps in developing cell-based therapies for CLTI. We have interrogated their therapeutic efficacy in multiple murine CLTI models using a combination of clinically relevant endpoints, histology, and tissue-based RNAseq analysis.

**Results:** We found that VPC-treated mice exhibited significantly improved perfusion ratios and preserved limb function, reduced inflammation, and increased physiological neovascularization without pathological malformations.

**Conclusions:** Genetic modification conferring hypoimmune status coupled with a robust differentiation process enables large scale production of an “off-the shelf” high-quality VPC product with the potential to address unmet need in CLTI patients regardless of HLA status.

## Background

Chronic limb-threatening ischemia (CLTI), the end-stage manifestation of peripheral artery disease, represents a critical unmet medical need, with 1-year mortality rates exceeding 20% and major amputations occurring in nearly 30% of patients despite revascularization efforts(1–3). While current therapies, including surgical bypass and endovascular interventions, may provide relief to some CLTI patients(4,5), up to 20-30% of patients are not amenable to revascularization approaches(6–8). Furthermore, these surgical interventions fail to address the underlying microvascular rarefaction, persistent inflammation, and skeletal muscle degeneration that drive disease progression.

Over the past two decades, multiple gene therapy approaches have been evaluated in CLTI with the goal of promoting therapeutic angiogenesis in the ischemic tissue to limit major limb events (i.e major amputation) and improve ulcer healing(3). More than 15 early- and mid-phase clinical trials used plasmid or viral vectors engineered to express VEGF-A, HGF, SDF1, HIF1α or FGF1, based on their established roles in endothelial proliferation, arteriogenesis, and tissue repair(3,9). Across studies, gene transfer was generally safe and well tolerated, with no major vector-related or systemic toxicities reported. However, despite encouraging signals in some smaller phase I/II trails such as improvement in ulcer healing, rest pain, or perfusion, large randomized controlled studies have not demonstrated meaningful benefit on key clinical endpoints, including amputation-free survival or major amputation. These limited outcomes likely reflect the oversimplified strategy of delivering a single angiogenic factor in a complex, heterogenous disease driven by multiple biological pathways (10). Cell therapy, in contrast to single-factor gene delivery, offers a mechanistic advantage by providing a sustained secretion of multiple therapeutic factors to target multiple pathways relevant to CLTI pathogenesis (11–13). There is growing appreciation for the multi-faceted pathophysiology occurring in CLTI with severe implications in myopathy(14–23), metabolism(15,24,25), and inflammation(26–32), in addition to vascular insufficiency. Early, small, single-center cell therapy trials primarily using bone marrow- or peripheral blood-derived mononuclear or enriched progenitor cell populations have consistently demonstrated favorable safety profiles and signals of efficacy, typically reflected by improvement in amputation-free survival, major amputation rates, ulcer healing, and rest pain reduction (12,33–36). However, large, multi-center Phase III trials conducted in broader and more diverse populations have not shown significant benefit on some of these same primary clinical endpoints to date(37,38). While this may be in part due to clinical trial design (e.g. a heterogeneous trial population, endpoint selection) and/or non-optimal therapeutic delivery, there is a need to further enhance the underlying cell therapy product and preclinical testing.

Regenerative cell therapies have emerged as a promising strategy to restore perfusion and tissue repair in CLTI. A wide range of cell types and sources are under preclinical and clinical investigation, each with distinct advantages and limitations. Autologous bone marrow mononuclear cells (BM-MNC) and peripheral blood mononuclear cells (PB-MNC) have shown safety but inconsistent efficacy in CLTI clinical trials, likely due to the heterogeneity of MNC populations and the impaired function of MNCs in patients with diabetes, advanced age, and CLTI(12,39–42). Enrichment for endothelial progenitor-containing fractions (e.g., CD34⁺, CD133⁺) can increase the proportion of bioactive cells and reduce cellular heterogeneity(43–45). However, these approaches often require high doses of enriched population (>10_ total or >10_/kg) despite likely reduced bioavailability and impaired function of target cells in patients with diabetes, advanced age, and CLTI(46–51). Cell enrichment involves multi-day target cell mobilization, collection of white blood cells through leukapheresis and onerous cell sorting and lot release procedures(52), and yet enriched cells may remain susceptible to donor-to-donor variability hindering successful clinical application^46–50^. Meta-analyses of randomized controlled trials indicate that autologous CD34^+^ cell therapy has yet to demonstrate consistent, statistically significant and clinically meaningful benefits(51). Allogeneic, ex-vivo-expandable mesenchymal stromal cells (MSCs) can overcome some donor-related constraints, offering scalable production(53) and intrinsic immunomodulatory properties(54). Yet, placebo-controlled trials with unmodified MSCs have demonstrated modest or non-superior amputation-free survival and inconsistent ulcer healing(12,38,55), likely reflecting limited potency, short in-vivo persistence, immune rejection risks, and variability from donor differences and manufacturing processes.

Recent studies have demonstrated an additive benefits (such as greater perfusion recovery and more mature or stable neovascular networks) of co-administration of endothelial progenitor cells such as endothelial colony forming cells (ECFC) and supporting cell types (eg. smooth muscle, pericytes, MSC) in CLTI preclinical models(56–58). However, scalable production of ECFC/endothelial progenitor cells in a compliant process while addressing potential product variability and immunogenicity remains challenging.

Vascular cells derived from human induced pluripotent stem cells (hiPSC) are a promising avenue to develop a consistent product from an inexhaustible source. This also enables extensive quality control prior to off-the-shelf usage to ensure the highest quality product. Recent advances in cell engineering, including recombinant adeno-associated virus (rAAV) and CRISPR-mediated disruption of MHC Class I/II and overexpression of immune checkpoint molecules (e.g., HLA-E, CD47), now enable the creation of “universal”, off-the-shelf donor cells that evade host immunity(59–63). However, differentiating such engineered hiPSCs into therapeutically functional VPC with consistent endothelial-perivascular duality remains challenging, particularly under clinically scalable conditions. While several reports have demonstrated efficacy of hiPSC/hESC-derived vascular cells in pre-clinical CLTI models, most have lacked clinical translatability due to the use of research grade scale(64) and raw materials(65–68). Furthermore, the preclinical models themselves are often prophylactic and do not mirror the chronic nature and complex pathophysiology of human CLTI, including diabetes-mediated microvascular dysfunction.

Here, we address the gaps in both cell sourcing and translation by developing a scalable source of universally immunocompatible VPC and evaluating their efficacy in diabetic models of CLTI with clinically relevant endpoints. Our study aims to (1) establish a GMP-compatible robust differentiation protocol yielding VPC derived from universal iPSCs at scale with angiogenic, anti-inflammation and cytoprotective functions, (2) demonstrate durable perfusion recovery and improved limb function in delayed-treatment models, and (3) validate the safety and localized action limited to injection sites of these cells across immunocompetent and immunodeficient hosts.

## Materials and Methods

### Maintenance of hiPSCs and VPC differentiation

The hiPSC bank used here was derived through mRNA-based reprogramming of human dermal fibroblasts, as previously described(69) and maintained on mitomycin treated human dermal fibroblast (either ATCC, OSC-201-011 or an in-house HDF line) in StemFit media (Amsbio, Cat. No. SFB-503-GMP). Cells were passaged with VPC differentiation was performed similarly as previously described(70). Briefly, hiPSCs were lifted with Cell Dissociation Buffer (CDB, Gibco, 13151-014). and resuspended in fresh medium Stemfit containing 10µM of Y27632 (Fujifilm, Cat. No. 252-00701). Cells were seeded on human collagen IV-coated flasks (Advance Biomatrix, Cat. No. 5022) and incubated at 37°C in normoxic conditions (5% CO_2_, 20% O_2_) for 24 hr. Media was changed to StemSpan (STEMCELL Technologies, Cat. No. 100-0130) supplemented with 50ng/mL of human VEGF165 (Proteintech, Cat. No. HZ-1038-GMP), 50ng/mL of human FGF2 (Proteintech, Cat. No. HZ-1285-GMP) and 25ng/mL of human BMP4 (Proteintech, Cat. No. HZ-1045-GMP). Cells were grown for 6 days under hypoxic conditions (5% CO_2_, 5% O_2_). On day 6, cells were harvested with TryplE Select (Gibco, Cat. No. 12563029) to make a single cell suspension and frozen in Cryostor® CS10 (STEMCELL technologies, Cat. No. 07930). For extended culture VPC (EXT-VPC), which represent the full 10-day differentiation protocol, day-6 VPC were thawed in complete StemDiff Endothelial Expansion Media (STEMCELL Technologies, Cat. No. #08007) and seeded on human Collagen IV coated vessels. Cells were grown for 4 additional days under hypoxic conditions then EXT-VPC were harvested with TryplE Select to make a single cell suspension and frozen in CS10.

### scRNA-sequencing of VPC lots

Cryopreserved samples were thawed and library preparation was performed using a 10X Genomic Chromium 3’ gene expression RNA-seq library preparation and 10x library sequencing on the Illumina HiSeq Series. Reads were aligned to human reference using the CellRanger pipeline (10x Genomics). scRNAseq analysis was performed with Seurat version 4.3.0.1(71) in R version 4.3.0. Cells with low gene number, high percentage of mitochondrial transcripts, or potential doublets with high UMI counts were excluded. After initial quality filtering, an integrated reference sample set was generated using reciprocal PCA (RPCA). Briefly, normalization and identification of variable features were found individually for six representative VPC lots which were then using the RPCA algorithm while regressing cell cycle genes during scaling. Additional query datasets (5 additional VPC lots, HUVEC, and PSC lots) were mapped to this reference(71,72).

### VPC surface marker expression by flow cytometry

For flow cytometry analysis, previously prepared frozen single cell vials were thawed into FACS buffer (2% FBS in PBS). Cell surface staining was analyzed with a SONY spectral analyzer using the antibodies outlined in Supplementary Table 4. Isotype controls and negative cell samples were used as controls to establish a threshold for positive staining and subset gating.(59)

### Secretome analysis of VPC conditioned media

Cryopreserved VPC were thawed and seeded at ∼50,000 cells /cm^2^ on fibronectin coated 6-well plates (Corning, 354402) in complete Vasculife VEGF Endothelial medium (Lifeline Cell Technology, LL-003) for 24-hours. After 24-hours, media was changed to “triple-depleted” Vasculife (i.e. Vasculife without VEGF, FGF2, or FBS) and cultured for an additional 48-hours.

Media was collected, centrifuged to remove any dead cells, and supernatants were analyzed using the Angiogenesis Array G1000 (Cat. No AAH-ANG-G1000-4) which was scanned and quantified at RayBioTech or with the following human Quantikine ELISA (R&D Systems) kits according to manufacturer’s instructions: HGF (DHG00B), CCL2/MCP-1 (CDP00), ANGPT2 (DANG20), ANGPT1 (DANG10), FGF2/bFGF (DFB50), CXCL12/SDF1 (DSA00), FST/follistatin (DFN00), ANG/angiogenin (DNA00), MMP2 (DMP2F0), EGF (DEG00) and ANGPTL4 (Invitrogen, EHANGPTL4).

### Endothelial cytoprotection

Primary, pooled donor HUVEC (Promocell, C-12203) were labeled with CellTrace-Far Red (ThermoFisher, C34564) at 1:100 in PBS for 20min at 37°C. Following incubation, HUVEC were quenched and washed with excess FBS (10%). Labeled HUVEC were then plated in fibronectin-coated well of 6-well plates (100K/well, ∼10K/cm^2^). Cryopreserved VPC were thawed, counted, and seeded at 50K/cells per well of 6-well plate containing labeled HUVEC (∼10K/cm^2^) in 3mL Complete Vasculife media for 24-hours. After 24-hours, media was changed to depleted media (i.e Vasculife without FBS supplementation) for all conditions except positive control, which received Complete Vasculife growth media. Cells were cultured for an additional 96-hours in 5% O_2_ before being harvested with Accutase, stained with AnnexinV / 7-AAD (Invitrogen, 50-927-7) according to manufacturer’s instructions and quantified on the Novocyte Advanteon (Agilent).

For each experiment, VPC lot(s) were tested against the following controls: Complete Vasculife only, depleted Vasculife only (control), and unlabeled HUVEC.

### HUVEC tube stabilization

Growth factor-reduced Matrigel (Corning, 354230) was added (290uL/well) to 24-well plate and incubated at 37°C for 20-30 minutes to gel. GFP-HUVEC (Angio Proteome, cAP-0001GFP) were then seeded onto the Matrigel at 22.2K/cm^2^/well (40K/well) in 500uL complete Vasculife media and cultured in 1% O_2_ for 24-hours to enable for self-assembly of HUVEC tubes (Day 0). After 24 hours, GFP-HUVEC networks were imaged in a 3×3 grid at 4x with a DMi8 epifluorescence microscope (Leica) to assure sufficient network assembly. Cryopreserved VPC were then thawed, counted, and overlaid on top of GFP-HUVEC tube networks at 22.2K/cm^2^/well (40K/well) in 500uL depleted Vasculife (i.e. Vasculife not supplemented with FBS, VEGF, or FGF2) (Day 1). Plates were incubated in 1% O_2_ for 6 additional days (Day 7) without media changes. On Day 7, wells were imaged as described previously. Optionally after initial imaging, networks were stained using pan-cell stain, CellTracker Orange (ThermoFisher, C34551). Briefly, wells were incubated with 10uM staining solution in 500ul depleted Vasculife for 45 minutes in normoxia at 37C. Wells were then quenched and washed 2-3 times in complete Vasculife (i.e. FBS containing), fixed with 4% PFA for 20 minutes, and imaged. Total network tubes, total tube length, covered area, branch points, and loops were quantified on both GFP-HUVEC and phase contrast images using custom Wimasis WimTube software. (Onimagin Technologies, Cordoba, Spain).

### Mice

Animal studies were approved by internal review boards at either Invivotek (Hamilton, NJ, USA) or Pharmaseed (Ness Ziona, Israel) and conformed to recommendations in the *Guide for the Care and Use of Laboratory Animals* (US National Institutes of Health) and Association for Assessment and Accreditation of Laboratory Animal Care (AAALAC) guidelines. Male athymic Balb/c nude mice (Charles River, CAnN.Cg-Foxn1^nu^/Crl, #194) and db/db (JAX, BKS.Cg-Dock7^m^+/+Lepr^db^/J, #000642) aged 7-12 weeks were used for all studies, except non-GLP 9-month safety study in which both male and female Balb/c nude mice were used. For db/db mice, blood glucose levels were measured prior to surgery and weekly until study completion using a standard glucometer. Only mice with blood glucose levels ≥ than 250 mg/dL were used for study.

### Hindlimb ischemia model

Hindlimb ischemia surgery was performed at Pharmaseed LLC as previously described(73–75) and in compliance with “The Israel Animal Welfare Act” and after “The Israel Board for Animal Experiments” had approved it. Briefly one hour before the surgery, Meloxicam was administered (5 mg/kg SC), and mice were anesthetized by 4% isoflurane in a mixture of 70% N_2_ and 30% O_2_ and maintained with 1.5-2% isoflurane. The right femoral artery was ligated with 6-0 silk thread and transected between the two ligatures. The wound was closed with 5-0 Vicryl absorbable thread and the mouse was allowed to recover. Meloxicam was administered (5 mg/kg SC) for the next two days. All animals received Enrofloxacin (Baytril) in the drinking water (0.2mg/mL) for two weeks to prevent infection and wet food to prevent weight loss. Buprenorphine SC at 0.02 mg/kg was given twice a day to animals that exhibited suffering. Body weight, limb function(76), and limb necrosis(77) were evaluated twice a week or daily in animals exhibiting ≥10% reduction in body weight.

### Laser Speckle Contrast Imaging (LSCI)

Blood flow in both legs for each mouse was measured using laser speckle contrast imaging (LSCI) device (moor-FLPI-2, Moor Instruments), before surgery and immediately after surgery. Animals were stratified into groups in balanced fashion according to the blood-flow results. Further measurements were performed weekly and were expressed as the ratio of the flow in the ischemic limb to that in the normal, unoperated left limb.

### Non-GLP 9-month safety study

The non-GLP 9-month study, clinical hematology, and clinical chemistry in athymic Balb/c nude mice were conducted at Invivotek (Hamilton, NJ). Vehicle, 1 million (1M) VPC, or 2 million (2M) VPC were injected into 2 sites in the quadriceps of 8-10 week old male and female Balb/c nude mice. Naïve (age, sex matched non-injected animals) were kept as an additional control.

Animals were observed up to 9-months following cell injection and monitored biweekly for body weight. At 9-months, animal health was assessed by a Board Certified Veterinarian, as well as routine clinical hematology and automated clinical chemistry from peripheral blood samples.

### DiR labeling, IVIS Imaging, and Biodistribution

All biodistribution studies were performed at Invivotek (Hamilton, NJ). Cryopreserved VPC were thawed, assessed for viability, and stained with 10µM IVISense DiR 750. Briefly, after thaw and wash, cells were resuspended in pre-warmed PBS (2 million cells / mL) with 10µM IVISense DiR 750 (Revvity, Cat. No. 125964) and incubated at 37°C for 20 minutes. After 20 minutes, cells suspension was quenched with 10x excess volume of complete Vasculife media (with FBS) then washed with complete Vasculife media 2x. Cells were then resuspended in a sterile isotonic buffer (proprietary) at 2 million cells/ 100µL. VPC or vehicle were injected into the quadriceps (50µL) and the gastrocnemius muscle (50µL) of the left limb of either naïve athymic Balb/c nude mice (male, 10-12 weeks, Charles River) or those that were subjected to hindlimb ischemia surgery 24-hours before. 10K VPC were also reserved, plated into a 96-well plate, and imaged immediately following labeling to confirm successful labeling. Animals were anesthetized via isofluorane and the ventral aspect was imaged using an IVIS Spectrum (Revvity). For in-plate, whole body, and ex vivo imaging 710nm excitation and 760nm emission filters were used. Following ex vivo imaging, tissues were snap-frozen and stored at −80C until further hAlu PCR analysis.

### Treadmill running

Exercise test for maximal walking distance via treadmill was performed using the UGO Basile Rodent Treadmill (UGO Basile, Cat. No 47300/47350) before surgery, on Day 21 pre injection and Day 84/85. Two days prior to testing, all animals were subjected to an acclimatization period at 1m/min for 5 minutes. On the day of testing, animals started at 1 m/min where speed was increased by 1 m/min every 30 seconds, until reaching 5 m/min. Then speed was increased by 1 m/min every minute until reaching a maximum of 10 m/min. Animals were allowed to run until exhaustion which was defined as an accumulation of total electric pulses exceeding 3 seconds or total of 30 shocks. The average of two runs was recorded.

### StO_2_ measurement

Oxygen saturation was measured non-invasively on the foot pad of both the ischemic and contralateral, non-operated limb on Days: 21 (pre-cell injection), 43 and 85 using the mooR-VMS Oxy (moor Instruments, United Kingdom). Readings were performed under 1.5-2% isoflurane in a mixture of 70% N_2_O and 30% O_2_. The average of three readings was used per animal per timepoint.

## Results

### Generation of GMP-compatible, scalable, universal hiPSC-derived VPC

To develop a universal, off-the-shelf allogeneic vascular progenitor cell product, we utilized universal donor human induced pluripotent stem cells (hiPSCs) as previously described(59–61,63,70). These cells have been genetically edited such that MHC Class I and Class II are knocked out with overexpression of HLA-E to deter lysis by NK cells. Additionally, these hiPSCs have been edited to include a thymidine kinase (TK) suicide gene as an additional safety switch.

Here, we developed a modified hemogenic endothelium (HE) protocol to produce vascular progenitor cells (VPC) in a scalable format with compliant materials within 6 days (Figure 1A, Supplemental Figure 1). These VPC can additionally be replated either directly post-harvest or after freeze/thaw to generate EXT-VPC (extended-culture VPC), which maintain viability and function while aiding in the elimination of any residual undifferentiated or partially differentiated PSC-like contaminants. Single-cell RNA-sequencing (scRNA-seq) of six VPC lot productions confirmed the consistency of cell phenotype across a variety of formats and operators (Figure 1B). Using the integrated sample set of these six VPC scRNA-seq samples to act as a reference for additional scRNA-seq analysis, we mapped an additional 5 VPC lots, as well as both HUVEC and iPSCs to further interrogate subpopulations by scRNA-seq (Figure 1C). Endothelial (PECAM1^+^ CDH5^+^) and mesenchymal/perivascular (HAND1^+^ ACTC1^+^ KDR^+^) phenotypes were the primary subpopulations with minimal residual mesoderm-like (APLNR^+^ CDH2^+^ KDR^+^ DLK^+^) or endoderm-like (EPCAM^+^SOX17^+^APJ^-^) populations and no overlap with iPSCs (POU5F1^+^ SOX2^+^) based on established subpopulation gene marker expression (Figure 1C, Supplemental Figure 2). Freeze-thaw and extended culturing (EXT-VPC) further purified the two dominant subpopulations and increased the maturity of the endothelial population shown by closer proximity to HUVEC (Figure 1B,C). Flow cytometry further confirmed endothelial and mesenchymal/perivascular subpopulations as the dominant phenotypes in VPC. Across multiple operators and scale-up formats (T175 flasks, 2-layer, 5-layer CellStack) VPC are approximately 50-60% CD144+/VE-Cadherin+ (endothelial) and 25-35% CD140b/PDGFRb+ (perivascular). EXT-VPC not only maintain the overall endothelial phenotype with 50-60% CD144+/VE-Cadherin+ but the endothelial maturity also tends to increase with extended culture seen by 2-fold decrease in early HE marker (CD144+CD43+, 33.4% in VPC vs 6.9% in EXT-VPC in T175 format) and 2-fold increase in CD144+CXCR4+ cells (11.1% in VPC vs. 34.6% in EXT-VPC in T175 format), consistent with a heighted arterial endothelial phenotype (Figure 1E). Interestingly, the endothelial subpopulation of both VPC and EXT-VPC was also CD271+, which is typically considered a stromal cell marker but has been reported to be co-expressed with CD31 in rare in-vivo population termed regeneration associated endothelial cells (hRECs)(78,79).

**Figure 1:**
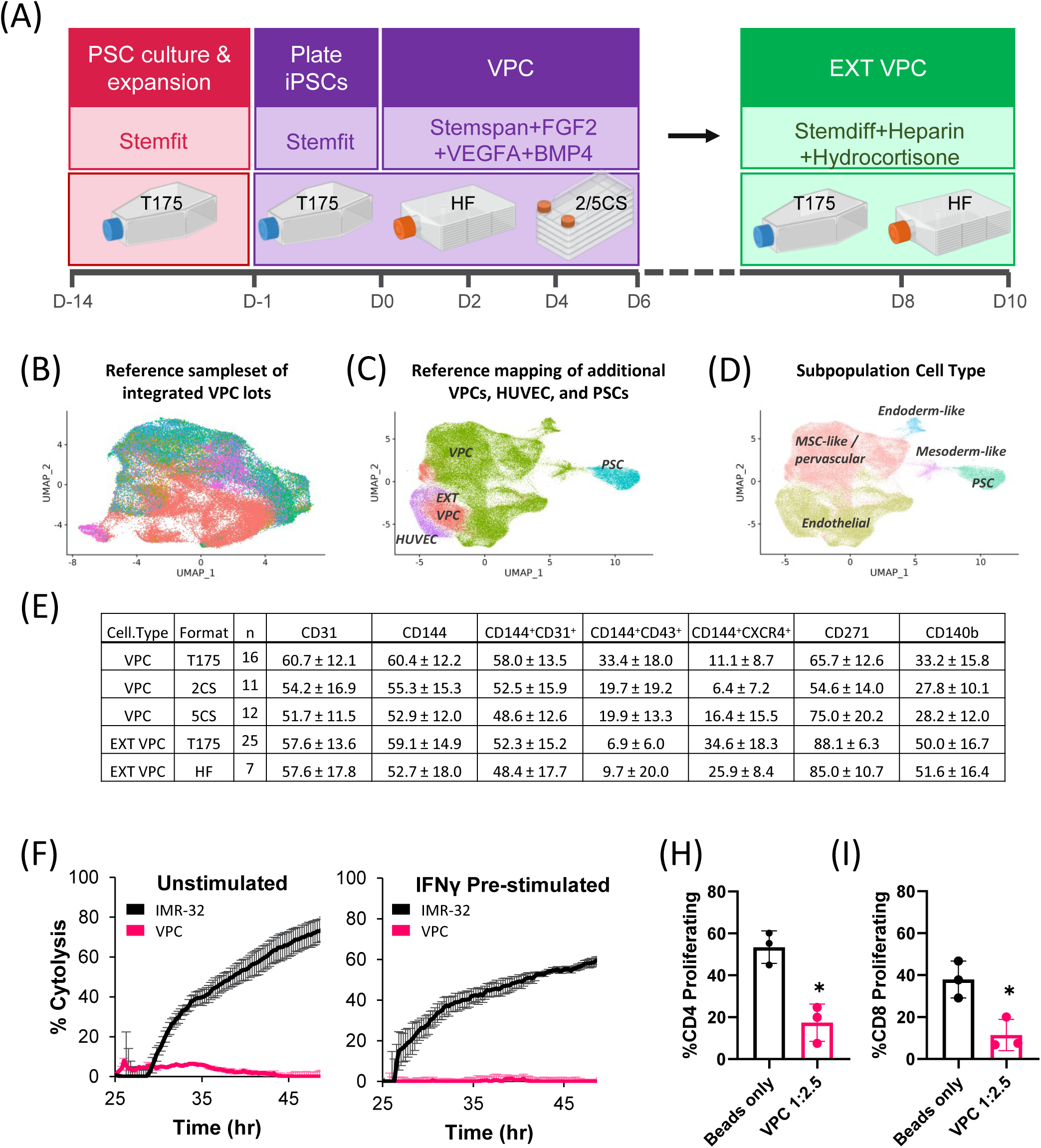
Generation of cGMP-compliant, scalable hiPSC-derived vascular progenitor cells (VPC). (A). Differentiation protocol of passagable VPC across several scale-up formats (B) UMAP plot of integrated analysis of scRNA-seq datasets from 6 independent VPC lot productions (>58K cells) to create a reference sample for additional scRNA-seq analyses (C,D) scRNA-seq reference mapping of additional 5 VPC lots, including compliant, scale-up, and extended culture VPC as well as non-VPC reference cell lines (>100K cells total) labeled by cell type (C) and subpopulation (D) based on phenotypic marker expression. (E) Summary of spectral flow cytometry analysis of key surface markers to define identity and purity across independent VPC lot productions in various scale-up formats (2CS = 2-layer CellStack; 5CS = 5-layer CellStack; HF = HyperFLASK). Data is mean % cells ± standard deviation. (F) Natural killer (NK) cell cytolysis with and without 8ng/mL IFNγ, n=3 technical replicates / group/per timepoint. (H) T-cell activation by flow cytometry after PBMC co-culture with VPC (1:2.5 VPC:PBMC) and stimulation with 8ng/mL IFN and CD3/CD28 beads for 72 hours. Data is mean +/- SEM, n=3 with separate PBMC donors.*p <0.05 vs. beads only; unpaired Student’s *t*-test.

The universal editing technology we used here has been previously reported(59). We confirmed absence of surface expression of MHC Class I (HLA-ABC) and MHC Class II (HLA-DP, DQ, DR) on differentiated VPC by flow cytometry. As expected, HLA-E was expressed and up-regulated upon IFNγ stimulation (Supplemental Figure 3). Notably, VPC expressed several complement inhibitors (CD46, CD55, CD59) and minimally expressed CD142/TF, several co-stimulatory factors (CD80, CD86), E-selectin, P-selectin, and FasL. Upon IFNγ stimulation, VPC up-regulated several inhibitors of T-cell activation (CD274/PDL1 & CD273/PDL2). IFNγ stimulation also increased ICAM-1 expression, indicative of a normal endothelial response to inflammation (Supplemental Figure 3). Functionally, VPC were able to avoid NK lysis both with and without IFNγ stimulation compared to the positive control cell line, IMR-32, a neuroblastoma cell line that undergoes lysis by NK cells. (Figure 1F). Additionally, VPC reduced proliferating CD4+ (Figure 1H) and CD8+ (Figure 1I) T-cells when co-cultured with PBMCs stimulated with CD3/CD28 beads + IFNγ.

### VPC exhibit cytoprotective, stabilizing effects in vitro

We next wanted to assess the VPC secretome. An initial screen of 43 human angiogenic factors revealed that several angiogenic analytes were consistently secreted in VPC conditioned media, including MCP-1/CCL2, ANGPT2, ANGPTL4, HGF, Follistatin (FST), and Angiogenin (ANG) (Supplemental Figure 4). We followed up on this initial screen with multiple ELISAs to confirm the expression of multiple proteins critical in angiogenesis, arteriogenesis, wound repair, myocyte metabolism, and immune modulation including HGF, CCL2, ANGPT1, ANGPT2, FGF2, SDF1a, FST, ANG, MMP2, ANGPTL4, and EGF. Secretion profiles were consistent between VPC and EXT-VPC with increased levels of ANGPT2 (2022 ± 325 vs. 1085 ± 150 pg/mL in EXT vs VPC, p<0.05), ANGPT1 (346 ± 93 vs. 56 ± 8 pg/mL in EXT vs VPC, p<0.05), and ANGPTL4 in EXT-VPC (436 ± 61 vs. 52± 25 pg/mL in EXT vs VPC, p<0.05) (Figure 2A). VPC did not express pro-inflammatory cytokines IL6, IL8, or IP-10/CXCL10 which have been correlated to decreased rates of amputation-free survival in CLTI patients(28) (Supplemental Figure 4).

**Figure 2:**
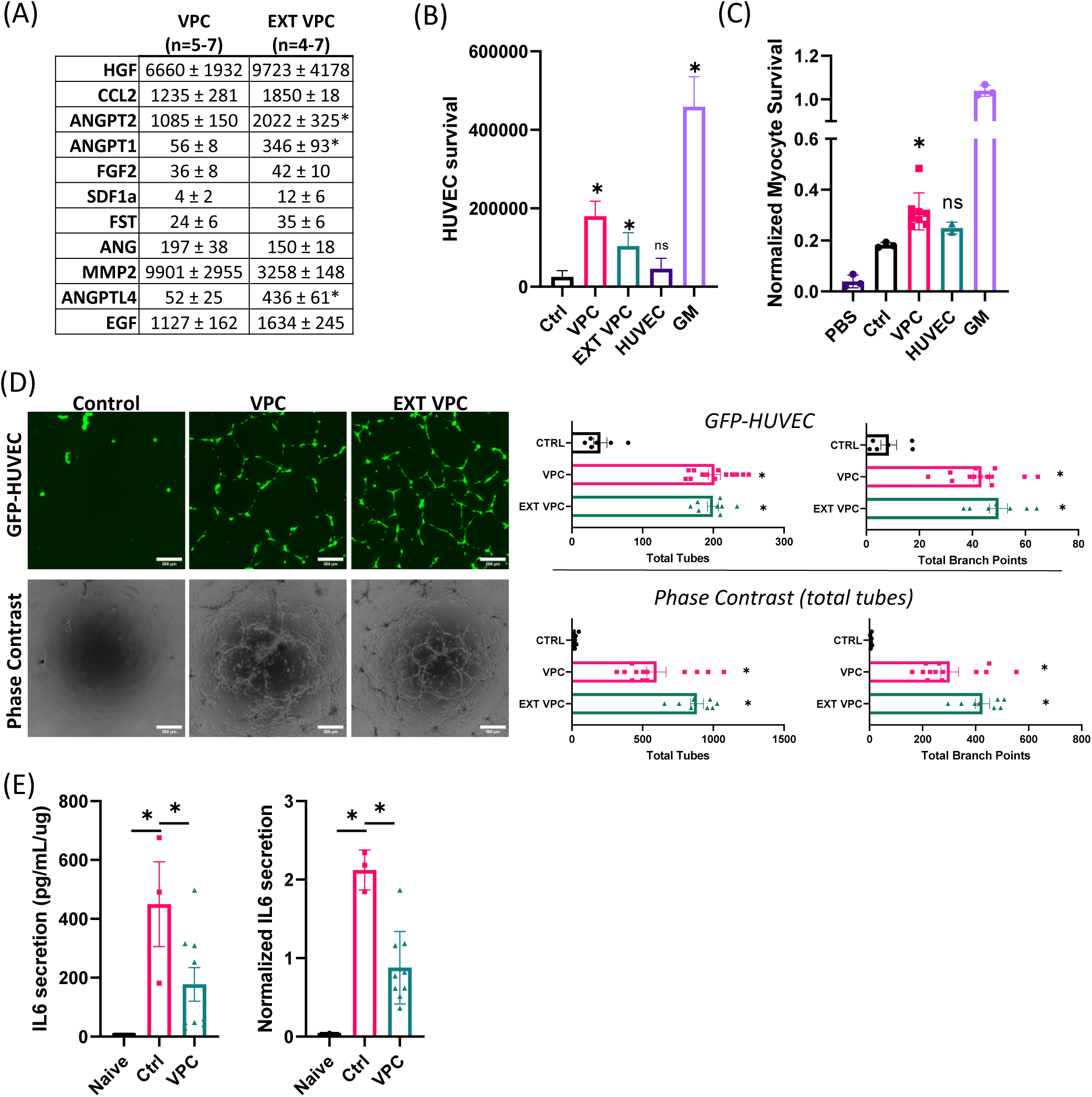
VPCs secrete numerous angiogenic, myogenic, and immunomodulatory factors, reduce IFN induced inflammation, and exhibit cytoprotective, stabilizing effects on endothelial cells and myocytes in-vitro. (A) Summary of secreted protein expression in VPC conditioned media by ELISA. Data is mean concentration (pg/mL) ± standard deviation in VPC and extended culture VPC (EXT VPC). *p<0.05 vs. VPC, Student’s *t*-test. (B) Endothelial cytoprotection indicated as live HUVEC counts following co-culture with VPC (1:2 HUVEC:VPC) and AnnexinV/7-AAD staining. Data is mean ± standard deviation, n=17 for ctrl & GM, n=16 for VPC, n=15 for EXT VPC, and n=16 for HUVEC and represent independent biological repeats. GM=complete Vasculife growth media; *p<0.05 vs ctrl; One-Way ANOVA, Tukey’s post-hoc. (C) Normalized myocyte survival with VPC conditioned media after 7 days in 1% hypoxia and serum starvation; *p<0.05 vs ctrl; One-Way ANOVA, Tukey’s post-hoc. N=3 for PBS, ctrl, and complete Vasculife growth media (GM), n=2 for HUVEC, n=8 for VPC and represent independent biological repeats. Data is mean ± SEM (D) Tube stabilization of pre-formed vascular networks by VPC quantified by total tubes and total branchpoints of both GFP-HUVEC+ and phase contrast images. Note phase contrast images demonstrate incorporation of VPCs into the HUVEC network. Data is mean ± SEM. (E) IL6 secretion 48-hours after co-culture of VPC with THP1 monocytes (1:2 ratio) with 20ng/mL IFNγ stimulation as pg/mL/ug total protein. *p<0.05 vs ctrl; One-Way ANOVA, Tukey’s post-hoc, n=3 (naïve), n=3 (ctrl, THP1 + IFN (no VPC)), n=9 (VPC+THP1+IFN) biological replicates

Interestingly, VPC did not express VEGF-A (Supplemental Figure 4), a potent regulator of vascular growth, though it has been suggested VEGF-mediated vascular growth occurs via inflammation and glycolysis which may not be ideal for CLTI patients(80).

Next, we sought to determine if VPC were able to protect endothelial cells and/or myocytes against hypoxia induced cell death. To do this, we developed an endothelial cytoprotection assay whereby labeled HUVEC were co-cultured with VPC for four days in depleted media and hypoxic (5% O_2_) culture then live HUVEC were counted by flow cytometry following AnnexinV / 7-AAD staining (Figure 2B). VPC demonstrated a significant (7-fold for VPC and 4-fold for EXT-VPC) increase in EC survival over control, across multiple independent VPC lot productions spanning multiple operators and scale-up formats. Interestingly, additional HUVEC instead of VPC at the same density did not increase labelled EC survival compared to control (p=0.67), suggesting unique cytoprotective properties of VPC (Figure 2B). As skeletal muscle survival is another vital aspect for a putative CLTI therapeutic, we sought to assess the cytoprotective ability of VPC on human skeletal muscle myocytes (HuSkM). Myotubes from differentiated myoblasts were co-cultured with VPC conditioned media in hypoxic (1% O_2_) and serum starvation conditions for seven days then assessed for viable cell count. Similar to the endothelial effect, VPC increased myotube survival in severe hypoxia and starvation conditions (1.72-fold VPC vs. control, p<0.05) whereas HUVEC did not significantly improve myotube survival (p=0.40). (Figure 2C).

As vascular rarefaction and endothelial dropout is a consequence of prolonged hypoxia in CLTI patients, we sought to evaluate VPC on the ability to promote stabilization of pre-formed vascular constructs and prevent their dropout. To do this we adapted the common tube formation assay whereby we enabled for robust tube formation of GFP-HUVEC to occur on growth factor reduced Matrigel for 24 hours. We then overlaid VPC on these networks and cultured in hypoxia (1% O_2_) and serum starvation for an additional seven days (Supplemental Figure 5). Both VPC and EXT-VPC protected GFP-HUVEC vascular networks (total tubes and total branches) to a similar degree whereas they were nearly eliminated without VPC addition (Figure 2D, Supplemental Figure 5). This was even more apparent by examining total network by phase contrast imaging (GFP-HUVEC + incorporated VPC) as VPC incorporated into the pre-existing network, both acting as a supporting cell and as cell replacement for regions where GFP-HUVEC had dropped out (Figure 2D, Supplemental Figure 6). Interesting, the VPC can also adopt a “sprouting” phenotype with nascent, small outgrowths from pre-existing tubes despite severe hypoxia and growth factor reduced conditions (Supplemental Figure 5-6).

In addition to the cytoprotective and stabilization effects VPC exerted on endothelial cells and myocytes, we sought to test the anti-inflammatory function of VPC. VPC were co-cultured with THP-1 monocytes in pro-inflammatory conditions for 2 days then assessed for total IL-6 secretion. VPC co-culture significantly lowered IFNγ induced IL-6 secretion from THP-1 cells (2-fold vs. THP1-only control, p<0.05) (Figure 2E). Similar results were seen using a higher IFNγ concentration (50pg/mL) or LPS (100pg/mL) (data not shown).

### VPC significantly improve limb perfusion in common preclinical models of CLTI

Given the consistent cytoprotective, immunomodulatory, and secretion of multiple angiogenic growth factors *in vitro*, we sought to test VPC functionality in the most common preclinical model of CLTI, the murine hindlimb ischemia (HLI) model(74,77,81). Blood flow was assessed by laser speckle contrast imaging (LSCI) following injection of VPC (1 million cells) into the quadriceps muscle of both immunocompromised athymic Balb/c nude (Figure 3A) and immunocompetent, diabetic db/db mice (Figure 3B) immediately following HLI. VPC were able to significantly improve blood flow compared to vehicle control (p<0.0001) as early as Day 14. Blood flow continued to increase over time and VPC treated animals exhibited a sustained improvement over vehicle treated animals at Day 64 post-HLI (1.7x in Balb/c and 1.65x in db/db, p<0.0001). In diabetic animals, VPC treated animals exhibited increased vascular density given by increased αSMA^+^ arteriole density in the quadriceps muscle at Day 64 post-HLI (Figure 3C-D). Additionally, VPC improve the skeletal muscle survival and morphology at the injection site. To this end, VPC reduced the total amount of fibro-adipose tissue (Figure 3D), demonstrated more mature myofibers indicative of intact dystrophin staining (Figure 3C) without centrally located nuclei (Supplemental Figure 7), unlike vehicle controls which still had weak/broken dystrophin (Figure 3C) staining and myofibers with centrally located nuclei (Supplemental Figure 7). As noted previously, EXT-VPC exhibit an *in vitro* phenotypic and functional profile having increased endothelial cell maturity, increased purity (i.e. only two subpopulations), and improved safety profile as compared to VPC due to the further elimination of partially differentiated cells through additional processing and additional days of culture. EXT-VPC were also tested in the HLI model in db/db mice in which we also observed a significantly increased blood flow recovery following EXT-VPC injection compared to VPC in HLI-treated mice (Supplemental Figure 8).

**Figure 3:**
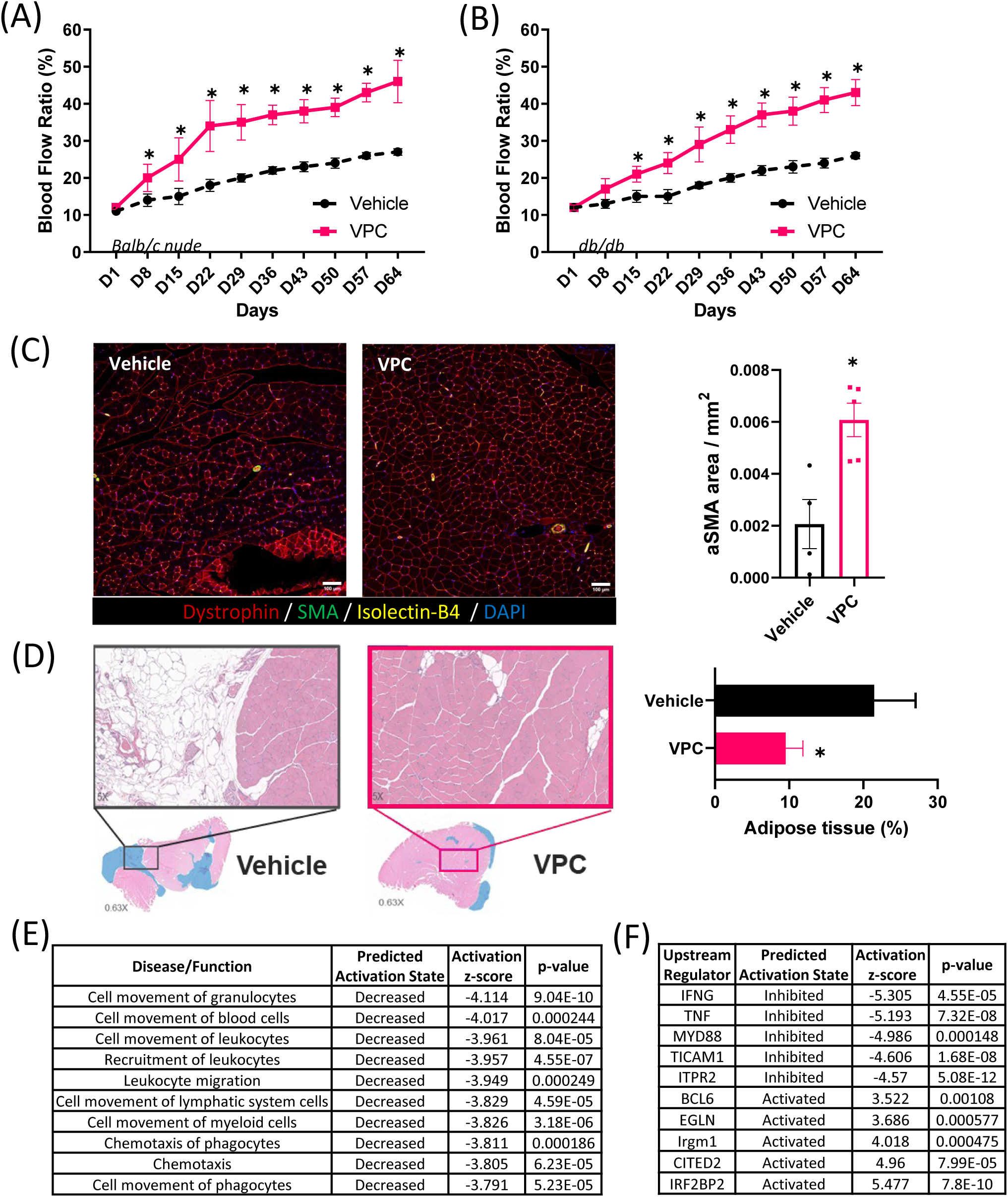
VPC significantly improves limb perfusion in both athymic nude and db/db mice. (A) Laser speckle contrast imaging (LSCI) of Balb/c nude mice following HLI and treatment with VPC or vehicle control on same day as HLI surgery. Data is mean ± SEM, n= 12 mice / group. *p<0.05 vs. vehicle; Two-way ANOVA, Tukey post-hoc test (B) LSCI of db/db mice following HLI and treatment with VPC or vehicle control on same day as HLI surgery. Data is mean ± SEM, n= 15-16 mice / group. *p<0.05 vs. vehicle; Two-way ANOVA, Tukey post-hoc test (C) Representative immunofluorescence images of quadriceps muscle at Day 64 post HLI/cell injection in db/db mice. Quantification of aSMA+ vessel density analyzed 4-5 10x images per animal in blinded fashion. Data is mean± SEM per individual animal, n=4-5 mice/group. *p<0.05 vs. vehicle; Student’s *t*-test (D) Whole slide H&E imaging of quadriceps muscle at Day 64 post HLI/cell injection in db/db mice. Blue area indicates fibroadipose tissue and pink indicates viable muscle tissue. Quantification is mean± SEM with n=4-5 mice / group. *p<0.05 vs. vehicle; Student’s *t*-test (E,F) Top 10 differentially regulated (E) disease/function pathways and (F) predicted upstream regulators by activation-score of Ingenuity Pathway Analysis of RNA-seq from quadriceps muscle at Day 64 post HLI/cell injection in db/db mice in VPC vs. vehicle. Decreased activated state or negative activation z-score indicates predicted decreased activity in VPC-treated vs. vehicle.

RNA-seq pathway analysis of quadriceps muscles from these db/db animals strongly implicates a persistent anti-inflammatory effect of VPC treatment following HLI. To this end, nearly all the most differentially regulated disease/function pathways by activation z-score were related to cell movement of immune cells and were predicted to exhibit decreased activation (Figure 3E, Supplemental Table 1). Furthermore, two of the most strongly predicted upstream regulators, IFNγ and TNFα, were predicted to be inhibited with VPC treatment (Figure 3F, Supplemental Table 2). Interestingly, the “insulin-dependent diabetes mellitus”, “diabetes mellitus”, and “glucose metabolism disorder” pathways were highly significant (p<10^-8^) with VPC treatment vs. vehicle and predicted to be downregulated (Activation z-scores of −1.863, −2.301, and −2.51, respectively) (Supplemental Table 1). While there was no significant decrease in non-fasting glucose (Supplemental Figure 9) nor do we propose VPC as a diabetes treatment per se, it does support the functionality and anti-inflammatory effects of VPC in a severely diabetic model.

### VPC are well tolerated and exhibit a limited extent of long-term persistence in vivo

We next wanted to examine the biodistribution and confirm the safety profile of these VPC in immunocompromised mice. We injected vehicle, 1 million VPC, or 2 million VPC into the quadriceps muscle of athymic Balb/c nude mice without induction of any additional injury model for up to 9-months. There were no significant alterations in body weight, animal welfare, clinical pathology (hematology & clinical chemistry), and clinical histopathology was within range of control animals across 11-tissues reviewed by independent histopathologist (Supplemental Figure 10).

To track VPC biodistribution *in vivo*, VPC were labeled with IVISense DiR 750 and injected into both the quadriceps and gastrocnemius muscles of one limb in athymic Balb/c nude mice either without surgery (VPC group, green) or in mice subjected to hindlimb ischemia (HLI) surgery the day prior (VPC+HLI, pink) and compared to untreated, naïve animals. Whole-body IVIS imaging was performed pre-injection and days 3, 8, 14, 35, and 64 after cell injection. All animals exhibit DiR+ signal in VPC-injected limbs up to Day 14. Interestingly, more cells appear to persist longer in the hypoxic environment present following HLI injury than in naïve mice (Figure 4B). However, by Day 35, both the number of animals exhibiting DiR+ signal (66% DiR+ in VPC, and 83% DiR+ in VPC+HLI animals) and the total signal magnitude significantly decreased and by Day 64 no animals exhibited DiR+ signal above background (Figure 4A-B). Throughout the study, a subset of animals were removed from the VPC+HLI group to perform *ex vivo* imaging to increase imaging resolution and determine organ-specific distribution. As expected, there was significant signal in VPC injected tissues, the quadriceps and gastrocnemius muscles, that decreased over time. In contrast to the whole-body imaging, ex vivo imaging indicated presence of DiR+ cells in the gastrocnemius even out to Day 64 whereas a signal was at background in the quadriceps between Day 7-14 (Figure 4C). This is again consistent with the hypothesis that VPC persist to a greater extent in hypoxic environments as the gastrocnemius is known to be the more hypoxic of the two muscle groups following HLI(82–84). Importantly, there was no significant off-target migration and VPC remained at the sites of injection (Figure 4C).

**Figure 4:**
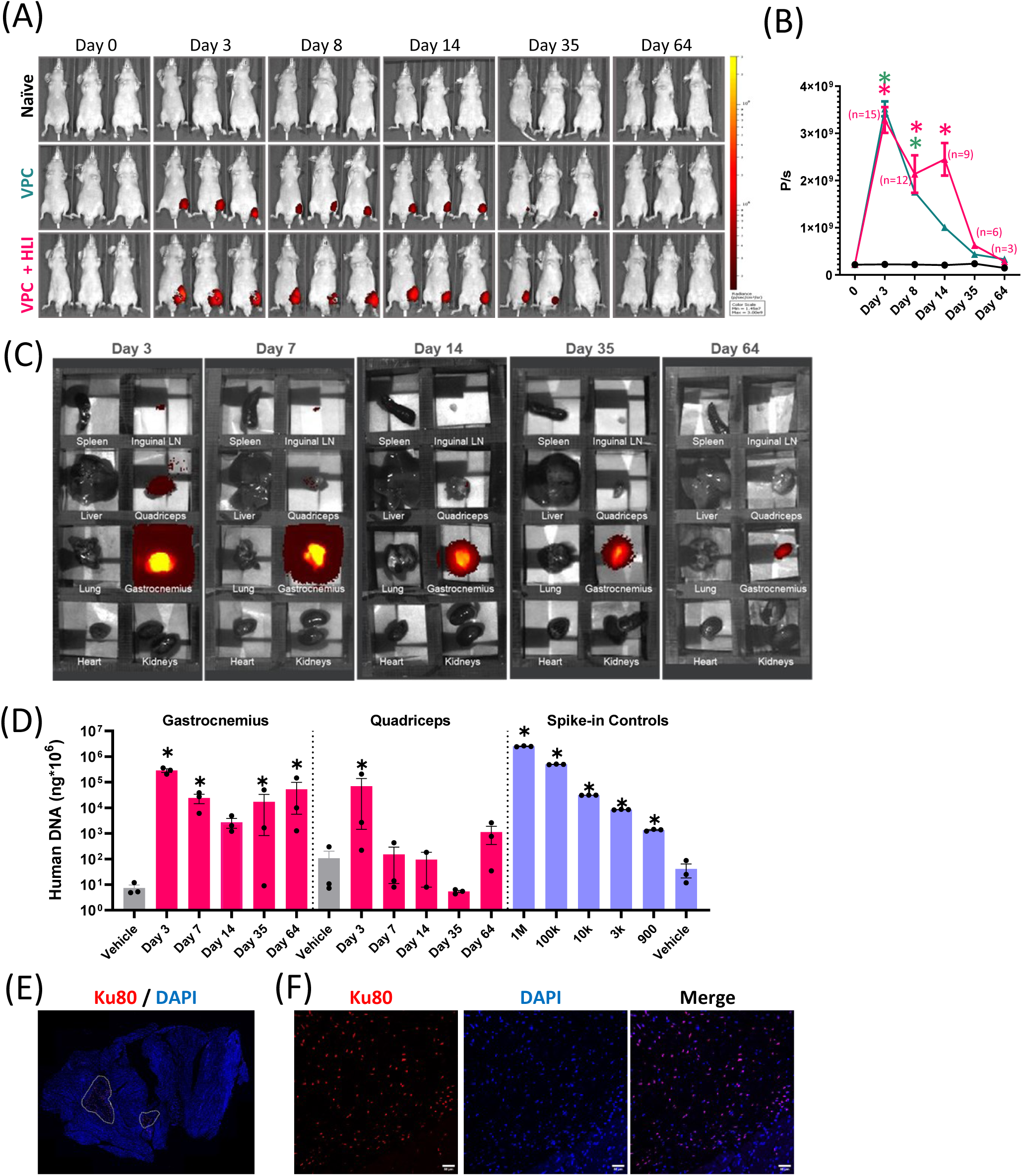
VPC are well tolerated and exhibit a limited extent of long-term persistence in-vivo. (A) Representative images of sequential whole body IVIS imaging of DiR-labeled VPC injected into quadriceps and gastrocnemius muscles of naïve Balb/c nude mice or 24-hours following HLI surgery. (B) Quantification of whole body IVIS imaging (photon/second, P/s) over time. *p<0.05 vs. naïve control Two-way ANOVA then Dunnett’s test; n=3 (vehicle), n=3 (VPC), n≥3 (VPC+HLI); Number of animals decreased over time in VPC+HLI group due to planned removal for subsequent *ex vivo* organ analysis. (C) Representative *ex vivo* IVIS images of VPC+HLI group across 8 different tissues at various time after HLI surgery/cell injection. (D) Quantification of human DNA via Alu PCR in quadriceps and gastrocnemius muscles at various time-points following HLI surgery/cell injection. Vehicle tissue was collected at Day 64 only. Spike-in controls used the same VPC lot diluted in naïve mouse quadriceps tissue. *p<0.05 vs. vehicle (muscle-group specific) of log-transformed data; One-way ANOVA followed by Dunnett’s test. Data is mean ± SEM, n=3 independent animals / group (E,F) Representative immunofluorescent images of VPC engraftment in Balb/c nude quadriceps muscle Day 64 after HLI surgery / cell injection. (E) Whole section imaging of entire quadriceps muscle with Ku80+ regions outlined in white (F) 20x image demonstrating nuclear localization of human-specific marker, Ku80. Scale bar = 50µm.

Using primate-specific Alu detection by PCR(85,86), we were able to confirm presence of human cells in the mouse quadriceps muscle up to Day 3 and the mouse gastrocnemius muscle up to Day 64. These studies suggest that while VPCs can persist in the hypoxic gastrocnemius, the total number is low with only ∼10,000 cells out of the 1 million injected after Day 7 as determined by PCR (Figure 4D). Again, examination of other organs confirmed no off-target migration of VPCs (Supplemental Figure 11). Despite the low number of persistent cells, we were able to visualize human cells by immunofluorescence using Ku80 as a human-specific marker up to Day 64 following HLI and cell injection in athymic Balb/c nude mice (Figure 4E-F). We did not observe any long-term persistence (i.e Day 64 after cell injection) of VPCs in immunocompetent db/db mice by either hAlu PCR or Ku80 immunofluorescence (Supplemental Figure 12). Together, these results indicate host revascularization and improved limb perfusion were able to occur with limited / transient donor cell engraftment, supporting a paracrine mechanism of action for VPC *in vivo*.

### VPC significantly improve limb perfusion and vascular integrity in therapeutic model in diabetic mice

To bridge preclinical and clinical relevance, we developed a therapeutic model of experimental CLTI in diabetic mice and attempted to incorporate endpoints (LSCI perfusion imaging(87), tissue oxygenation(87), functional mobility assessments(88)) aligned with CLTI therapeutic trials as much as reasonably possible in a mouse. To this end, EXT-VPC were injected 21 days after HLI surgery to more closely mimic the chronic nature and therapeutic window of CLTI (Figure 5A, white arrow). Of note, we tested cells generated in scale-up formats (either 2CS or 5CS for Day 0-6 and HyperFLASK for Day 7-10) using GMP compatible materials that were cryopreserved in a, “clinic-ready” RICEP (reconstitute and inject cell product) format, in which cell vials are packaged at high density and can be immediately thawed with high viability and recovery, and directly diluted and injected without intermediate washing. Cells were injected into four sites along the length of the limb to more closely model the multiple injection sites seen in CLTI clinical trials.

**Figure 5:**
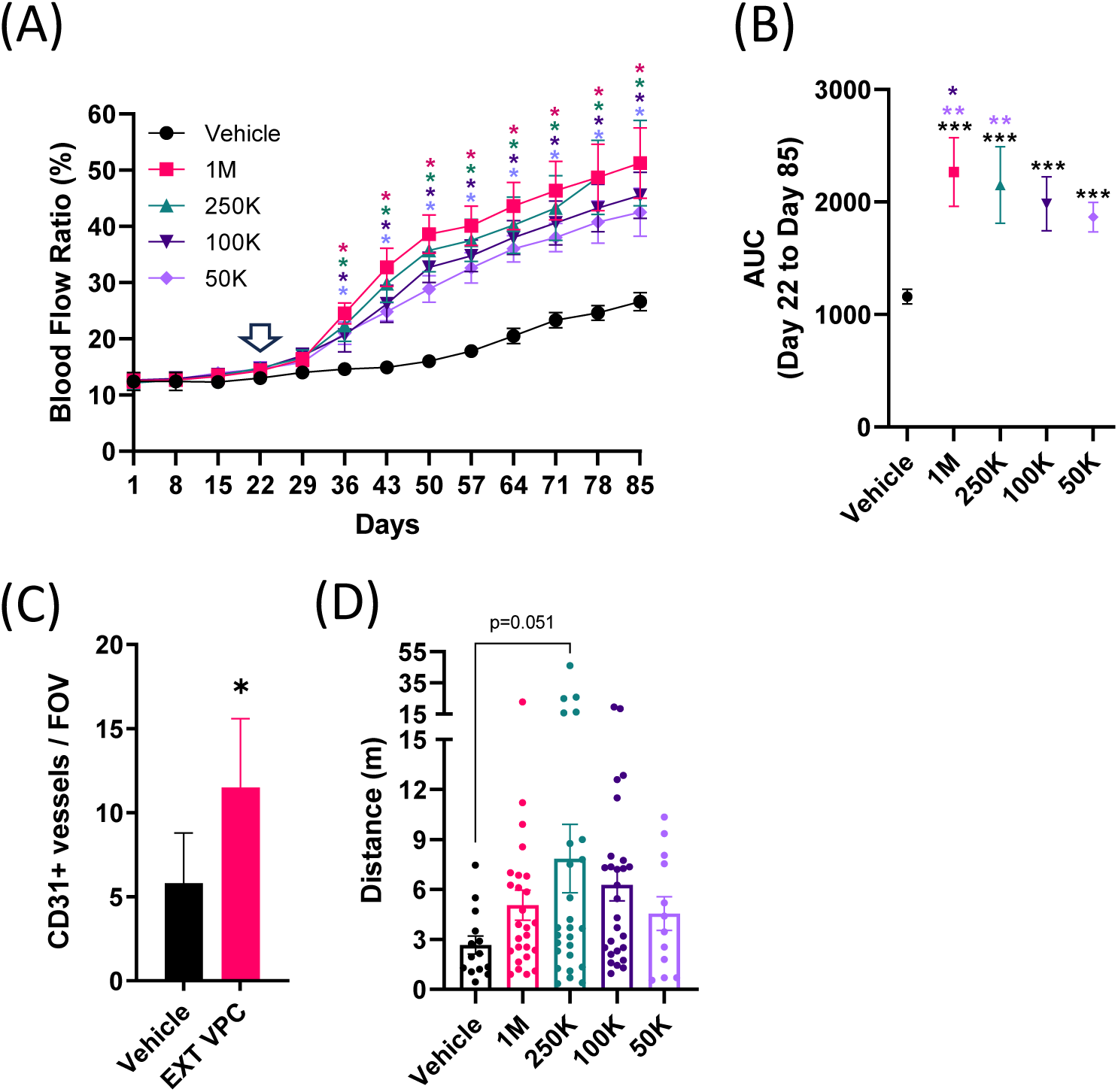
VPC significantly improve limb perfusion and vascular integrity in therapeutic model in db/db mice. (A) Laser Speckle Contrast Imaging (LSCI) of db/db mice in therapeutic model with treated with extended culture VPCs or vehicle across multiple injection sites. HLI was performed as before on Day 1 then cells were injected at 2 sites in the quadriceps and 2 sites in the gastrocnemius at Day 22 (indicated by white arrow). Data is mean ± SD, n = 14-27 per group, *p<0.05 vs. vehicle; Two-way ANOVA, Tukey post-hoc test. (B) LSCI area under the curve from Day 22 through Day 85 (course of cell treatment). Data is mean ± SEM, n = 14-27 per group, ***p<0.05 vs. vehicle; **p<0.05 vs. 50K dose;*p<0.05 vs 100K dose; One-way ANOVA followed by Tukey post-hoc test; (C) Immunohistochemisty analysis of CD31+ vessels in quadriceps muscle at Day 85. Data is mean ± SEM, n=6 mice/group, *p<0.05 vs. vehicle; Student’s t-test (D) Total average treadmill running distance at Day 85 (2 runs), Data is mean ± SEM; n=14-27, One-way ANOVA followed by Dunnett’s test

In this therapeutic model, VPC exhibited a dose-dependent sustained improvement in blood flow following HLI despite delaying cell injection until Day 22 post-HLI surgery. All VPC doses significantly (p<0.0001) improved blood flow recovery compared to vehicle control as early as 14 days after cell injection (Figure 5A). This improvement in blood flow over vehicle treated animals was sustained up to 12-weeks later (Day 85) across all doses (1.9X, 1.9X, 1.7X, 1.6X vs. vehicle for 1M, 250K, 100K, and 50K VPC doses respectively; p<0.0001) (Figure 5A). Using the area under the curve (AUC) of blood flow ratio over time as a cumulative comparison of the VPC dose response, we saw that there was no significant difference in blood flow recovery between 1M and 250K doses (p=0.51) whereas both were significantly greater than the 50K dose (p<0.05) (Figure 5B). Furthermore, there was a higher CD31^+^ vessel density in VPC treated animals compared to controls at Day 85, confirming the cytoprotective and regenerative properties of VPC treatment (Figure 5C). This was further supported by microCT analysis showing increased vessel density in VPC treated animals (data not shown). Additionally, there was a modest trend in exercise capacity at Day 85 with VPC treatment, particularly for mice treated with 250K VPC (p=0.051) compared to vehicle (Figure 5D). Hind paw oxygen saturation (StO_2_) was significantly impaired in ischemic limb compared to non-operated contralateral controls in vehicle treated animals from Day 22 through Day 85 post-HLI surgery, although a trend toward higher StO_2_ was noted, particularly with the 250K VPC dose at Day 85, VPC treatment did not yield a significant improvement (Supplemental Figure 13). In this study, we investigated significantly later time-point than had previously been reported following experimental HLI in the mouse(89,90). Consistent with our previous findings, VPC were well tolerated, and we did not observe any significant alterations compared to vehicle controls in terms of body weight, animal welfare, clinical hematology or clinical chemistry in this therapeutic model (Supplemental Figure 14).

## Conclusions

This study demonstrates that universal hiPSC-derived VPC, produced via a scalable 10-day differentiation protocol (EXT-VPC), achieve sustained limb reperfusion and tissue repair in diabetic CLTI and Balb/C CLTI models through multimodal mechanisms exerted by paracrine factors locally produced by VPC in the injection sites. By combining HLA engineering with a chemically defined GMP-compatible manufacturing process, we generated VPC that evade innate and adaptive immune responses while secreting several pro-angiogenic (e.g. HGF, ANGPT2, ANGPTL4, EGF, CCL2, angiogenin) and cytoprotective (e.g. HGF, EGF, FGF2) factors critical for inducing host tissue neovascularization and stabilization of new and pre-existing host vessels. Notably, the absence of VEGF-A secretion, a hallmark of inflammatory angiogenesis and lack of expression for pro-inflammatory cytokines IL6, IL8, or IP-10/CXCL10 may explain the observed increased aSMA^+^CD31^+^ stable vessel formation in VPC-injected limb tissues(91). Similar to endogenous endothelial cells, VPC may exhibit a context-dependent signaling profile and function. For example, in the hyperproliferative environment of pulmonary vasculature, VPC reduced pulmonary arterial pressure indicative of a reduction in excessive vascular growth and increased vessel stabilization (data not shown) but in the context of vascular insufficiency as in the CLTI model, VPC are able to promote revascularization

Our findings advance prior work in three key areas. First, we provide a detailed characterization of VPC phenotypic and functional attributes supporting their multifactorial therapeutic potential, including angiogenic, cytoprotective, and immunomodulatory properties relevant to the complex pathobiology of CLTI. Importantly, the EXT-VPC process enhances product safety by reducing residual pluripotent or partially differentiated cells while preserving angiogenic, cytoprotective, and immunomodulatory properties. Moreover, unlike autologous cell therapies requiring patient-specific cell harvest/expansion, these universal VPC function as an off-the-shelf product, retaining potency after cryopreservation and scalable expansion in a GMP-compatible manufacturing process (94). Second, we demonstrate that the universal donor cell (UDC) edits do not impair VPC function, enabling immune-evasive behavior without altering their core angiogenic, cytoprotective, and immunomodulatory properties. While immune-evasive edits may offer some protection from innate immune recognition, their impact is expected to be less pronounced than in engraftable cell therapies, as VPC mediate benefit primarily through paracrine mechanism rather than durable engraftment. Additionally, the incorporation of a thymidine kinase (TK) safety switch further strengthens the risk-mitigation profile. Third, (95) we enhanced preclinical transability by incorporating diabetic models, delayed-treatment paradigms, and clinically aligned functional endpoints, thereby addressing known limitations of standard hindlimb ischemia models.

While there is no preclinical model that fully recapitulates all aspects of the various etiologies and manifestations of CLTI(92), we attempted to overcome some of the inherent weaknesses of the typical hindlimb ischemia model. To this end, we utilized diabetic, db/db mice, which incorporates a major co-morbidity of the CLTI population and are known to have a prolonged reduction in limb perfusion with limited spontaneous recovery due to vascular dysfunction(93). We sought to move beyond the prophylactic model in which most conventional HLI models administer drug candidates immediately or 1-day after the HLI operation(94). We therefore developed a therapeutic model in which we administered our cell therapy 21-days post-HLI operation. At this timepoint, there is still severe limb ischemia with reduced perfusion that exhibited limited additional spontaneous recovery, as well as on-going myocyte damage(23,93). This underscores the relevance to clinical CLTI, where patients often present with chronic ischemia and established tissue damage. Finally, conventional HLI models tend to limit functional analysis to semi-quantitative scoring and histology, neither of which is an objective endpoint directly relating to clinical CLTI studies. We sought to include oxygen saturation as an additional functional endpoint to perfusion to these traditional pre-clinical measures. While further optimization of the technique may enhance outcomes, we observed a trend toward improved functional capacity via treadmill testing after EXT-VPC treatment. It is notable that this trend occurred with the 250K cell dose, suggesting a potential non-linear dose-response relationship. Such a pattern could reflect complex biological interactions, including optimal cell-to-tissue integration thresholds, paracrine factor signaling dynamics, or microenvironmental constraints at higher cell densities, warranting deeper investigation in future studies. Nonetheless, clinical translation may be aided by more routine inclusion of clinically meaningful, functional readouts into CLTI preclinical models

Our data showing the therapeutic efficacy of VPC in diabetic animals with a delayed-treatment model, underscores their relevance to clinical CLTI. The ∼2-fold improvement in perfusion ratios and ∼50% reduction in fibroadipose degeneration correlate with histologic evidence of mature or stable neovessels and myofiber regeneration, suggesting VPC reduces both microvascular dropout and myocyte apoptosis. RNA-seq data from treated vs untreated animals further suggest VPC-mediated suppression of IFNγ and TNFα pathways, which drive excessive M1 polarization resulting in chronic inflammation and tissue destruction in chronic ischemia(95) experienced by CLTI patients. These mechanistic insights align with recent preclinical advancements in testing cell therapies in hindlimb ischemia models that better replicate chronic and severe ischemia(96). Similarly, recent reports suggest that genetically engineered iPSC-derived hypoimmunogenic “universal” endothelial cells(97), and M2-like macrophages(98) improve perfusion and vascularization in rodent models. Novel delivery approaches, including mRNA-LNPs encoding ETV2(99) and hydrogel scaffolds co-delivering MSCs and mast cells(100), further improve therapeutic outcomes in preclinical studies. These innovations collectively improve disease modeling, delivery, and efficacy assessment of vascular regenerative therapies. While independent validation remains essential, these findings mark an important step toward optimizing cell-based therapies and accelerating clinical translation for ischemic vascular diseases, particularly in high-risk, surgical and endovascular treatment-ineligible CLTI populations.

Some of the limitations of this study include the inherent differences between murine and human immune systems and differences in size/surface area of limbs or differences in CLTI wound healing properties, necessitating future studies in humanized models or large animals with diabetic wounds or atherosclerotic comorbidities. Additionally, it is unknown how dosing translates from rodent to human. While preclinical studies often report efficacy in rodent hindlimb ischemia models, human trials have struggled to replicate these results which may be in part due to the suboptimal delivery of therapeutic, including route of administration, ensuring sufficient coverage to the affected area, and inadequate dose adjustments for species-specific biological responses. To this end, several porcine models that more closely mirror the anatomy and scale of humans as well as more readily enable the incorporation of clinical readouts including TcPO_2_, MRI angiography, duplex ultrasound, and treadmill walking have been developed(101–107). Finally, while VPC persisted in hypoxic niches for 64 days, their gradual decline suggests paracrine effects, rather than long-term engraftment, dominate therapeutic outcomes. Establishing relationships between cell persistence, therapeutic factor pharmacokinetics, and efficacy may be beneficial for both future pre-clinical and clinical studies.

In conclusion, our platform bridges critical gaps in cell therapy-based regenerative medicine for CLTI, offering the opportunity for developing a scalable, safe, and mechanism-driven alternative to palliative care currently available for CLTI. By prioritizing immune stealth, functional potency, and clinical trial-relevant endpoints, this work lays the foundation for GMP-compatible manufacturing of universal VPC in a clinical scale for Phase I/II trials evaluating VPC for “poor-option” CLTI patients. Future efforts may explore therapeutic cell engineering or combinatorial approaches, such as engineered cells for CLTI specific therapeutic cargo delivery or co-administering VPC with biomaterials, to further enhance therapeutic durability.

## List of abbreviations

CLTI: Chronic limb threatening ischemia
hiPSC: human induced pluripotent stem cell
VPC: vascular progenitor cell
GMP: good manufacturing process
BM-MNC: bone marrow mononuclear cell
PB-MNC: peripheral blood mononuclear cell
MSC: mesenchymal stromal cell
ECFC: endothelial colony forming cell
TK: thymidine kinase
scRNA-seq: single-cell RNA sequencing
EXT-VPC: extended culture vascular progenitor cell
EC: endothelial cell
HUVEC: human umbilical vein endothelial cell
HLI: hindlimb ischemia
αSMA: smooth muscle alpha actin
RICEP: reconstitute and inject cell product

## Declarations

### Ethics approval and consent to participate

All animal studies were approved by internal review boards at either Invivotek (Hamilton, NJ, USA) or Pharmaseed (Ness Ziona, Israel) and conformed to the *Guide for the Care and Use of Laboratory Animals* (US National Institutes of Health) and Association for Assessment and Accreditation of Laboratory Animal Care (AAALAC).

### Consent for publication

Not applicable

### Availability of data and materials

The datasets generated and/or analyzed during the current study are not publicly available due to potential commercial sensitivity. However, the data are available from the corresponding author upon reasonable request and subject to a Non-Disclosure Agreement (NDA).

### Competing interests

All authors are current or former employees of Astellas Pharma.

## Funding

All research was sponsored by Astellas Pharma, which may develop and/or license products using the study’s findings

## Authors’ contributions

JH, PL, NP, EK conceived of the study and article. JH and NP wrote the manuscript and prepared the figures. JH, WS, HC, MSK, BS, SC, AA, RH, and SS acquired study data. JH, WS, PL, HC, MSK, DP, BS, SC, AA, RH, SS and NP analyzed the data. All authors contributed to specific experimental design, the interpretation of the experimental data, and reviewed and approved the final manuscript.

## Acknowledgements

We thank Pharmaseed Ltd for their support in executing the HLI efficacy studies and Invivotek for their support in executing the IVIS biodistribution study and 9-month non-GLP safety study. We would also like to thank Travis Wallace for technical / cell culture support and Akina Hoshino, Lisa Petek, and their teams for generating and providing the HLA-edited clones used in this manuscript.

## Notes

### Competing Interest Statement

The authors have declared no competing interest.

