## Supplementary material for "Scalable Generation of Universal hiPSC-Derived Vascular Progenitor Cells for Safe and Sustained Revascularization in Chronic Limb-Threatening Ischemia": CLTI VPC paper_supplemental figures.pdf

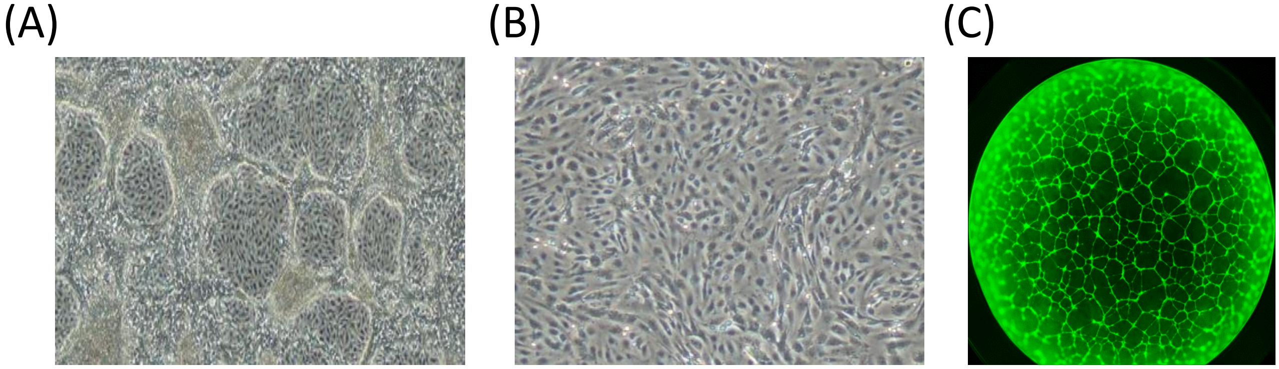


**Supplemental Figure 1: VPC and EXT VPC morphology.**
(A,B) Phase contrast images of (A) VPC and (B) EXT VPC immediately before harvest in a T175 flask . (C) Whole well image of tube formation of VPC when plated on Matrigel and stained with AcLDL-AlexaFluor488

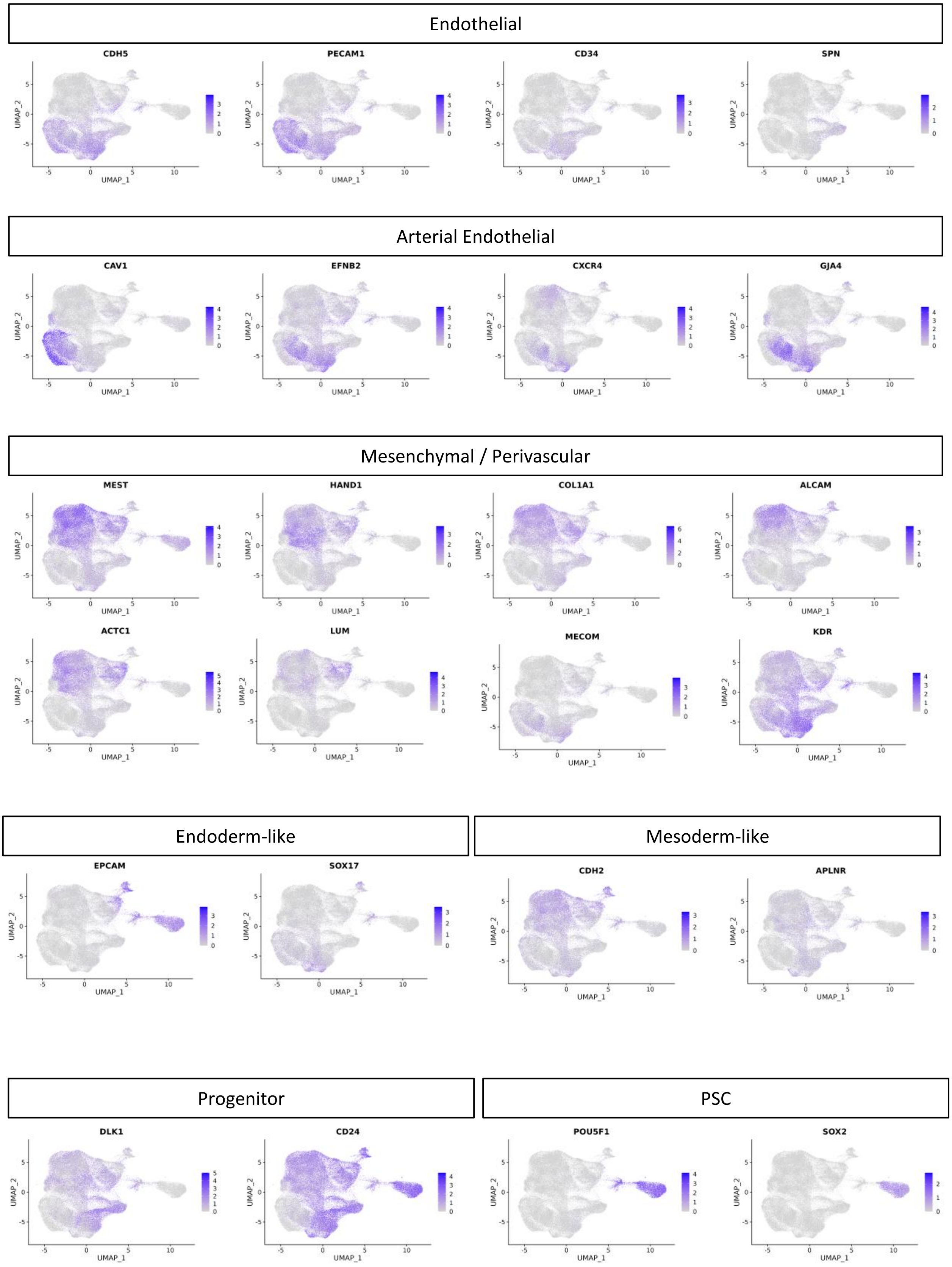


**Supplemental Figure 2: VPC subpopulation marker scRNA-seq expression.**
UMAP plots of various markers using scRNA-seq reference mapping from Figure 1C of 11 total VPC lots, including compliant, scale-up, and extended culture VPC as well as non-VPC reference cell lines (PSC and HUVEC) (>100K cells total) labeled by expression level

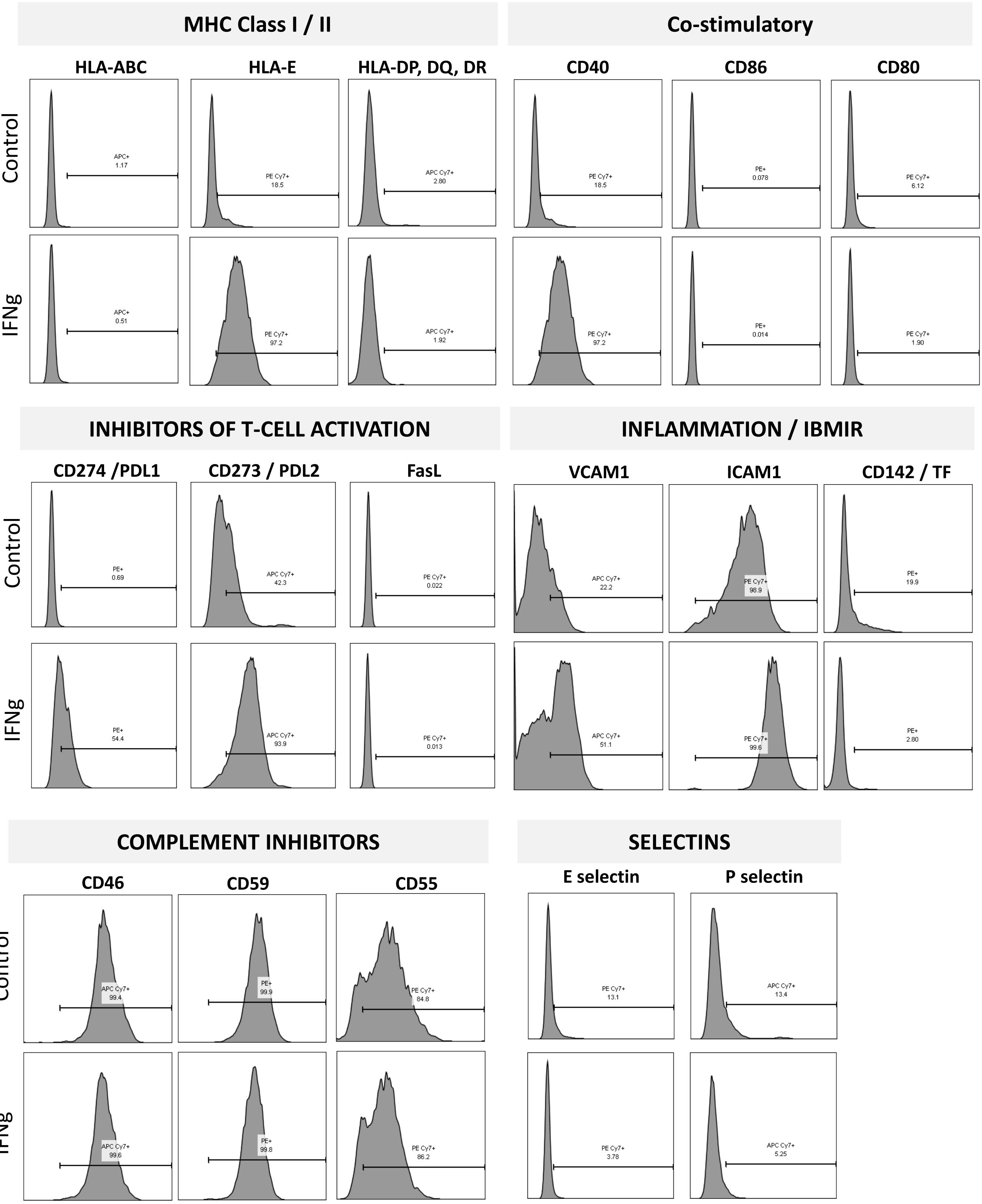


**Supplemental Figure 3: Immunophenotyping of VPC by flow cytometry with and without IFN stimulation.** Histograms of various immune / inflammation markers of VPC cultured for 2 days ± 8ng/mL IFNγ as determined by flow cytometry

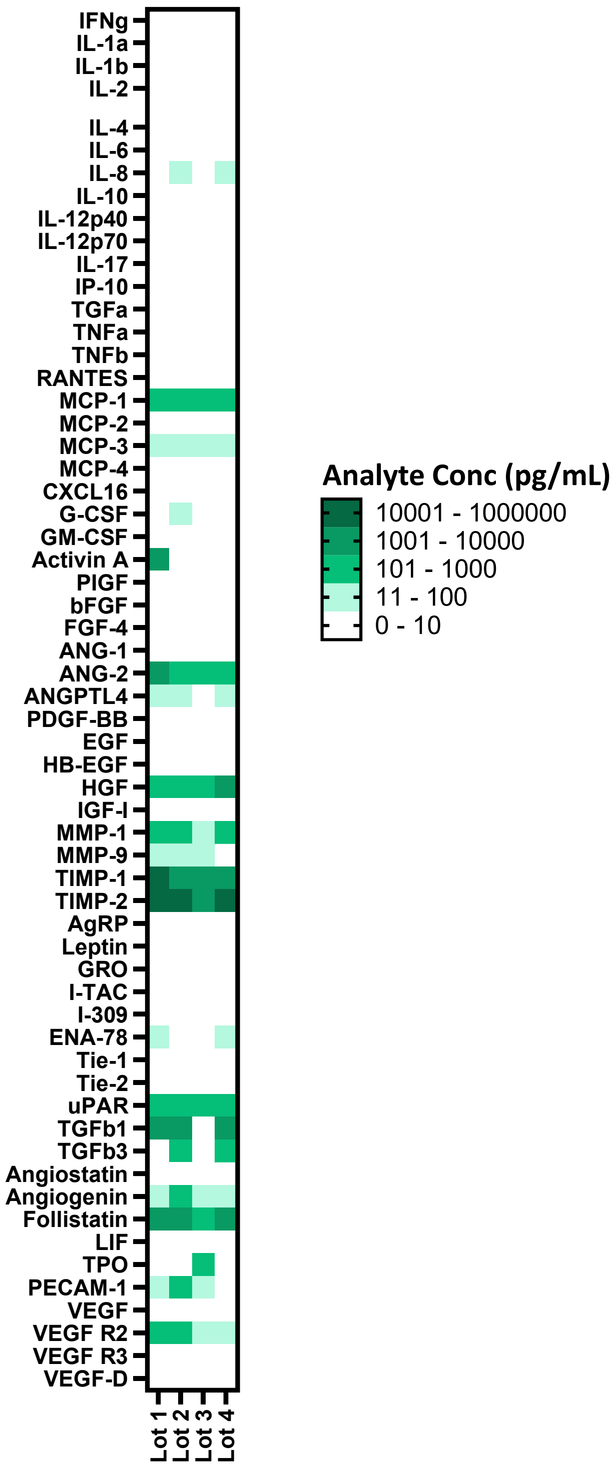


**Supplemental Figure 4: Angiogenesis array of secreted factors in VPC conditioned media.**
Heatmap of angiogenesis related protein expression from conditioned media generated from 4 independent VPC lots.

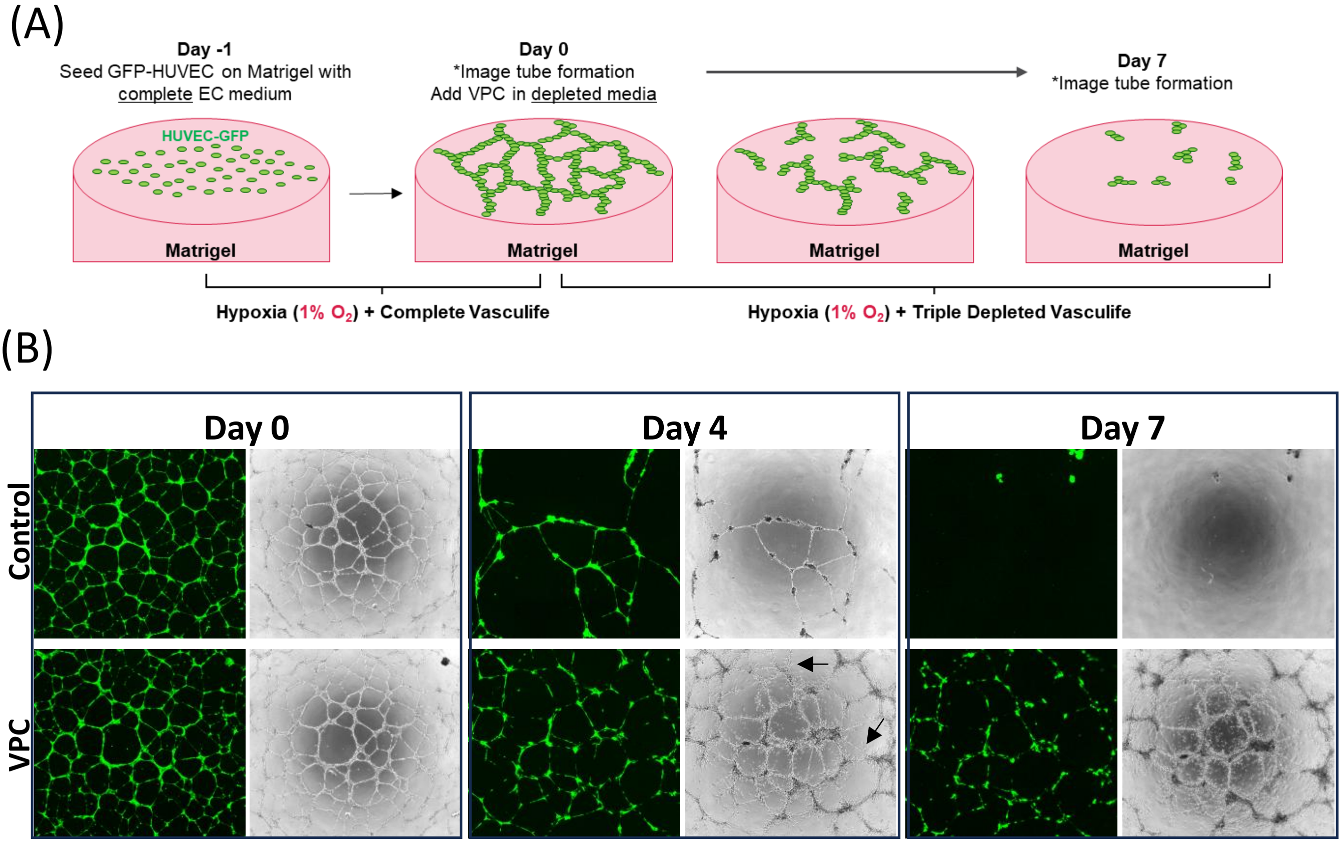


**Supplemental Figure 5: Schematic and time-course of tube stabilization assay.**
(A) Schematic of tube stabilization assay. GFP-HUVEC are plated on Matrigel in growth media and allowed to self-assemble into a vascular tube network for 24-hours in hypoxic conditions (1% O2). VPC are then seeded in “triple depleted Vasculife” media (i.e. Vasculife media without VEGF, FGF2, or FBS) and cultured in 1% O2 for an additional 7 days with no media changes. (B) Epifluorescence imaging of GFP-HUVEC and total tube network (phase contrast) at day 0 , 4, and 7 in control (no cells, triple depleted Vasculife only) and VPC. Arrows indicate examples of nascent “sprouting” by VPC

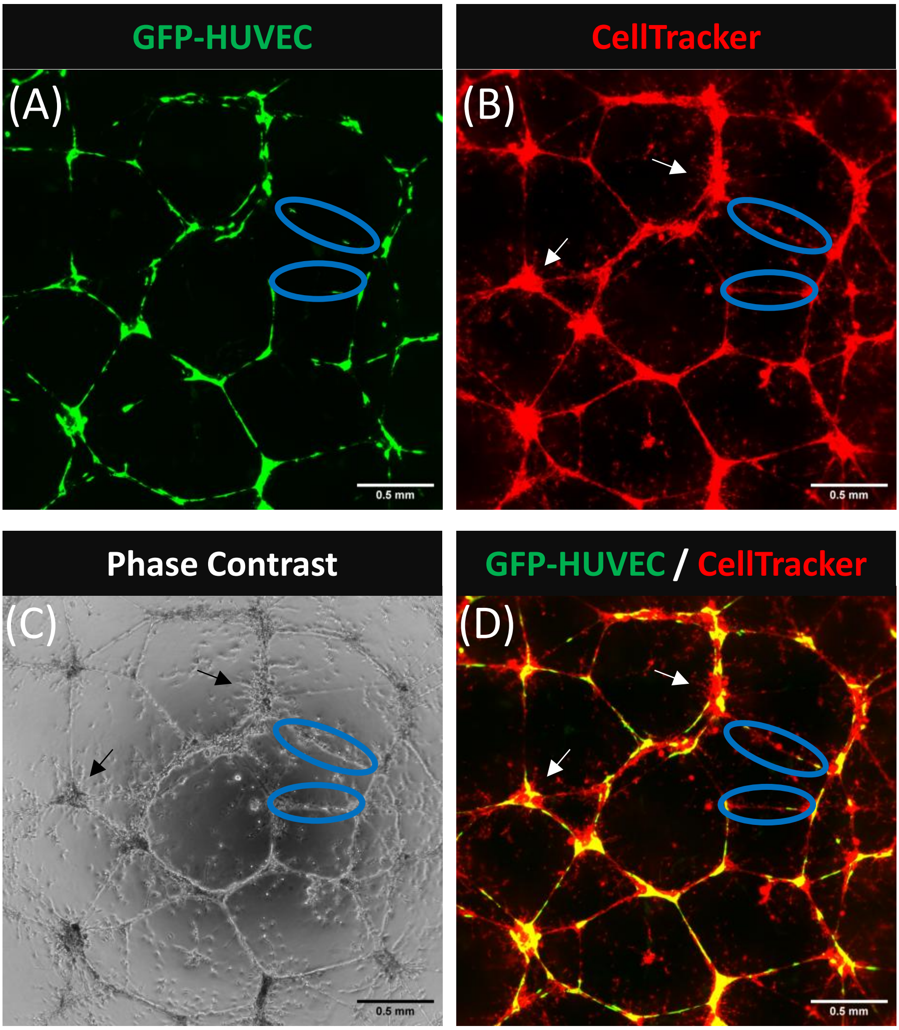


**Supplemental Figure 6: Phenotype of VPC in tube stabilization assay.**
Representative images from GFP-HUVEC tube stabilization assay with VPC treatment at D7. (A) GFP-HUVEC (B) CellTracker labeled representing all cells (HUVEC + VPC) (C) Phase contrast representing all cells (HUVEC + VPC) and (D) Merged image of GFP-HUVEC and CellTracker labeling. Arrows indicate examples of nascent "sprouting" presumably by VPC. Blue ovals indicate examples of VPC only segments where VPC has replaced previous HUVEC tubes. Scale bar is 0.5mm in all images.

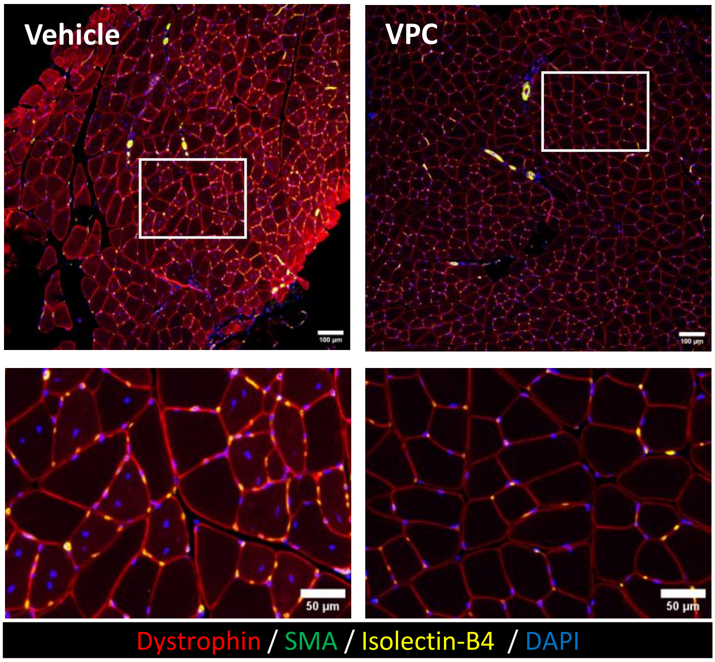


**Supplemental Figure 7: VPC-treated muscles display mature muscle fiber phenotype whereas vehicle-treated animals still have regions of regenerating myofibers in db/db mice**

Representative images from vehicle and VPC treated quadriceps muscles from db/db mice at day 64 post-HLI surgery / treatment injection. Top row indicates representative 10X image with scale bar = 100 µm. Bottom row is inset from top image where scale bar = 50 µm. Dystrophin stains myofiber border, SMA = smooth muscle α-actin for vascular smooth muscle cells , isolectin-b4 stains mouse endothelial cells, and DAPI for nuclei.

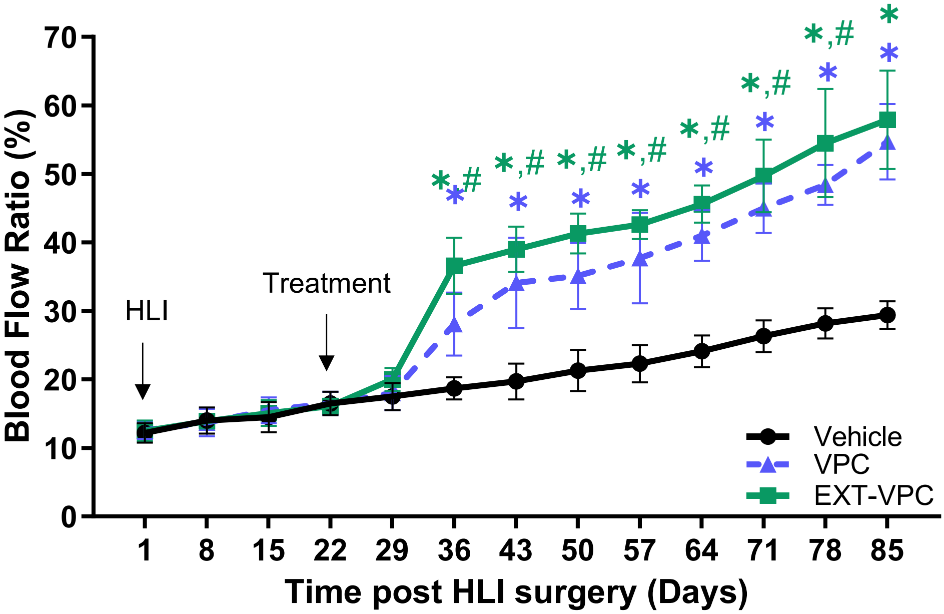


**Supplemental Figure 8: EXT-VPC exhibits increased blood flow recovery in therapeutic diabetic HLI model**

Db/db mice underwent HLI sugery at Day 1. Mice were injected i.m. in 2 sites in quadriceps muscle with either 1 million VPC or EX-VPC or vehicle control (PlasmalLyte A) on Day 22. Blood flow was monitored by LSCI. All doses divided between 2 sites. N>10 for each group; Data is mean ± SEM; *p<0.05 vs Vehicle; #p<0.05 vs. VPC. Two-way ANOVA+Tukey's test;

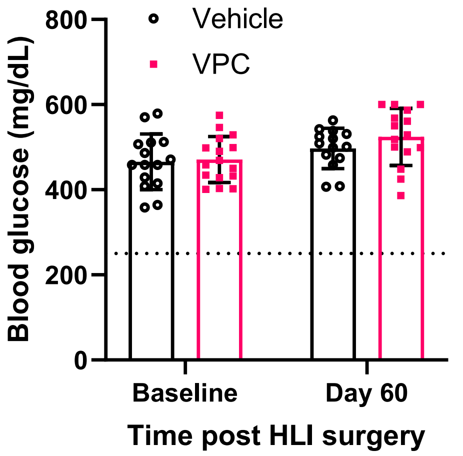


**Supplemental Figure 9: Non-fasting blood glucose in db/db mice**
Non-fasting blood glucose in db/db mice in vehicle and VPC treated animals at various timepoints following HLI surgery. 250mg/dL was used as the threshold for an animal to be deemed “diabetic”. Data is mean ± SD.

(A)

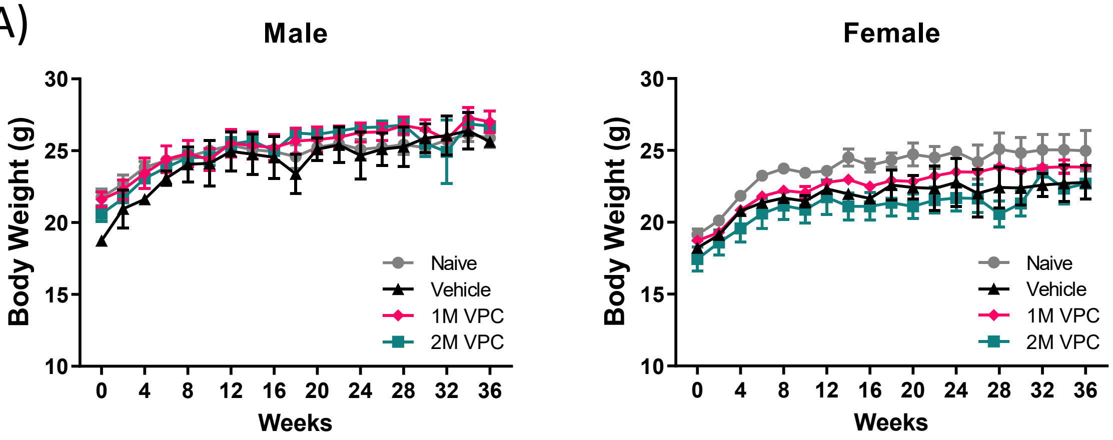


(B)

| | | Naïve | | | Vehicle | | | VPC 1M dose | | | VPC 2M dose | | |
|---|---|---|---|---|---|---|---|---|---|---|---|---|---|
| | | *n* | *Mean* | *SEM* | *n* | *Mean* | *SEM* | *n* | *Mean* | *SEM* | *n* | *Mean* | *SEM* |
| Albumin | (g/dL) | 3 | 2.83 | 0.12 | 4 | 2.75 | 0.16 | 7 | 2.73 | 0.08 | 6 | 2.20 | 0.42 |
| Total Prot | (g/dL) | 4 | 4.20 | 0.75 | 2 | 4.35 | 0.65 | 7 | 4.70 | 0.08 | 8 | 5.43 | 0.26 |
| Alk Phos | (u/L) | 4 | 57.50 | 4.01 | 2 | 54.00 | 5.00 | 7 | 54.86 | 2.01 | 9 | 63.00 | 3.53 |
| Alt | (u/L) | 3 | 163.33 | 16.33 | 3 | 70.67 | 63.17 | 8 | 184.75 | 79.48 | 9 | 157.78 | 46.92 |
| Ast | (u/L) | 4 | 213.50 | 44.13 | 3 | 175.00 | 76.14 | 9 | 208.22 | 47.65 | 9 | 201.89 | 48.32 |
| CK | (u/L) | 4 | 1066.75 | 702.81 | 3 | 1631.33 | 794.61 | 6 | 1946.17 | 642.23 | 7 | 925.00 | 525.19 |
| Bun | (mg/dL) | 2 | 33.00 | 11.00 | 2 | 20.50 | 1.50 | 9 | 21.44 | 0.94 | 8 | 23.63 | 1.28 |
| Calcium | (mg/dL) | 4 | 9.05 | 0.16 | 3 | 6.30 | 2.80 | 9 | 7.46 | 0.93 | 8 | 9.54 | 0.07 |
| Tbili | (mg/dL) | 4 | 0.38 | 0.05 | 2 | 0.40 | 0.10 | 7 | 0.37 | 0.05 | 9 | 0.37 | 0.05 |
| Creatinine | (mg/dL) | 2 | 0.45 | 0.12 | 0 | nd | nd | 2 | 0.36 | 0.00 | 1 | 0.45 | na |
| Trig | (mg/dL) | 4 | 93.50 | 10.40 | 4 | 89.50 | 19.59 | 8 | 101.13 | 7.39 | 9 | 103.00 | 3.99 |
| Phosphorus | (mg/dL) | 4 | 7.43 | 0.75 | 3 | 7.00 | 0.74 | 8 | 6.01 | 0.79 | 8 | 7.81 | 0.37 |
| Na | (mmol/L) | 4 | 147.00 | 0.69 | 4 | 147.40 | 0.55 | 9 | 152.88 | 1.89 | 9 | 150.98 | 1.51 |
| K | (mmol/L) | 4 | 5.66 | 0.19 | 4 | 6.17 | 0.46 | 9 | 6.21 | 0.27 | 9 | 7.15# | 0.34 |
| Cl | (mmol/L) | 4 | 115.18 | 1.17 | 4 | 116.33 | 0.29 | 9 | 121.59# | 1.35 | 9 | 119.01 | 0.58 |
| Globulin | (g/dL) | 3 | 2.10 | 0.12 | 2 | 1.50 | 0.60 | 7 | 1.97 | 0.09 | 6 | 3.15 | 0.52 |
| A/G Ratio (g/dL) | | 3 | 1.35 | 0.02 | 2 | 2.25 | 0.87 | 7 | 1.40 | 0.09 | 6 | 0.87 | 0.23 |
| Cholestrol | (mg/dL) | 4 | 76.50 | 6.95 | 4 | 59.00 | 13.67 | 9 | 73.89 | 5.04 | 9 | 79.67 | 5.08 |
| Glucose | (mg/dL) | 3 | 234.33 | 58.77 | 4 | 172.00 | 30.63 | 9 | 225.44 | 8.14 | 8 | 224.25 | 8.74 |

(C)

| | Naïve (n=4) | Vehicle (n=4) | VPC - 1M (n=9) | VPC-2M (n=9) |
|---|---|---|---|---|
| ***WBC ($x10^3$)*** | 5.45 ± 0.99 | 2.94 ± 0.50 | 5.74 ± 0.73 | 5.55 ± 0.51 |
| ***NE ($x10^3$)*** | 2.07 ± 0.30 | 0.12 ± 0.02 | 2.94 ± 0.59 | 2.00 ± 0.26 |
| ***LY ($x10^3$)*** | 3.24 ± 0.83 | 0.02 ± 0.01 | 2.95 ± 0.32 | 3.30 ± 0.36 |
| ***MO ($x10^3$)*** | 0.13 ± 0.01 | 0.01 ± 0.00 | 0.19 ± 0.03 | 0.22 ± 0.03 |
| ***EO ($x10^3$)*** | 0.02 ± 0.00 | 11.59 ± 0.35 | 0.03 ± 0.01 | 0.02 ± 0.01 |
| ***BA ($x10^3$)*** | 0.01 ± 0.00 | 673.75 ± 51.11 | 0.01 ± 0.00 | 0.01 ± 0.00 |
| ***RBC ($x10^6$)*** | 11.01 ± 0.57 | 11.59 ± 0.35 | 11.52 ± 0.22 | 11.16 ± 0.48 |
| ***Platelets ($x10^3$)*** | 606.75 ± 41.45 | 673.75 ± 51.11 | 589.20 ± 52.51 | 636.56 ± 41.80 |

**Supplemental Figure 10: VPC were well tolerated in 9-month non-GLP safety study**
(A) Weekly body weight (B) blood clinical chemistry and (C) clinical hematology of naïve athymic Balb/c nude mice (2 male/2 female), vehicle treated (2 male / 2 female) , and VPC treated with either 1 million VPC (1M VPC, 4 male / 5 female) or 2 million VPC (2M VPC, 4 male/ 4 female) up to 9-months after injection. Data is mean ± SEM. Either 2-way ANOVA (body weight) or one-way ANOVA for each analyte (clinical chemistry and hematology) following by Tukey post-hoc tests.*p<0.05 vs vehicle, #p<0.05 vs. naïve

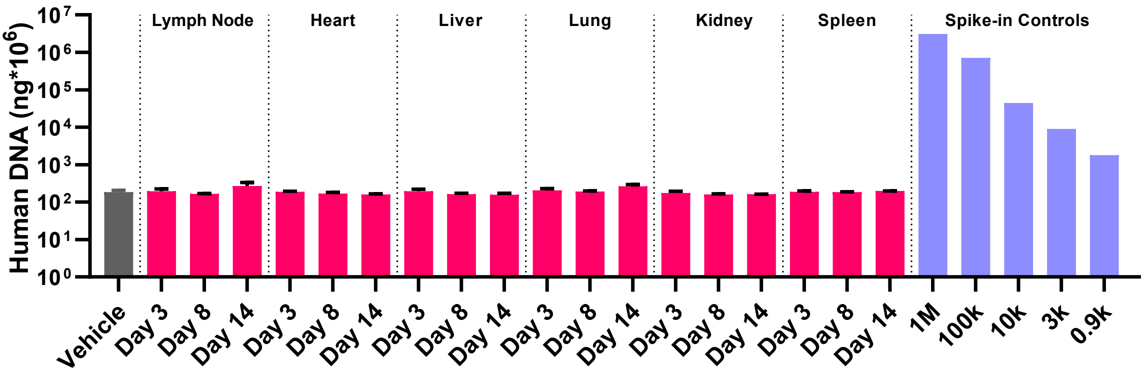


**Supplemental Figure 11: No off-target detection of human cells in non-injected organs by hAlu PCR**
Calculated human DNA based on hAlu PCR spike-in controls in athymic Balb/c nude mice following injection of 2 million VPC into quadriceps and gastrocnemius muscles 24-hours following HLI surgery. Data is mean ± SEM; n=3 animals per organ per time-point performed in technical triplicate

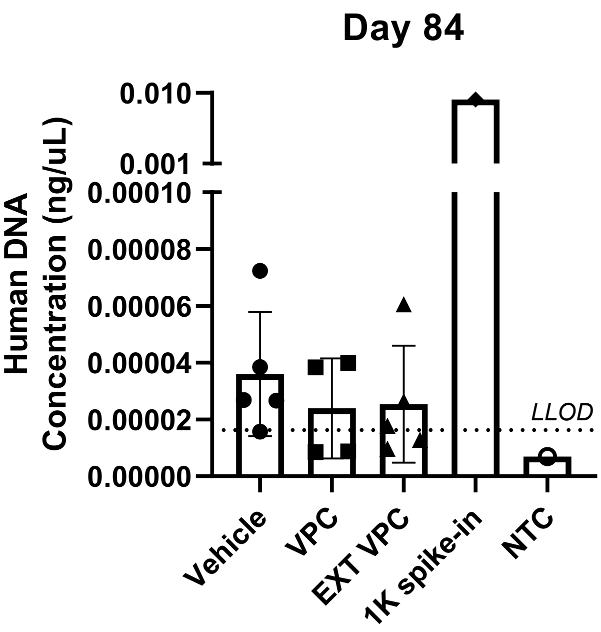


**Supplemental Figure 12: No long-term detection of VPC or EXT VPC in db/db mice**

Calculated human DNA based on hAlu PCR in therapeutic db/db model 9-weeks (Day 85) after injection of 1 million VPC into quadriceps muscles . Data is mean ± SEM; n=4 animals performed in technical triplicate

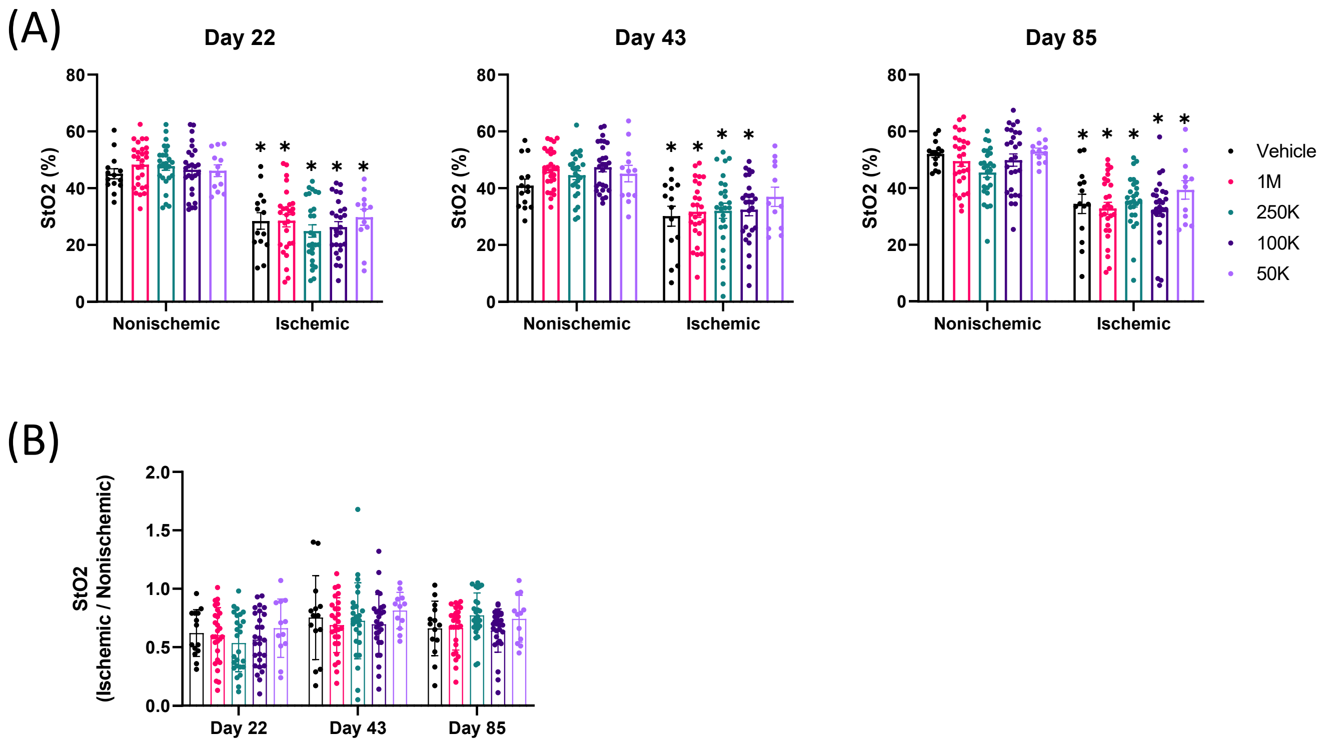


**Supplemental Figure 13: No significant improvement in StO2 with EXT VPC treatment in therapeutic model in db/db mice**

(A) St02 (%) in hindpaw of both non-ischemic (non-operated) and ischemic / HLI-operated limb in db/db mice at Day 22 (day of treatment), Day 43, and Day 85 after HLI-surgery in vehicle and EXT VPC treated mice. (B) Ratio of % StO2 in ischemic paw to non-ischemic paw in same animal over time. Data is mean ± SEM; *$p<0.05$ vs. respective non-ischemic paw

(A)

| Clinical Chemistry | | | |
|---|---|---|---|
| | **Vehicle (n=6)** | **EXT VPC (n=6)** | ***p-value*** |
| Creat (mg/dl) | 0.3 ± 0.1 | <0.17 | NA |
| Calc (mg/dl) | 10.9 ± 0.4 | 10.5 ± 0.8 | 0.273 |
| Phos (mg/dl) | 10.4 ± 1.8 | 9.2 ± 1.5 | 0.248 |
| Gluc (mg/dl) | 801.3 ± 92.9 | 668.3 ± 90.0 | 0.030 |
| Urea (mg/dl) | 41.1 ± 4.7 | 36.0 ± 4.5 | 0.087 |
| Chol (mg/dl) | 123.2 ± 25.3 | 101.7 ± 25.6 | 0.174 |
| TP (g/dl) | 6.5 ± 0.4 | 5.9 ± 0.7 | 0.102 |
| Alb (g/dl) | 4.7 ± 0.5 | 3.4 ± 0.5 | 0.001 |
| Glob (g/dl) | 1.8 ± 0.3 | 2.5 ± 0.9 | 0.098 |
| T. Bil (mg/dl) | 0.0 ± 0.0 | 0.0 ± 0.0 | 0.511 |
| Alk Phos (IU/L) | 186.8 ± 20.1 | 111.0 ± 41.0 | 0.002 |
| SGOT (IU/L) | 181.7 ± 77.6 | 93.3 ± 55.7 | 0.047 |
| SGPT (IU/L) | 181.0 ± 35.8 | 108.7 ± 75.1 | 0.059 |
| Na (mmol/L) | 157.2 ± 4.2 | 152.0 ± 6.7 | 0.141 |
| K (mmol/L) | 6.3 ± 1.4 | 6.2 ± 0.3 | 0.911 |
| Cl (mmol/L) | 102.3 ± 3.1 | 95.3 ± 3.7 | 0.006 |

(B)

| Clinical Hematology | | | |
|---|---|---|---|
| | **Vehicle (n=6)** | **EXT VPC (n=6)** | ***p-value*** |
| WBC (10*3/µl) | 2.5 ± 1.4 | 5.3 ± 3.3 | 0.095 |
| RBC (10*6/µl) | 9.7 ± 2.0 | 10.2 ± 0.9 | 0.596 |
| HGB (g/dl) | 15.1 ± 2.6 | 14.8 ± 1.9 | 0.840 |
| HCT (%) | 45.4 ± 8.0 | 46.7 ± 4.7 | 0.738 |
| MCV (FL) | 46.9 ± 1.8 | 45.7 ± 1.9 | 0.299 |
| MCH (pg) | 15.6 ± 0.9 | 14.4 ± 0.8 | 0.043 |
| MCHC g/dl) | 33.2 ± 1.7 | 31.6 ± 1.0 | 0.063 |
| Neut (%) | 46.5 ± 25.2 | 38.1 ± 11.8 | 0.517 |
| Bands (%) | 0.0 ± 0.0 | 0.0 ± 0.0 | NA |
| Lymph (%) | 42.1 ± 24.0 | 46.4 ± 16.0 | 0.772 |
| Mono (%) | 7.0 ± 2.8 | 12.4 ± 6.2 | 0.295 |
| Eos (%) | 3.9 ± 1.7 | 2.4 ± 1.7 | 0.329 |
| Baso (%) | 0.6 ± 0.2 | 0.3 ± 0.2 | 0.248 |
| Plate (10*3/µl) | 528.2 ± 219.5 | 1,314.3 ± 277.8 | 0.0003 |
| Neutr Abs (10*3/µl) | 1.5 ± 0.5 | 2.3 ± 2.1 | 0.645 |
| Bands Abs (10*3/µl) | 0.0 ± 0.0 | 0.0 ± 0.0 | NA |
| Lymph Abs (10*3/µl) | 1.6 ± 1.2 | 2.0 ± 0.5 | 0.412 |
| Mono Abs (10*3/µl) | 0.3 ± 0.1 | 0.8 ± 0.9 | 0.460 |
| Eos Abs (10*3/µl) | 0.1 ± 0.0 | 0.1 ± 0.0 | 0.472 |
| Basos Abs (10*3/µl) | 0.0 ± 0.0 | 0.0 ± 0.0 | 0.604 |

**Supplemental Figure 14: No adverse effects of EXT VPC injection in therapeutic model in db/db mice**
(A) Clinical chemistry and (B) clinical hematology of vehicle and EXT VPC treated db/db mice at Day 85 post-HLI sugery (9-weeks post-treatment). Data is mean ± SD; n=6 / group
