## Supplementary material for "Scalable Generation of Universal hiPSC-Derived Vascular Progenitor Cells for Safe and Sustained Revascularization in Chronic Limb-Threatening Ischemia": Supplemental File_Additional Materials and Methods.docx

***NK cytolysis assays***

NK cytolysis assays were performed as previously reported(1), with modifications. VPC were seeded at 3.5x10^4^ cells/cm^2^ in Vasculife VEGF Endothelial Medium (Lifeline Cell Technology, LL-003) ±40ng/mL interferon gamma (IFNγ) (Fisher, 300-02-100UG). IMR32 neuroblastoma cells (CCL-186, ATCC), served as the positive control and were cultured in IMR32 media [Eagle’s Minimal Essential Media (EMEM) + 10% FBS] ±40ng/mL IFNγ. Cells were cultured for three days without media change.

Three days after seeding, 10,000 cells/well (VPC) or 15,000 cells/well (IMR32) were seeded into fibronectin-coated plates xCelligence E-Plate View plates (Agilent). Plates were then analyzed by an xCelligence RTCA MP device (Agilent) with 1 sweep every 15 minutes for 24 hours. The next day, media in the xCelligence plates was replaced with NK cells in NK media, consisting of RPMI 1640 base media, 10% Human serum (heat-inactivated), 1x l-glutamine, 50 µM beta mercaptoethanol, 1mM sodium pyruvate, 1x NEAA, 400 U/mL IL-2 (Catalog Number: 11147528001, Sigma), and 25 mM HEPES, at the following NK:Target Cell ratios: 5:1, 1:1, 0.5:1, 0.1:1, and 0.1. Data collection then resumed in the xCelligence RTCA MP systems at a frequency of 1 sweep every 15 min for another 48hr. After completion, the percent target cell cytolysis was calculated using the xCelligence software.

***Proliferation of CD4+ and CD8+ T-cells***

Cryopreserved VPC were thawed and seeded at ~5000 cells / cm^2^ on fibronectin coated 10 cm dishes (Corning, 354451) in complete Vasculife VEGF Endothelial medium (Lifeline Cell Technology, LL-003). IFN-gamma (Fisher, 300-02-100UG) was added to cultures at 40 ng/ml. After 72 hours, mitomycin C (STEMCELL Technologies, 73274) was added to cultures at 10mM. After 3 hours, media was aspirated, cells were washed three times in PBS (Fisher, 14190-144), harvested with Accutase (STEMCELL Technologies, 07922), centrifuged, resuspended in RPMI (Fisher, 11875-093) with 10% FBS (Hyclone, SH30406.02) and seeded in 96-well flat bottom fibronectin plates (Corning, 354409) at a range of 60,000 cells / cm^2^ to 600,000 cells / cm^2^. Human peripheral blood mononuclear cell (PBMC) preparations were thawed, washed, centrifuged, and resuspended at 1 million (M) cells / ml in PBS. PBMC were labeled with CellTrace-Far Red (ThermoFisher, C34564) at 1:100 in PBS for 20min at 37°C. Following incubation, PBMC were quenched and washed with excess FBS (10%). Labelled PBMC were centrifuged and resuspended in RPMI with 10%FBS and added to VPC plated in 96-well for co-culture at 200K / well. After 72 hours co-culture, 200K CD3/CD28 activator beads (Fisher, 11131D) were added to wells. After 48 hours of activation, supernatant was collected, attached cells were harvested with Accutase, and the combined supernatant and attached cells were centrifuged and washed and prepared for flow cytometry.

***Flow cytometry assessment of T-cell proliferation and VPC immunophenotype***

For flow cytometry analysis of immunophenotyping of VPC after 48 hours of culture ± IFNγ stimulation and CD4^+^ and CD8^+^ T-cell proliferation after co-culture with VPC and activator beads, cells were prepared as a single cell suspension in FACS buffer (2% FBS containing D-Phosphate buffered saline (PBS). Cell surface staining was completed using the antibodies outlined in Supplementary Table 4. Cells were analyzed with a MACS quant analyzer 10 (Miltenyi Biotec). An unstained cell sample was used as control to establish a threshold for positive staining and subset gating.

***Generation of VPC conditioned media for myocyte cytoprotection***

Cryopreserved VPC were thawed and seeded at ~50K cells /cm^2^ on fibronectin coated 6-well plates (Corning, 354402) in complete Vasculife VEGF Endothelial medium (Lifeline Cell Technology, LL-003) for 24-hours. After 24-hours, media was changed to RPMI and VPC were cultured for an additional 48-hours. Media was then collected, centrifuged to remove any dead cells, and supernatants were either stored at -80°C prior to use for myocyte cytoprotection assays.

***Myocyte cytoprotection***

Human skeletal muscle myoblasts (HSMM) (Lonza, CC-2580, passage 3-5) were subcultured in SkBM-2 media (Lonza, CC-3246) until passage 3-5 according to manufacturer’s instructions. Myoblasts were then seeded at 15.6-23.4K/cm^2^/well in a 96-well plate (Corning, 3917) and cultured for four days in DMEM (ThermoFisher, 11885-084) supplemented with 2% horse serum (Gibco,16050-130) to differentiate them into myotubes. After 4 days, myotubes were incubated with either RPMI (control) or VPC conditioned media for an addition seven days in 1% O_2_ without media changes. After seven days, 100uL CellTiter-Glo® 3D (Promega, G9681) was added directly to the plate and luminescence was quantified. DMEM with 2% horse serum was used as a positive control and PBS as a negative control for cell survival. Data was normalized by the difference between positive and negative control for a given experiment / plate.

***THP-1 monocyte co-culture***

Cryopreserved VPC were thawed, counted, and plated at 91K/cm^2^/well (100K total cells/well) in a 48-well plate (Corning, 353078) in 250µL Complete Vasculife media for 24-hours. After 24-hours, media was changed to RPMI (ThermoFisher. 11875-093) with 20ng/mL IFNγ (PeproTech, 3000220UG) and 181K/cm^2^/well (200K) THP-1 monocytes (ATCC, TIB-202). Both unstimulated (i.e. THP1 cells in RPMI without VPC or IFNγ) and stimulated THP1 (20ng/mL IFNγ without VPC) were used as controls. Plates were then cultured in 1% O_2_ for 48-hours, conditioned media was collected, and secreted IL-6 was quantified using the human IL6 Quantikine ELISA kit according to manufacturer’s instructions (R&D Systems, D6050). IL6 concentration was normalized to conditioned media total protein as determined by the Pierce™ BCA protein assay kit (ThermoFisher, 23225) per manufacturer’s instructions. To control for variability between extent of stimulation between experiments, data was normalized by the average of a consistent VPC lot used as an assay control and the unstimulated and stimulated controls.

***H&E staining and immunofluorescence***

Quadriceps muscles from operated limbs were excised, fixed in 2.5% PFA, and embedded in paraffin. Whole slide analysis on 5µm sections was performed on H&E stained sections (Reveal Biosciences, San Diego, CA). For vascular density, muscle morphology, and VPC persistence analyses, 5µm sections were de-parafinized, rehydrated, and underwent heat-activated epitope retrieval (IHC-Tek, IW-1100). After permeabilization and blocking in 0.1% saponin (Alfa Aesar, 1004142) supplemented Carbofree blocking buffer (Vector Laboratories, SP-5040), slides were stained with primary antibody overnight at 4ºC. The following day, sections were washed 3-5x in PBS+Tween20, incubated 1-2 hours with appropriate secondary antibodies, washed and mounted with Vectashield Vibrance anti-fade mounting medium with DAPI (Vector Laboratories, H-1800). Slides were imaged on DMi8 epifluorescence microscope (Leica) at 10-20X.

For quantification of alpha smooth muscle actin (αSMA) area, 3-5 independent 10X ROIs were selected per animal in a blinded fashion. Total muscle area and αSMA-positive vessel area was quantified using semi-automated CellProfiler(2) image analysis pipeline in a blinded manner (CellProfiler).

***Clinical Hematology***

Blood was collected via the retro-orbital vein under isoflurane anesthesia into EDTA coated tubes. Hematological analysis of the peripheral blood was performed on the Hemavet HV950FS analyzer on the mouse setting (Drew Scientific). The number of total white blood cells (WBC), lymphocytes (LY), eosinophils (EO), neutrophils (NE), monocytes/macrophages (MO), basophils (BA), red blood cells (RBC) and platelets (PLT) were determined.

***Clinical Chemistry***

Blood was collected via cardiac puncture under isoflurane anesthesia. For serum preparation, the blood was allowed to coagulate for 30 minutes at room temperature and then centrifuged at 1000 × g for 15 minutes at 4^o^C. The Alfa Wasserman Vet Axcel clinical chemistry analyzer was used to measure the standard serum panel (15 analytes & 5 electrolytes).

***Tissue processing for gDNA and RNA isolation***

Frozen tissues were homogenized using a liquid nitrogen-cooled, stainless steel mortar (1600 MiniG, SPEX Sample Prep). Briefly, tissues were loaded into 5mL polyethylene tubes (ColeParmer sampleprep, #2240-PEF) prechilled in LN2, loaded with sterile, pre-chilled (in LN2), stainless steel grinding balls (ColeParmer Sampleprep, #2155), and loaded into the MiniG system. The MiniG was then run at maximum speed for 30s x 2. Tubes were then placed back in LN2 to enable aliquoting.

***gDNA isolation and hAlu PCR***

Genomic DNA (gDNA) was isolated from ~ 25 mg aliquots of powdered tissue using a DNeasy® Blood and Tissue kit (Qiagen, 69506) coupled with a QiaCube automated nucleic acid extractor (Qiagen) according to the manufacturer’s protocol. The degree of cellular engraftment was quantified using previously reported(3,4), using 15ng gDNA per reaction and custom TaqMan™ probe (FAM-ATTAGCCGGGCGTGGT) and primers (forward: 5’-CATGGTGAAACCCCGTCTCTA-3’; reverse: 5’-CTCGGCTCACTGCAACCTC-3’) designed to detect human ALU (hALU) elements on a on QuantStudio 7 Flex (Applied Biosystems) . To generate a standard curve, 5 ng/µL human gDNA was serially diluted over seven orders of magnitude (5 ng/µL, 5x10^-1^ ng/µL, 5x10^-2^ ng/µL, 5x10^-3^ ng/µL, 5x10^-4^ ng/µL, 5x10^-5^ ng/µL, and 5x10^-6^ ng/µL human gDNA) with 5 ng/µL mouse gDNA from an untreated control animal. Expression levels were calculated by minimal cycle threshold values (Ct) and quantified from standard curve.

***Total RNA Isolation and RNA Sequencing of mouse quadriceps muscle***

Frozen quadriceps muscles from db/db mice were processed as described above. Total RNA was isolated from ~25 mg aliquots of powdered tissue using the Trizol Plus RNA purification system (Invitrogen, 12183555). Isolated total RNA concentration was measured using a Nanodrop 2000 and quality checked using Agilent TapeStation. Libraries were prepared with poly-A enrichment and according to the NEBNext® Ultra™ II Directional RNA Library Prep kit for Illumina (New England Biolabs, Cat. No. E7490) using 1µg total RNA as input and 10 PCR amplification cycles. Paired-end reads 100 bp total for the pair of reads were sequenced using the NextSeq 2000. Reads were mapped to mm10 (refseq011015) mouse genome reference using STAR v.2.6.1, subjected to FastQC v.0.11.8 filtering, and gene/isoform expression levels were estimated using RSEM v.1.3.1. Transcripts per million reads (TPM) were used as a normalized unit of gene expression. Differentially expressed genes were identified using Matlab V.2023.1 with Bioinformatics Toolbox. Pathway analysis was performed using Ingenuity Pathway Analysis (IPA) using built-in modules (i.e. Disease Function, Canonical Pathways, Predicted Upstream Regulators) using genes with ≥ 2-fold differential expression.
