## Supplementary material for "Scalable Generation of Universal hiPSC-Derived Vascular Progenitor Cells for Safe and Sustained Revascularization in Chronic Limb-Threatening Ischemia": Supplemental Table 1_disease_function.pdf

**Supplemental Table 1. Ingenuity Pathway Analysis: Disease Function**

| Diseases or Functions Annotation | p-value | Predicted Activation State | Activation z-score |
|---|---|---|---|
| Insulin-dependent diabetes mellitus | 1.78E-15 | | -1.863 |
| Experimental colitis | 3.04E-15 | | -0.266 |
| Systemic autoimmune syndrome | 4.83E-15 | | -1.312 |
| Dextran sodium sulfate-induced colitis | 1.36E-14 | | -0.106 |
| Chronic inflammatory disorder | 5.83E-12 | | 0.042 |
| Inflammation of body cavity | 5.92E-12 | | 1.339 |
| Chronic colitis | 2.74E-11 | | |
| Acute colitis | 4.44E-11 | | |
| Infectious lung disease | 4.9E-11 | | |
| Inflammatory Bowel Disease | 8.45E-11 | | 0.021 |
| Infection of mammalia | 8.94E-11 | Increased | 2.435 |
| Bacterial pneumonia | 2E-10 | | |
| Late-onset hearing loss | 2.14E-10 | | |
| Diabetes mellitus | 4.02E-10 | Decreased | -2.301 |
| Cell movement of granulocytes | 9.04E-10 | Decreased | -4.114 |
| Colitis | 1.64E-09 | | -0.225 |
| Inflammation of organ | 2.18E-09 | | 0.54 |
| Glucose metabolism disorder | 3.26E-09 | Decreased | -2.51 |
| Early stage Staphylococcus aureus pneumonia | 3.35E-09 | | |
| Inflammation of gastrointestinal tract | 3.55E-09 | | 0.353 |
| Gastroenteritis | 3.65E-09 | | 0.561 |
| Inflammation of absolute anatomical region | 5.56E-09 | | 0.477 |
| Abnormality of large intestine | 6.67E-09 | | -0.45 |
| Enteritis | 7.97E-09 | | -0.035 |
| Infection by Staphylococcus aureus | 1.72E-08 | | |
| Recruitment of mononuclear leukocytes | 2.75E-08 | Decreased | -3.129 |
| Recruitment of lymphocytes | 0.000000045 | Decreased | -3.052 |
| Recruitment of neutrophils | 4.52E-08 | Decreased | -2.829 |
| Recruitment of granulocytes | 5.15E-08 | Decreased | -3.428 |
| Cellular infiltration by granulocytes | 5.47E-08 | Decreased | -3.193 |
| Cell movement of eosinophils | 8.84E-08 | Decreased | -2.936 |
| Symptomatic Huntington disease | 9.19E-08 | | |
| Cell movement of neutrophils | 0.000000256 | Decreased | -3.553 |
| Accumulation of leukocytes | 0.000000276 | Decreased | -2.659 |
| Accumulation of blood cells | 0.000000398 | Decreased | -2.752 |
| Recruitment of leukocytes | 0.000000455 | Decreased | -3.957 |
| Accumulation of myeloid cells | 0.000000704 | Decreased | -2.524 |
| Infection by Bacilli | 0.000000876 | | 0.54 |
| Quantity of IgG | 0.00000112 | | -0.031 |
| Recruitment of T lymphocytes | 0.0000014 | Decreased | -2.636 |
| Influx of leukocytes | 0.0000014 | | -1.058 |
| Damage of digestive system | 0.00000153 | | -0.858 |
| Inflammation of lung | 0.00000154 | | -0.241 |
| Accumulation of cells | 0.00000163 | Decreased | -2.729 |
| Recruitment of phagocytes | 0.00000172 | Decreased | -3.535 |
| Recruitment of myeloid cells | 0.00000202 | Decreased | -3.702 |
| Pneumonia | 0.00000209 | | |
| Multiorgan inflammation | 0.00000264 | | |
| Influx of cells | 0.00000291 | | -0.809 |
| Cell movement of myeloid cells | 0.00000318 | Decreased | -3.826 |
| Accumulation of mononuclear leukocytes | 0.00000335 | | -1.522 |
| Accumulation of phagocytes | 0.00000337 | | -1.867 |
| Response of CD4+ T-lymphocytes | 0.00000413 | | -0.895 |
| Atherosclerosis | 0.0000044 | Decreased | -2.374 |
| Cellular infiltration by myeloid cells | 0.00000473 | Decreased | -3.083 |
| Chemotaxis of leukocytes | 0.00000483 | Decreased | -3.785 |
| Homing of leukocytes | 0.00000559 | Decreased | -3.544 |
| Quantity of neutrophils | 0.00000569 | | -1.001 |
| T cell migration | 0.00000596 | Decreased | -2.935 |
| Cell movement of T lymphocytes | 0.00000624 | Decreased | -2.494 |
| Function of immune system | 0.00000686 | | |
| Rheumatoid arthritis | 0.00000693 | | -0.338 |
| Binding of blood vessel | 0.00000695 | Decreased | -2.069 |
| Influx of myeloid cells | 0.00000696 | | -1.032 |

| | | | |
|---|---|---|---|
| Onset of arthritis | 0.00000706 | | 0.896 |
| Migration of mononuclear leukocytes | 0.00000707 | Decreased | -3.557 |
| Quantity of IgA | 0.00000719 | | -0.246 |
| Infiltration by neutrophils | 0.00000728 | Decreased | -2.792 |
| Quantity of immunoglobulin | 0.00000778 | | 1.266 |
| Production of antibody | 0.00000806 | | 1.052 |
| Chemotaxis of granulocytes | 0.00000809 | Decreased | -2.667 |
| Quantity of cells | 0.00000842 | Decreased | -2.238 |
| Hepatic injury | 0.00000926 | | -1.577 |
| Migration of phagocytes | 0.0000105 | Decreased | -3.082 |
| Quantity of IgG1 | 0.0000111 | | -0.086 |
| Liver Damage | 0.0000136 | | -0.416 |
| Influx of phagocytes | 0.0000138 | | -1.164 |
| Occlusion of blood vessel | 0.0000151 | Decreased | -2.135 |
| Inflammation of respiratory system | 0.0000154 | | -0.766 |
| Quantity of granulocytes | 0.0000164 | | -1.613 |
| Cell movement of lymphoma cell lines | 0.0000173 | | -1.822 |
| Accumulation of granulocytes | 0.0000176 | | -1.307 |
| Production of protein | 0.0000176 | | 0.864 |
| Airway hyperresponsiveness | 0.0000179 | | -0.481 |
| Migration of lymphatic system cells | 0.0000195 | Decreased | -3.74 |
| Occlusion of artery | 0.0000227 | Decreased | -2.188 |
| Lymphocyte migration | 0.0000234 | Decreased | -3.635 |
| Inflammation of respiratory system component | 0.0000236 | | -0.69 |
| Vaso-occlusion | 0.0000238 | Decreased | -2.006 |
| Abnormal morphology of retinal layer | 0.0000246 | | |
| Accumulation of lymphocytes | 0.0000247 | | -1.701 |
| Familial neurological disorder | 0.000026 | | |
| Fertilization | 0.0000266 | Decreased | -2.213 |
| Polyneuropathy | 0.0000276 | | |
| Onset of experimentally-induced arthritis | 0.0000276 | | 0.447 |
| Quantity of IgG2a | 0.0000307 | | -0.102 |
| Homing of lymphoma cell lines | 0.0000313 | | -1.054 |
| Trafficking of cells | 0.0000343 | Decreased | -2.939 |
| Abnormal morphology of neurosensory retina | 0.0000351 | | |
| Cell movement of dendritic cells | 0.0000368 | Decreased | -2.583 |
| Cellular infiltration by blood cells | 0.0000371 | Decreased | -2.793 |
| Quantity of IgG3 | 0.0000371 | | -1.211 |
| Diabetic nephropathy | 0.0000397 | | |
| Sjögren syndrome | 0.0000426 | | |
| Activation of cells | 0.0000441 | | -1.756 |
| Cell movement of lymphatic system cells | 0.0000459 | Decreased | -3.829 |
| Cell-mediated response | 0.0000475 | | -1.144 |
| Quantity of protein in blood | 0.0000505 | | -0.692 |
| Abnormal function of neutrophils | 0.0000522 | | |
| Cell movement of phagocytes | 0.0000523 | Decreased | -3.791 |
| Non-traumatic arthropathy | 0.0000542 | | -0.675 |
| Trafficking of blood cells | 0.0000546 | Decreased | -2.765 |
| Quantity of helper T lymphocytes | 0.0000573 | | -1.285 |
| Accumulation of T lymphocytes | 0.0000596 | | -0.752 |
| Chemotaxis | 0.0000623 | Decreased | -3.805 |
| Inflammation of urinary tract | 0.0000625 | Increased | 2.556 |
| Size of vascular lesion | 0.0000625 | | |
| Quantity of metal | 0.0000661 | | -1.955 |
| Quantity of Th17 cells | 0.000071 | | -0.847 |
| Compartmentalization of cells | 0.0000723 | | |
| Accumulation of neutrophils | 0.0000733 | | -0.836 |
| Parasitic Infection | 0.0000745 | Increased | 2.01 |
| Size of atherosclerotic lesion | 0.0000776 | | |
| Cell movement of leukocytes | 0.0000804 | Decreased | -3.961 |
| Chemotaxis of lymphoma cell lines | 0.0000811 | | -1.671 |
| Activation of Phospholipase C | 0.0000811 | | |
| Mesoderm development | 0.000082 | | 0 |
| Quantity of IgG2b | 0.0000859 | | -1.028 |
| Commitment of cells | 0.0000861 | | -0.707 |
| Cellular infiltration by leukocytes | 0.0000881 | Decreased | -2.631 |

| | | | |
|---|---|---|---|
| Inflammation of joint | 0.0000899 | | -0.623 |
| Concentration of hormone | 0.000102 | | -0.188 |
| Erosion of ankle joint | 0.000103 | | |
| Atherosclerotic lesion | 0.000107 | | -1.213 |
| Migration of neutrophils | 0.000109 | Decreased | -2.626 |
| Differentiation of cytotoxic T cells | 0.000114 | | -1.159 |
| Chemotaxis of lymphatic system cells | 0.000121 | Decreased | -2.575 |
| Emotional behavior | 0.000121 | | -0.608 |
| Weight loss | 0.000125 | | 1.338 |
| Function of phagocytes | 0.000126 | | |
| Cell movement of lymphocytes | 0.000127 | Decreased | -3.733 |
| Mobilization of Ca2+ | 0.000129 | | -1.278 |
| Immune mediated inflammatory disease | 0.000131 | | -1.96 |
| Homing of cells | 0.000131 | Decreased | -3.648 |
| Chemotaxis of mononuclear leukocytes | 0.000136 | Decreased | -3.08 |
| Morphology of retina | 0.000136 | | |
| Migration of granulocytes | 0.000143 | Decreased | -2.917 |
| Cell movement | 0.000146 | Decreased | -3.389 |
| Migration of myeloid cells | 0.000147 | Decreased | -2.972 |
| Apoptosis of helper T lymphocytes | 0.000147 | | -0.762 |
| Recruitment of B lymphocytes | 0.000147 | | -1.432 |
| Activation of leukocytes | 0.000148 | Decreased | -2.427 |
| Nephritis | 0.000156 | Increased | 2.63 |
| Chemotaxis of neutrophils | 0.000156 | Decreased | -2.291 |
| Homing of mononuclear leukocytes | 0.000156 | Decreased | -3.226 |
| Development of ectoderm | 0.00017 | | |
| Spinocerebellar ataxia type 7 | 0.00017 | | |
| Trafficking of leukocytes | 0.00017 | Decreased | -2.586 |
| Inflammatory response | 0.000173 | Decreased | -3.627 |
| Movement of CD4+ T-lymphocytes | 0.000174 | | -1.65 |
| Binding of gonadal cells | 0.000175 | | -1.98 |
| Chemotaxis of myeloid cells | 0.000176 | Decreased | -3.603 |
| Cell movement of mononuclear leukocytes | 0.000181 | Decreased | -3.68 |
| Chemotaxis of phagocytes | 0.000186 | Decreased | -3.811 |
| Thickening of skin | 0.000187 | | -0.64 |
| Influx of granulocytes | 0.000191 | | -0.618 |
| Feeding | 0.000194 | | 0.132 |
| Function of leukocytes | 0.000209 | | 0.326 |
| Mobilization of leukocytes | 0.00021 | Decreased | -2.778 |
| Commitment | 0.000213 | | -0.788 |
| Cellular infiltration | 0.000216 | Decreased | -2.87 |
| Pancreatic cancer | 0.000219 | | -1.664 |
| Ingestion by mice | 0.00022 | | 1.234 |
| Homing of lymphatic system cells | 0.000224 | Decreased | -2.753 |
| Cellular infiltration by phagocytes | 0.000233 | Decreased | -2.704 |
| Erosion of joint | 0.000239 | | -0.372 |
| Onset of autoimmune disease | 0.000239 | | 0.055 |
| Synthesis of inositol phosphate | 0.00024 | | 0.464 |
| Inflammatory response of respiratory system component | 0.00024 | Decreased | -2.219 |
| Activation of DNA endogenous promoter | 0.000243 | Decreased | -2.879 |
| Cell movement of blood cells | 0.000244 | Decreased | -4.017 |
| Leukocyte migration | 0.000249 | Decreased | -3.949 |
| Clearance of cells | 0.000252 | | -1.997 |
| Influx of neutrophils | 0.000257 | | -0.711 |
| Quantity of anion | 0.000259 | | -1.06 |
| Cell tethering or rolling of leukocytes | 0.000259 | Decreased | -2.063 |
| Pancreatic adenocarcinoma | 0.000259 | | |
| Fungal Infection | 0.000272 | | -1.268 |
| Extravasation of leukocytes | 0.000274 | Decreased | -3.112 |
| Familial encephalopathy | 0.000278 | | |
| Quantity of regulatory T lymphocytes | 0.000278 | | -1.661 |
| Adhesion of venule | 0.000278 | | -1.932 |
| Accumulation of Th17 cells | 0.000278 | | -1.109 |
| Chemotaxis of eosinophils | 0.000278 | | -1.342 |
| Quantity of IgE | 0.000284 | | -0.853 |
| Familial amyloid polyneuropathy type I | 0.000291 | | |

| | | | |
|---|---|---|---|
| Recruitment of myeloid-derived suppressor cells | 0.000291 | | -1.972 |
| Ethanol aversion | 0.000291 | | |
| Onset of collagen-induced arthritis | 0.000291 | | 1 |
| Compartmentalization of leukocytes | 0.000294 | | |
| Transmembrane transport of ion | 0.000298 | | |
| Cellular infiltration by eosinophils | 0.000299 | Decreased | -2.117 |
| Migration of cells | 0.000301 | Decreased | -3.233 |
| Familial central nervous system disease | 0.000313 | | |
| Drinking | 0.000313 | | -1.154 |
| Narcolepsy with cataplexy | 0.000318 | | |
| Erosion of knee joint | 0.000318 | | |
| Intoxication | 0.000318 | | |
| Immune response of CD4+ T-lymphocytes | 0.000339 | | -0.816 |
| Emigration of leukocytes | 0.00034 | | -1.807 |
| Vascular lesion | 0.000345 | Decreased | -2.459 |
| Binding of germ cells | 0.000355 | | |
| Development of germ layer | 0.000361 | | 0 |
| Clearance of virus | 0.000379 | | -0.535 |
| Neuromuscular disease | 0.000383 | | -1 |
| Infiltration by T lymphocytes | 0.000388 | | -0.099 |
| Activation of blood cells | 0.000389 | Decreased | -2.604 |
| Development of genitourinary system | 0.000395 | Decreased | -2.114 |
| Malignant neoplasm of retroperitoneum | 0.000396 | | -1.709 |
| Aversive behavior | 0.000419 | | -1.214 |
| Quantity of leukotriene | 0.00042 | | -0.053 |
| Cytolysis | 0.000424 | | -0.472 |
| Binding of female germ cells | 0.000427 | | |
| Dyskinesia | 0.000434 | | |
| Chemotaxis of dendritic cells | 0.000439 | | -1.698 |
| Glomerulonephritis | 0.000443 | | 1.877 |
| Migration of Langerhans cells | 0.000443 | | -0.776 |
| Fertilization of ova | 0.000452 | | |
| Mobilization of blood cells | 0.000456 | Decreased | -2.375 |
| Mobilization of cells | 0.000464 | Decreased | -2.576 |
| Mobilization of myeloid cells | 0.000468 | Decreased | -2.63 |
| Huntington Disease | 0.000471 | | |
| Fusion of germ cells | 0.000493 | | |
| Function of helper T lymphocytes | 0.000516 | | |
| Development of reproductive system | 0.000522 | | -1.475 |
| Ophthalmia | 0.000527 | | 0.013 |
| Cell rolling of leukocytes | 0.000527 | | -1.255 |
| Quantity of B lymphocytes | 0.000529 | | -1.354 |
| Development of forelimb | 0.000546 | | |
| Infection of lung | 0.000546 | | 0.083 |
| Trafficking of T lymphocytes | 0.000546 | Decreased | -2.219 |
| Homing of tumor cell lines | 0.000547 | Decreased | -2.048 |
| Quantity of leukocytes | 0.000551 | Decreased | -3.254 |
| Pancreatic ductal adenocarcinoma | 0.000567 | | |
| Chemotaxis of lymphocytes | 0.000611 | Decreased | -2.606 |
| Development of genital organ | 0.000614 | | -1.668 |
| Function of antigen presenting cells | 0.000618 | | |
| Activation of mononuclear leukocytes | 0.000624 | Decreased | -3.085 |
| Function of blood cells | 0.000624 | | 0.326 |
| Accumulation of CD4+ T-lymphocytes | 0.000634 | | -1.408 |
| Modification of skeletal muscle | 0.000642 | | |
| Adhesion of postcapillary venule | 0.000642 | | -1.932 |
| Genodermatosis | 0.000642 | | |
| Response of Th17 cells | 0.000642 | | |
| Immune response of helper T lymphocytes | 0.000642 | | |
| Taste aversion | 0.000646 | | |
| Chemotaxis of tumor cell lines | 0.00065 | Decreased | -2.159 |
| Cell-cell adhesion | 0.000663 | | -1.891 |
| Area of lesion | 0.000668 | | |
| Extravasation | 0.000669 | Decreased | -3.536 |
| Migration of dendritic cells | 0.000669 | Decreased | -2.023 |
| Response of mononuclear leukocytes | 0.000695 | | -1.647 |

| | | | |
|---|---|---|---|
| Dorsal-ventral axis patterning | 0.000707 | | |
| Quantity of metal ion | 0.000707 | Decreased | -2.287 |
| Experimentally-induced arthritis | 0.00071 | | -0.028 |
| Morphology of eye | 0.000722 | | |
| Abnormal morphology of mouth | 0.000725 | | |
| Homing of lymphocytes | 0.000735 | Decreased | -2.78 |
| Development of hindlimb | 0.00075 | | |
| Development of sensory neurons | 0.000757 | | -1.432 |
| Sensory system development | 0.000791 | Decreased | -2.155 |
| Morphology of leukocytes | 0.000796 | | |
| Hearing loss | 0.000812 | | -0.908 |
| Response of helper T lymphocytes | 0.000842 | | -0.339 |
| Parasitemia | 0.000842 | | -0.132 |
| Homeostasis of D-glucose | 0.000845 | | |
| Quantity of interleukin | 0.000849 | | -1.445 |
| Ion homeostasis of cells | 0.000852 | Decreased | -2.339 |
| Inflammation of peritoneum | 0.000905 | | -0.761 |
| Abnormal morphology of immune system | 0.000906 | | |
| Delay in encephalomyelitis | 0.000913 | | |
| Cell-mediated response of CD4+ T-lymphocytes | 0.000913 | | -0.447 |
| Recruitment of dendritic cells | 0.000913 | Decreased | -2.425 |
| Synthesis of terpenoid | 0.000915 | | 0.174 |
| Immune response of leukocytes | 0.000922 | Decreased | -2.465 |
| Metabolism of eicosanoid | 0.000936 | Decreased | -2.481 |
| Development of endocrine gland | 0.000943 | | |
| Adhesion of germ cells | 0.000964 | | |
| Cytolysis of blood cells | 0.000966 | | -1.067 |
| Damage of lung | 0.000971 | | 0.35 |
| Abnormal morphology of retina | 0.000991 | | |
| Morphology of phagocytes | 0.000998 | | |
| Activation of lymphatic system cells | 0.00102 | Decreased | -2.533 |
| Quantity of sensory neurons | 0.00103 | | -1.372 |
| Locomotion | 0.00107 | | -0.179 |
| Hyperesthesia | 0.00107 | | -0.329 |
| Neuropeptide signaling pathway | 0.0011 | | |
| Transmigration of leukocytes | 0.00113 | Decreased | -3.191 |
| Chemoattraction of phagocytes | 0.0012 | Decreased | -2.19 |
| Congenital anomaly of skin | 0.0012 | | |
| Polarization of T-cell hybrid cells | 0.00121 | | |
| Lack of oligodendrocytes | 0.00121 | | |
| Formation of 9-cis-retinol | 0.00121 | | |
| Neovascularization of adductor muscle | 0.00121 | | |
| Zymosan-induced peritonitis | 0.00121 | | |
| Quantity of polymorphonuclear granulocytic MDSC cells | 0.00121 | | |
| Distribution of B lymphocytes | 0.00121 | | |
| Transformation of phagocytes | 0.00121 | | |
| Induction of naive T lymphocytes | 0.00121 | | |
| Formation of utricle | 0.00121 | | |
| Transformation of antigen presenting cells | 0.00121 | | |
| Proliferation of immune cells | 0.00121 | | -1.901 |
| Morphology of urothelium | 0.00121 | | |
| Spinal cord compression | 0.00121 | | |
| Experimental peritonitis | 0.00121 | | |
| Monocytosis | 0.00121 | Decreased | -2 |
| Damage of genitourinary system | 0.00122 | | -1.83 |
| Activation of innate lymphoid cells | 0.00122 | | -1.863 |
| Efflux of cholesterol | 0.00122 | | -0.21 |
| Abnormality of skin morphology | 0.00123 | | 0.592 |
| Efflux of lipid | 0.00124 | | -0.21 |
| Transcription of DNA | 0.00125 | Decreased | -2.767 |
| Diarrhea | 0.00125 | | 0.785 |
| Carditis | 0.00125 | | -0.726 |
| Metabolism of terpenoid | 0.00125 | | -0.013 |
| Function of dendritic cells | 0.00125 | | |
| Chemoattraction of leukocytes | 0.00126 | Decreased | -2.401 |
| Vascular leak syndrome of lung | 0.00126 | | 1.633 |

| | | | |
|---|---|---|---|
| Release of reactive oxygen species | 0.00126 | | -1.06 |
| Migration of helper T lymphocytes | 0.00126 | Decreased | -2.414 |
| Damage of bone | 0.0013 | | -1.23 |
| Abnormal function of immune system | 0.00132 | | |
| Response of lymphatic system cells | 0.00132 | | -1.998 |
| Hypersensitive reaction | 0.00134 | Decreased | -3.218 |
| Activation of lymphocytes | 0.00138 | Decreased | -2.739 |
| Frequency of regulatory T lymphocytes | 0.00138 | | -0.849 |
| Vision | 0.0014 | | |
| Activation of regulatory T lymphocytes | 0.00142 | | -1.706 |
| Abnormal morphology of ganglion cell layer | 0.00151 | | |
| Cell movement of helper T lymphocytes | 0.00151 | Decreased | -2.964 |
| Quantity of tumor cell lines | 0.00151 | | -0.104 |
| Disorder of basal ganglia | 0.00152 | | |
| Quantity of phagocytes | 0.00152 | Decreased | -2.718 |
| Hydrolysis of protein | 0.00153 | | 0 |
| Antimicrobial response | 0.00157 | | -1.698 |
| Cell movement of sperm | 0.00158 | | -0.563 |
| Recruitment of antigen presenting cells | 0.0016 | Decreased | -2.821 |
| Antibacterial response | 0.00163 | | |
| Quantity of Th1 cells | 0.00168 | | 0.088 |
| Autoimmune response | 0.00168 | Increased | 2.55 |
| Development of dermis | 0.00169 | | |
| Mobilization of granulocytes | 0.00176 | Decreased | -2.236 |
| Migration of antigen presenting cells | 0.00177 | Decreased | -2.786 |
| Neurological signs | 0.0018 | | 0.707 |
| Morphogenesis of forelimb | 0.0018 | | |
| Synthesis of leukotriene | 0.0018 | | -1.832 |
| Differentiation of neuroendocrine cells | 0.0018 | | |
| Extravasation of myeloid cells | 0.0018 | Decreased | -2.401 |
| Synthesis of leukotriene B4 | 0.0018 | | -1.964 |
| Steroid metabolism | 0.00184 | | 0 |
| Leukocytosis | 0.0019 | | -1.342 |
| Morphology of mouth | 0.00191 | | |
| Development of regulatory T lymphocytes | 0.00194 | | -0.193 |
| Proliferation of mononuclear leukocytes | 0.00203 | | -1.645 |
| Quantity of B-1 lymphocytes | 0.00206 | | 0.314 |
| Distribution of leukocytes | 0.00207 | | |
| Loss of retinal ganglion cells | 0.00207 | | |
| Hereditary retinal degeneration | 0.00207 | | |
| Chronic skin disorder | 0.00207 | | 1.067 |
| Chemoattraction of myeloid cells | 0.00207 | | -1.96 |
| Steatorrhea | 0.00207 | | 1 |
| Polarization of leukocyte cell lines | 0.00207 | | -1.976 |
| Hyperplasia of joint | 0.00207 | Decreased | -2 |
| Cytolysis by cytotoxic T cells | 0.00208 | | |
| Cell-mediated response of T lymphocytes | 0.00208 | | -1.504 |
| Insulitis | 0.00208 | | 0.863 |
| Activation of T lymphocytes | 0.00211 | Decreased | -2.008 |
| Function of myeloid cells | 0.00213 | | |
| Synthesis of steroid | 0.00213 | | 0.396 |
| Function of Th1 cells | 0.00214 | | |
| Morphogenesis of hindlimb | 0.00214 | | |
| Differentiation of neural cells | 0.00217 | Decreased | -2.077 |
| Differentiation of neurons | 0.00217 | | -1.347 |
| Quantity of IgM | 0.00219 | | -0.035 |
| Quantity of eosinophils | 0.00219 | | 0.086 |
| Morphology of macrophages | 0.00221 | | |
| Glycemic control | 0.00224 | | 0.555 |
| Immune response of T lymphocytes | 0.00226 | | -1.633 |
| Localization of cells | 0.00229 | Decreased | -2.726 |
| Recruitment of monocytes | 0.00242 | | -1.564 |
| Function of urinary bladder | 0.00249 | | |
| Cell rolling of lymphocytes | 0.00249 | | -0.849 |
| Loss of polysaccharide | 0.00249 | | -1.964 |
| Patterning of forebrain | 0.00249 | | |

| | | | |
|---|---|---|---|
| Accumulation of neuroglia | 0.00249 | | -0.555 |
| Activation of TREG cells | 0.00249 | | -1.091 |
| Migration of CD4+ T-lymphocytes | 0.00251 | | -1.994 |
| Proliferation of memory T lymphocytes | 0.00251 | | 1.274 |
| Priming of leukocytes | 0.00253 | Decreased | -2.726 |
| Specification of cells | 0.00256 | | -1.732 |
| Healing of lesion | 0.00267 | | |
| Neutrophilia | 0.0028 | | -1.562 |
| Quantity of retinal cells | 0.0028 | | -0.269 |
| Quantity of CD4+ T-lymphocytes | 0.00282 | | -0.611 |
| Cell-cell contact | 0.00286 | Decreased | -2.61 |
| Area of gap junction plaques | 0.00287 | | |
| Lack of mesencephalon | 0.00287 | | |
| Induction of Th2 cells | 0.00287 | | |
| Lack of intervertebral disc | 0.00287 | | |
| Light sensitivity of cells | 0.00287 | | |
| Formation of uterine gland | 0.00287 | | |
| Binding of microvessel | 0.00287 | | |
| Stimulation of hypothalamic neurons | 0.00287 | | |
| Quantity of glucocorticoid | 0.00287 | | |
| Quantity of lipoxin A4 | 0.00287 | | |
| Size of hair follicle | 0.00287 | | |
| Astrocytosis of cerebellum | 0.00287 | | |
| Meiosis of male germ cells | 0.00288 | | |
| Abnormal morphology of trabecular bone | 0.00288 | | |
| Growth of breast carcinoma | 0.0029 | | 1 |
| Differentiation of effector T lymphocytes | 0.0029 | | -0.686 |
| Formation of granuloma | 0.0029 | | -0.816 |
| Proliferation of oligodendrocyte precursor cells | 0.0029 | Decreased | -2 |
| Number of activated T lymphocyte | 0.0029 | | 1.342 |
| Mechanical allodynia behavior | 0.00293 | Decreased | -2.218 |
| Transmigration of phagocytes | 0.00293 | Decreased | -2.132 |
| Cytosis of blood cells | 0.00295 | | -1.59 |
| Function of gastrointestinal tract | 0.00298 | | |
| Abnormal morphology of middle ear ossicle | 0.00302 | | |
| Cellular infiltration by mononuclear leukocytes | 0.00302 | | -1.326 |
| Activation of myeloid cells | 0.00308 | | -1.895 |
| Retinal degeneration | 0.00309 | | 1.844 |
| Quantity of dendritic cells | 0.00317 | Decreased | -2.591 |
| Accumulation of microglia | 0.00326 | | -1 |
| Pupillary light reflex | 0.00326 | | 0.218 |
| Synthesis of glucocorticoid | 0.00326 | | |
| Quantity of steroid | 0.00331 | | 1.041 |
| Proliferation of lymphocytes | 0.00332 | | -1.747 |
| Response of lymphocytes | 0.00332 | Decreased | -2.176 |
| Chemotaxis of monocytes | 0.00341 | Decreased | -2.219 |
| Delay in experimental autoimmune encephalomyelitis | 0.00341 | | |
| Differentiation of Th9 cells | 0.00341 | | -0.808 |
| Infection by mycobacteria | 0.00341 | | 1.131 |
| Abnormal pigmentation of retina | 0.00341 | | |
| Localization of lymphocytes | 0.00341 | Decreased | -2.236 |
| Synthesis of eicosanoid | 0.00343 | Decreased | -2.286 |
| Adhesion of blood cells | 0.00345 | Decreased | -3.362 |
| Th17 immune response | 0.00346 | | -0.239 |
| Cleavage of carbohydrate | 0.00348 | | -1.124 |
| Abnormal quantity of Immunoglobulin | 0.00348 | | |
| Differentiation of central nervous system cells | 0.00355 | Decreased | -2.057 |
| Metabolism of alpha-amino acid | 0.0036 | | |
| Activation of granulocytes | 0.0036 | | -1.89 |
| Proliferation of blood cells | 0.0036 | | -1.986 |
| Aggregation of fibroblast cell lines | 0.0037 | | 0 |
| Quantity of IL-10 in blood | 0.0037 | | 0 |
| Extravasation of granulocytes | 0.0037 | Decreased | -2.19 |
| Quantity of blood cells | 0.00371 | Decreased | -3.289 |
| Migration of breast cancer cell lines | 0.00372 | Decreased | -2.517 |
| Proliferative capacity of regulatory T lymphocytes | 0.00372 | | -1.795 |

| | | | |
|---|---|---|---|
| Function of CD4+ T-lymphocytes | 0.00372 | | |
| Response of granulocytes | 0.00372 | | -1.209 |
| Expansion of lymphoid cells | 0.00372 | | -0.424 |
| Development of helper T lymphocytes | 0.00378 | | -1.376 |
| Generation of helper T lymphocytes | 0.00378 | | -1.509 |
| Quantity of interferon | 0.00378 | | -0.73 |
| Benign thyroid disease | 0.00378 | | 1.373 |
| Cell proliferation of T lymphocytes | 0.00379 | | -1.745 |
| Gonadogenesis | 0.00379 | | -1.668 |
| Loss of hair | 0.00386 | Increased | 2.046 |
| Transcription of RNA | 0.0039 | Decreased | -3.07 |
| Quantity of lymphoid cells | 0.00396 | Decreased | -2.679 |
| Cytolysis of lymphocytes | 0.00399 | | |
| Expansion of T lymphocytes | 0.00401 | | -0.561 |
| Quantity of virus | 0.00405 | Increased | 2.345 |
| Chemotaxis of T lymphocytes | 0.00405 | Decreased | -2.95 |
| Development of body axis | 0.00411 | | -1.117 |
| Transmigration of cells | 0.00412 | Decreased | -2.927 |
| Expansion of leukocytes | 0.0042 | | -0.941 |
| Migration of monocytes | 0.00421 | | -1.195 |
| Activation of connective tissue cells | 0.00425 | | -0.962 |
| Loss of bone | 0.00426 | | |
| Cellular infiltration by lymphocytes | 0.00428 | | -0.854 |
| Quantity of cytokine | 0.00438 | | -0.538 |
| Binding of myeloid cells | 0.0045 | Decreased | -2.598 |
| Induction of helper T lymphocytes | 0.00457 | | 0.047 |
| Frequency of TREG cells | 0.00457 | | -0.021 |
| Eosinophilia | 0.00457 | | -1.41 |
| Formation of hair | 0.00465 | | |
| Microcytic anemia | 0.00465 | | -0.314 |
| Accumulation of regulatory T lymphocytes | 0.00465 | | -0.555 |
| Acephaly | 0.00466 | | |
| Apoptosis of Th2 cells | 0.00466 | | |
| Acute tubular necrosis | 0.00466 | | |
| Accumulation of astrocytes | 0.00466 | | |
| Activation of cartilage tissue | 0.00466 | | |
| Adhesion of high endothelial postcapillary venule | 0.00466 | | |
| Accumulation of beta-estradiol | 0.00466 | | |
| Apoptosis of microvascular endothelial cells | 0.00466 | | |
| Apoptosis of activated T lymphocyte | 0.00466 | | |
