## Supplementary material for "Scalable Generation of Universal hiPSC-Derived Vascular Progenitor Cells for Safe and Sustained Revascularization in Chronic Limb-Threatening Ischemia": Supplemental Table 2_upstream regulators.pdf

**Supplemental Table 2. Ingenuity Pathway Analysis: Predicted Upstream Regulators**

| Upstream Regulator | Expr Log Ratio | Molecule Type | Predicted Activation State | Activation z-score |
|---|---|---|---|---|
| GATA2 | -0.027 | transcription regulator | Inhibited | -2.728 |
| RGS10 | 0.026 | enzyme | | 1.219 |
| ITPR2 | 0 | ion channel | Inhibited | -4.57 |
| PRDM1 | -0.321 | transcription regulator | | 1.625 |
| ZC3H12C | -0.282 | other | Activated | 2.591 |
| KRT17 | -0.185 | other | Inhibited | -2.5 |
| FOXA2 | 0 | transcription regulator | Activated | 2.64 |
| IFNAR1 | -0.222 | transmembrane receptor | | -1.925 |
| MRGPRX3 | 4.392 | G-protein coupled receptor | Inhibited | -3.771 |
| STAT1 | -0.669 | transcription regulator | Inhibited | -2.383 |
| HNF1A | -0.551 | transcription regulator | | -1.489 |
| IRF2BP2 | -0.057 | transcription regulator | Activated | 5.477 |
| NEIL2 | 0.346 | enzyme | | -0.816 |
| PARP1 | -0.152 | enzyme | Inhibited | -4.407 |
| C5 | -0.915 | cytokine | | 1.429 |
| CASP4 | -0.326 | peptidase | Inhibited | -4.016 |
| IL36A | | cytokine | Inhibited | -2.646 |
| IL17RA | -0.473 | transmembrane receptor | Inhibited | -2.515 |
| CLEC4E | -2.166 | other | Inhibited | -2.805 |
| PIK3CG | -0.623 | kinase | | 1.063 |
| STAT3 | -0.306 | transcription regulator | | -1.315 |
| GLI2 | -0.174 | transcription regulator | | |
| CXCL2 | -1.246 | cytokine | Inhibited | -2.593 |
| EPCAM | -2.115 | other | Activated | 3.289 |
| CCR5 | -0.574 | G-protein coupled receptor | | -0.98 |
| RNF31 | -0.05 | enzyme | Activated | 3.357 |
| CARD9 | -0.02 | other | Inhibited | -3.065 |
| SLC7A2 | -0.058 | transporter | Activated | 2.359 |
| TICAM1 | -0.188 | other | Inhibited | -4.606 |
| TLR7 | -0.31 | transmembrane receptor | Inhibited | -3.153 |
| N4BP1 | -0.241 | enzyme | Activated | 2.911 |
| CCR2 | -1.101 | G-protein coupled receptor | | -1.682 |
| HSPD1 | 0.037 | enzyme | Inhibited | -2.599 |
| TNF | -1.097 | cytokine | Inhibited | -5.193 |
| IFNB1 | 0 | cytokine | | -0.692 |
| CASP8 | -0.096 | peptidase | | 0.365 |
| PTX3 | -0.584 | other | Activated | 2.199 |
| IRF3 | -0.105 | transcription regulator | Inhibited | -2.491 |
| ADORA2B | -0.335 | G-protein coupled receptor | | -0.085 |
| TLR3 | -0.059 | transmembrane receptor | Inhibited | -4.14 |
| CDKN2A | -1.618 | transcription regulator | Inhibited | -3.221 |
| NFKBIA | 0.085 | transcription regulator | | -0.783 |
| COP1 | | enzyme | Activated | 3.286 |
| SERPINE2 | -0.036 | other | Activated | 2.743 |
| PCGF6 | -0.162 | transcription regulator | Activated | 2.203 |
| IRAK3 | -0.283 | kinase | | 1.251 |
| ABCG1 | -0.042 | transporter | Activated | 3.226 |
| IL11RA | -0.044 | transmembrane receptor | Inhibited | -3.113 |
| TLR2 | -0.488 | transmembrane receptor | Inhibited | -3.082 |
| TRPV4 | -0.245 | ion channel | Inhibited | -2.828 |
| RC3H1 | -0.152 | enzyme | | |
| RORA | 0.018 | ligand-dependent nuclear receptor | | -1.337 |
| IL1R1 | -0.502 | transmembrane receptor | Inhibited | -2.219 |
| PRKD1 | -0.229 | kinase | Inhibited | -2.11 |
| C3 | -0.726 | peptidase | Inhibited | -2.841 |
| ITGA5 | -0.321 | transmembrane receptor | Inhibited | -2.449 |
| ELANE | -0.174 | peptidase | Inhibited | -2.219 |
| JUNB | -0.667 | transcription regulator | | -0.712 |
| POU4F1 | -0.209 | transcription regulator | | -0.165 |
| LRP6 | -0.063 | transmembrane receptor | Activated | 2.078 |
| NR5A2 | -0.24 | ligand-dependent nuclear receptor | | 0.816 |
| PDK2 | -0.111 | kinase | | -0.762 |
| LRP5 | 0.018 | transmembrane receptor | | 1.746 |
| ALOX15 | -2.393 | enzyme | | -1.539 |
| IL15 | 0.197 | cytokine | | -1.886 |
| mir-21 | | microRNA | | 0.304 |
| VCAN | -0.696 | other | | -0.478 |
| IL10 | -1.721 | cytokine | Activated | 3.433 |
| IL12 (complex) | | complex | | -1.644 |
| PTAFR | -0.632 | G-protein coupled receptor | Activated | 2.236 |
| PILRA | -1.528 | other | | 1 |

| | | | | |
|---|---|---|---|---|
| IRF4 | -0.603 | transcription regulator | | 1.863 |
| CDX1 | -3.459 | transcription regulator | | 1.91 |
| LECT2 | -5.764 | other | | 0.577 |
| CTSB | -0.085 | peptidase | | -1.953 |
| Ciap | | group | Activated | 2.284 |
| EIF4B | -0.057 | translation regulator | | -0.207 |
| SIGIRR | 0.088 | transmembrane receptor | Activated | 2.62 |
| CRX | 0 | transcription regulator | | -1.407 |
| WTAP | -0.129 | other | | 1.999 |
| MIF | 0.062 | cytokine | | -1.497 |
| ITGA9 | -0.135 | other | Inhibited | -2.135 |
| CCL5 | -0.705 | cytokine | Inhibited | -2.793 |
| IL13 | 0.937 | cytokine | Inhibited | -3.673 |
| GPR84 | -7.301 | G-protein coupled receptor | Inhibited | -2.449 |
| SLC39A8 | 0.104 | transporter | | -0.105 |
| PLCE1 | -0.173 | enzyme | Inhibited | -2.412 |
| FEM1A | -0.038 | transcription regulator | Activated | 2.418 |
| Histone h3 | | group | | |
| Ige | | complex | Inhibited | -3.243 |
| TLR5 | -0.614 | transmembrane receptor | Inhibited | -2.771 |
| CD80 | -0.987 | transmembrane receptor | | -1.792 |
| NR1H3 | -0.054 | ligand-dependent nuclear receptor | | 0.914 |
| MMP9 | -1.041 | peptidase | | 1.281 |
| KDM6A | -0.063 | enzyme | | -1.023 |
| YTHDF1 | -0.068 | other | Activated | 3.13 |
| IKBKE | -0.702 | kinase | | -1.035 |
| ZFP36 | -0.254 | transcription regulator | | 1.91 |
| TRAF3IP2 | -0.252 | enzyme | Inhibited | -3.916 |
| IRF9 | -0.405 | transcription regulator | | -1.505 |
| LAMA5 | -0.157 | other | Inhibited | -2.813 |
| NLRP3 | -1.628 | enzyme | | -0.534 |
| PRKD | | group | Inhibited | -3.312 |
| DNASE2 | -0.137 | enzyme | Activated | 2.429 |
| PGLYRP2 | -0.348 | transmembrane receptor | Inhibited | -2.646 |
| PTGIS | -0.212 | enzyme | Inhibited | -2.646 |
| IL6 | -1.761 | cytokine | Inhibited | -2.175 |
| MAPK8 | -0.13 | kinase | | -0.845 |
| IL22 | 0 | cytokine | | -1.829 |
| FCGR2A | -0.391 | transmembrane receptor | Inhibited | -2.939 |
| TLR9 | -1.147 | transmembrane receptor | Inhibited | -3.63 |
| NFkB (complex) | | complex | Inhibited | -3.615 |
| SHH | -1.666 | peptidase | | 0.206 |
| Tlr | | group | Inhibited | -3.052 |
| STAT2 | -0.677 | transcription regulator | | -1.069 |
| HFE | -0.139 | transmembrane receptor | | 0.453 |
| CLEC10A | -0.725 | other | Activated | 2.401 |
| NR3C1 | -0.13 | ligand-dependent nuclear receptor | Activated | 3.097 |
| IL18 | -0.663 | cytokine | Inhibited | -2.805 |
| ZBTB16 | -0.349 | transcription regulator | | 0.368 |
| MAPK9 | -0.22 | kinase | Inhibited | -2.042 |
| FSTL1 | -0.401 | other | Inhibited | -2.383 |
| OGG1 | 0.188 | enzyme | | -1.682 |
| COCH | 0.09 | other | | -1.342 |
| LAT2 | -0.11 | other | | 1.751 |
| CTTN | -0.165 | other | Inhibited | -2.425 |
| TLN1 | -0.235 | other | Inhibited | -3.162 |
| IRF7 | -0.382 | transcription regulator | Inhibited | -4.094 |
| IL1A | -1.711 | cytokine | Inhibited | -3.495 |
| TLR8 | -0.278 | transmembrane receptor | | -1.762 |
| CLEC6A | -0.9 | transmembrane receptor | Inhibited | -2.216 |
| CYP27B1 | -0.437 | enzyme | Activated | 2.331 |
| SMARCB1 | -0.108 | transcription regulator | Inhibited | -2.412 |
| STING1 | | other | Inhibited | -2.493 |
| Fcer1 | | complex | Inhibited | -3.077 |
| NR1H4 | -0.383 | ligand-dependent nuclear receptor | Activated | 2.319 |
| PPP4R3B | | other | | 1.195 |
| KDM6B | -0.193 | enzyme | Inhibited | -2.63 |
| CLEC7A | -0.526 | transmembrane receptor | Inhibited | -2.727 |
| PTPN22 | -0.453 | phosphatase | | -0.104 |
| ARRB2 | -0.381 | other | | 0.64 |
| LCN2 | -0.815 | transporter | | -0.401 |
| TNFAIP8L2 | -0.375 | other | Activated | 2.599 |
| PRF1 | -0.71 | transporter | | 1.706 |

| | | | | |
|---|---|---|---|---|
| BPIFA1 | 0 | other | | 1.633 |
| NLRC5 | -0.923 | transcription regulator | | -0.277 |
| CD14 | -0.344 | transmembrane receptor | Inhibited | -2.177 |
| ZBTB10 | -0.1 | transcription regulator | Inhibited | -4.562 |
| PARK7 | 0.099 | enzyme | | -0.766 |
| EHMT2 | -0.122 | transcription regulator | | |
| mir-223 | | microRNA | | -1.259 |
| CLEC14A | 0.048 | other | Activated | 2 |
| HIVEP1 | -0.122 | transcription regulator | Activated | 2 |
| EIF2AK1 | -0.103 | kinase | | -1.987 |
| LUM | -0.264 | other | | -1.969 |
| CCL22 | 0.301 | cytokine | | 1.999 |
| PELI2 | -0.079 | enzyme | Inhibited | -2 |
| DUSP1 | -0.485 | phosphatase | Activated | 3.361 |
| RARB | -0.043 | ligand-dependent nuclear receptor | Inhibited | -3.178 |
| BCL3 | -0.868 | transcription regulator | Activated | 2.428 |
| SOCS1 | -0.699 | other | Activated | 2.409 |
| NR1H2 | -0.182 | ligand-dependent nuclear receptor | | 1.864 |
| ESR1 | 0.161 | ligand-dependent nuclear receptor | | -0.189 |
| YAP1 | -0.12 | transcription regulator | Inhibited | -2.824 |
| ITGAM | -0.613 | transmembrane receptor | Inhibited | -2.462 |
| ADA | -0.153 | enzyme | Activated | 2.772 |
| PON2 | -0.036 | enzyme | Activated | 2.236 |
| USP9X | -0.157 | peptidase | | 0 |
| IL1R2 | -0.635 | transmembrane receptor | Activated | 2.17 |
| IRAK2 | 0.139 | kinase | | |
| OTX2 | 5.102 | transcription regulator | | 0.543 |
| PPIF | 0.13 | enzyme | Inhibited | -2.065 |
| TRAF6 | -0.244 | enzyme | Inhibited | -2.303 |
| IL-1R | | group | Inhibited | -2.408 |
| C5AR2 | -0.539 | G-protein coupled receptor | | 1.087 |
| MVP | -0.12 | other | | -1.943 |
| POU4F2 | -3.841 | transcription regulator | | -1 |
| TNFRSF1A | -0.253 | transmembrane receptor | Inhibited | -2.812 |
| RIPK3 | -0.421 | kinase | Inhibited | -2.573 |
| Cyp2c70 | -0.442 | enzyme | | 1.555 |
| PDPK1 | -0.144 | kinase | | 0.393 |
| CXCR2 | -1.932 | G-protein coupled receptor | | 0.053 |
| RELA | -0.181 | transcription regulator | Inhibited | -3.107 |
| LHX1 | 3.841 | transcription regulator | | 0.477 |
| TREM2 | -0.041 | transmembrane receptor | | -0.488 |
| MAPKAPK2 | -0.217 | kinase | Inhibited | -3.515 |
| NFAM1 | -0.558 | transmembrane receptor | Inhibited | -2.401 |
| GIPR | 0 | G-protein coupled receptor | | |
| BRD2 | -0.167 | kinase | Inhibited | -2.449 |
| IFNG | -0.959 | cytokine | Inhibited | -5.305 |
| F2RL1 | -0.138 | G-protein coupled receptor | Inhibited | -2.204 |
| P38 MAPK | | group | Inhibited | -3.665 |
| S100A8 | -0.79 | other | | 1.225 |
| NFKBIZ | -0.439 | transcription regulator | Inhibited | -2.234 |
| PPP4R3A | | other | | 1.195 |
| MAP3K8 | -0.405 | kinase | Inhibited | -2.247 |
| ERK1/2 | | group | Inhibited | -3.254 |
| MSR1 | -0.847 | transmembrane receptor | | 0.787 |
| NT5E | 0.018 | phosphatase | Activated | 2.621 |
| PDK4 | 0.151 | kinase | | -0.762 |
| IL33 | -0.279 | cytokine | Inhibited | -3.328 |
| RIPK1 | -0.313 | kinase | Inhibited | -2.309 |
| PTF1A | 0 | transcription regulator | | 1.153 |
| SLC9A3R1 | -0.234 | transporter | Inhibited | -2.967 |
| AIRE | -4.392 | transcription regulator | | -0.397 |
| IL17A | 0 | cytokine | | -1.666 |
| STAT6 | -0.161 | transcription regulator | | 1.059 |
| IL27RA | -0.406 | transmembrane receptor | | 1.645 |
| CITED2 | 0.094 | transcription regulator | Activated | 4.96 |
| MAS1 | 0.169 | G-protein coupled receptor | | 1.096 |
| CXCL3 | -1.368 | cytokine | Inhibited | -2.425 |
| CCL11 | -0.219 | cytokine | Inhibited | -2.4 |
| EHMT1 | -0.142 | transcription regulator | | |
| NOD2 | -0.385 | other | Inhibited | -3.24 |
| Cyp4a14 | 2.939 | enzyme | Inhibited | -2.236 |
| SAA | | group | | -1.276 |
| PPT1 | -0.158 | enzyme | Activated | 2.236 |

| | | | | |
|---|---|---|---|---|
| HCAR1 | -0.084 | G-protein coupled receptor | | 1.037 |
| ADGRF5 | | G-protein coupled receptor | Activated | 2.219 |
| UNC5B | -0.136 | transmembrane receptor | Activated | 2.213 |
| CD5L | -0.897 | transmembrane receptor | | 0 |
| Gm20703 | | other | Activated | 2.236 |
| ITK | -0.559 | kinase | Inhibited | -3.001 |
| ITGAV | -0.121 | transmembrane receptor | Inhibited | -2.967 |
| AGER | -0.456 | transmembrane receptor | Inhibited | -2.408 |
| TIRAP | -0.003 | other | Inhibited | -2.433 |
| LTB | 0.014 | cytokine | Inhibited | -2.423 |
| SAA1 | -1.101 | transporter | Inhibited | -2.621 |
| RBP4 | -0.132 | other | Inhibited | -2.608 |
| FOXC1 | 0.123 | transcription regulator | | 0.302 |
| TERT | 0.11 | enzyme | Inhibited | -2.714 |
| ETS1 | -0.054 | transcription regulator | | -1.922 |
| EZH2 | -0.255 | transcription regulator | Inhibited | -2.475 |
| Srgn | -0.221 | other | Inhibited | -2.887 |
| ADIPOQ | 0.074 | other | | -0.561 |
| SUZ12 | -0.025 | enzyme | | -0.555 |
| TGFBR1 | -0.12 | kinase | Activated | 2.673 |
| HCAR2 | -0.148 | G-protein coupled receptor | Activated | 2 |
| CTSC | -0.267 | peptidase | | -1.698 |
| Trbv13-1 | | other | | |
| IKBIP | -0.149 | other | Activated | 2 |
| DLG1 | -0.099 | other | | -1.987 |
| AMBP | -2.089 | transporter | | 1.969 |
| EMX2 | -0.581 | transcription regulator | | 0.927 |
| GAS5 | -0.078 | other | | 1 |
| Orm1 (includes others) | 0.577 | other | | 1.972 |
| IL36RN | | cytokine | | |
| Wfdc17 | -0.427 | other | | 1.044 |
| FLT3LG | -0.179 | cytokine | Inhibited | -2.449 |
| IFNLR1 | -0.906 | transmembrane receptor | | |
| PTGS2 | -0.369 | enzyme | | -1.599 |
| IL1F10 | 0 | cytokine | | 0.314 |
| USP38 | -0.102 | peptidase | | 1.434 |
| PLA2G2D | -0.35 | enzyme | | 1.408 |
| STAP2 | 0.216 | other | Inhibited | -2.401 |
| IL10RA | -0.511 | transmembrane receptor | Activated | 2.961 |
| FPR2 | -2.244 | G-protein coupled receptor | | 1.363 |
| MYD88 | -0.426 | other | Inhibited | -4.986 |
| Klrk1 | -0.231 | transmembrane receptor | Inhibited | -2.447 |
| GSK3B | -0.094 | kinase | | -0.708 |
| AR | -0.095 | ligand-dependent nuclear receptor | | -0.034 |
| BCL2L11 | -0.293 | other | | 1.45 |
| PADI4 | -2.311 | enzyme | Inhibited | -2.2 |
| CCDC88A | 0.007 | other | Activated | 2.191 |
| TPSAB1/TPSB2 | -0.239 | peptidase | Inhibited | -2.207 |
| ISL1 | 3.841 | transcription regulator | | -1.038 |
| IFIH1 | -0.168 | enzyme | Inhibited | -2.621 |
| NCOA2 | -0.177 | transcription regulator | | 1.3 |
| PLAU | -0.274 | peptidase | Inhibited | -2.739 |
| TGFBR2 | -0.27 | kinase | Activated | 2.877 |
| LDLR | -0.457 | transporter | | 0.081 |
| TRAIP | -0.611 | enzyme | Activated | 2.585 |
| CNR2 | -0.159 | G-protein coupled receptor | | -0.649 |
| KITLG | 0.108 | growth factor | Inhibited | -2.172 |
| FOXP3 | -0.595 | transcription regulator | | 1 |
| JUN | -0.27 | transcription regulator | | -1.364 |
| Tcf7 | -0.2 | transcription regulator | | 0.419 |
| ASCL1 | -0.698 | transcription regulator | | 0.242 |
| CA4 | 0.327 | enzyme | | 0 |
| RPSA | -0.116 | translation regulator | Activated | 2.768 |
| CBL | -0.56 | transcription regulator | Activated | 2.789 |
| TRAF3IP3 | -0.204 | other | | -0.97 |
| P2RX4 | -0.275 | ion channel | Inhibited | -2.449 |
| EGR1 | -1.277 | transcription regulator | Inhibited | -3.06 |
| CTNNB1 | -0.072 | transcription regulator | | -0.893 |
| TBK1 | -0.13 | kinase | Inhibited | -2.025 |
| GSTP1 | -0.174 | enzyme | | |
| LGR4 | 0.015 | transmembrane receptor | | 0.391 |
| CEBPE | -2.349 | transcription regulator | | -0.783 |
| mir-130 | 0 | microRNA | | -0.57 |

| | | | | |
|---|---|---|---|---|
| ICOSLG/LOC102723996 | -0.235 | other | | -1.71 |
| WNT5A | -0.12 | cytokine | | -1.063 |
| CD28 | -0.287 | transmembrane receptor | | -1.927 |
| CD3 | | complex | Inhibited | -3.289 |
| BMP | | group | | -1.342 |
| F630028O10Rik | -0.451 | other | Activated | 2.213 |
| FRS3 | -0.176 | other | Inhibited | -2.236 |
| MMP14 | -0.375 | peptidase | | 1.231 |
| IL18R1 | -0.684 | transmembrane receptor | | 0.555 |
| CLEC4D | -0.58 | other | Inhibited | -2 |
| PLA2G4E | -0.368 | enzyme | Activated | 2 |
| HSPB1 | 0.055 | other | | 1.977 |
| F3 | -0.048 | transmembrane receptor | | |
| MARCO | -0.45 | transmembrane receptor | | -1.131 |
| F13A1 | -0.611 | enzyme | Inhibited | -2 |
| ENTPD1 | 0.003 | enzyme | Activated | 2 |
| SERPIND1 | 0.441 | other | Activated | 2 |
| Hrg | 0 | other | | 1.982 |
| WNT3A | -0.903 | cytokine | | -0.487 |
| PDK | | group | | |
| C1QTNF12 | | other | | |
| SCIMP | -0.552 | other | | |
| Snhg8 | 0.041 | other | | |
| SFRP5 | 0.42 | transmembrane receptor | | |
| COLGALT2 | 0.114 | enzyme | | |
| SCARB2 | -0.155 | transmembrane receptor | | |
| THBD | -0.214 | transmembrane receptor | | |
| Ccdc50 | 0.015 | other | | |
| PIP5K1B | 0.349 | kinase | | |
| LRG1 | -0.316 | other | | |
| VTN | 0.006 | other | | |
| CD300LD | -0.333 | other | | |
| ADAMDEC1 | 0.903 | peptidase | | |
| Supt20 | -0.333 | other | | |
| PROCR | -0.108 | other | | 0.608 |
| PEBP1 | -0.009 | other | | -0.39 |
| NR3C2 | -0.056 | ligand-dependent nuclear receptor | | -1.2 |
| PTGER4 | -0.149 | G-protein coupled receptor | Activated | 3.387 |
| FYN | -0.196 | kinase | | -1.795 |
| IL12B | -1.77 | cytokine | | 0.168 |
| Ccl2 | -1.285 | cytokine | Inhibited | -2.585 |
| NFATC2 | -0.164 | transcription regulator | | 0.501 |
| LTB4R | -0.551 | G-protein coupled receptor | | -1.526 |
| PTPRJ | -0.515 | phosphatase | Inhibited | -2.714 |
| Irgm1 | -0.423 | other | Activated | 4.018 |
| IFN alpha/beta | | group | | -1.863 |
| BCL11B | -0.433 | transcription regulator | | -0.417 |
| HIPK2 | -0.555 | kinase | | -0.177 |
| KLF11 | -0.027 | transcription regulator | | 1.575 |
| OSM | -1.478 | cytokine | Inhibited | -3.612 |
| IRF8 | -0.298 | transcription regulator | | 0.148 |
| ACKR2 | -0.062 | G-protein coupled receptor | Activated | 2.2 |
| SPP1 | -0.95 | cytokine | | -1.731 |
| PCSK9 | -1.035 | peptidase | | -1.98 |
| ATG16L1 | -0.05 | enzyme | | 1.964 |
| CXCR4 | -0.07 | G-protein coupled receptor | | -0.816 |
| FCER1A | -0.01 | transmembrane receptor | | -1.646 |
| SERPINB7 | 0.497 | other | | 1.673 |
| CCR6 | -1.919 | G-protein coupled receptor | | -1.667 |
| KIT | 0.099 | transmembrane receptor | | |
| EGLN | | group | Activated | 3.686 |
| GLI3 | -0.16 | transcription regulator | | -1.687 |
| Traj18 | | other | Activated | 2.191 |
| NOD1 | -0.182 | other | | |
| HRH2 | 0.349 | G-protein coupled receptor | | 0.156 |
| CCR3 | -1.324 | G-protein coupled receptor | | -0.447 |
| FYB1 | -0.626 | other | | 0.447 |
| TNNI3 | 0.1 | transporter | Inhibited | -2.236 |
| PPIA | 0.018 | enzyme | | 1.219 |
| CCL3L3 | -1.226 | cytokine | | -0.958 |
| SIRPA | -0.212 | phosphatase | Activated | 2.19 |
| KLF2 | -0.255 | transcription regulator | | 1.258 |
| Brd4 | -0.103 | kinase | Inhibited | -3.965 |

| | | | | |
|---|---|---|---|---|
| PTPN2 | -0.101 | phosphatase | Activated | 2 |
| GNAI2 | -0.144 | enzyme | | -1.964 |
| PTPRC | -0.503 | phosphatase | | -1.172 |
| Tnfsf9 | -0.866 | other | | -0.61 |
| NFAT5 | -0.174 | transcription regulator | | 0.559 |
| Map3k7 | -0.255 | kinase | | -1.362 |
| NFKB1 | -0.075 | transcription regulator | | -1.692 |
| STAT5A | -0.189 | transcription regulator | | -1.265 |
| NFKB2 | -0.094 | transcription regulator | | -0.472 |
| RIPK2 | -0.303 | kinase | Inhibited | -3.116 |
| CASP1 | -0.294 | peptidase | Inhibited | -2.486 |
| ITGB2 | -0.457 | transmembrane receptor | | 1.658 |
| Aldose Reductase | | group | Inhibited | -2 |
| ANKRD42 | -0.206 | transcription regulator | | -1.982 |
| PIMREG | | other | | -1.982 |
| Hoxa11os | 0.125 | other | | -1.982 |
| SARM1 | -0.756 | transmembrane receptor | Inhibited | -2 |
| S100B | 0.321 | other | | -1 |
| MDK | 0.185 | growth factor | | -1.982 |
| SELE | -0.884 | transmembrane receptor | | -0.105 |
| DGKZ | -0.06 | kinase | | -1.826 |
| PDE10A | -0.291 | enzyme | | -1.974 |
| MANF | -0.109 | other | Activated | 2 |
| S1PR1 | 0.298 | G-protein coupled receptor | | 0.988 |
| TRPA1 | 0 | transporter | | -1.982 |
| NDFIP1 | -0.04 | other | | 1.958 |
| TGFB1 | -0.111 | growth factor | | 0.87 |
| MAP2K1/2 | | group | Inhibited | -2.449 |
| ZBP1 | -1.097 | other | Inhibited | -2.414 |
| NBEAL2 | -0.212 | other | Activated | 2.425 |
| MAPKAPK3 | 0.038 | kinase | Inhibited | -2.412 |
| CXCR3 | -0.515 | G-protein coupled receptor | | -1.964 |
| IL4R | -0.225 | transmembrane receptor | | -1.442 |
| IL1B | -2.154 | cytokine | Inhibited | -3.832 |
| NOS2 | -0.084 | enzyme | | -1.538 |
| SLC6A4 | 0.016 | transporter | | 0.333 |
| LYN | -0.18 | kinase | | 1.7 |
| SGPP1 | -0.144 | phosphatase | | 1.342 |
| RUBCN | | other | | 0.447 |
| LOXL1 | -0.481 | enzyme | Inhibited | -2.236 |
| DEF6 | -0.108 | other | | 1.342 |
| mir-127 | 0 | microRNA | | -0.655 |
| TRPC1 | -0.256 | ion channel | Inhibited | -2.236 |
| F2 | -1.093 | peptidase | Inhibited | -2.216 |
| TNFSF12 | 0.01 | cytokine | Inhibited | -3.622 |
| ACOX1 | 0.017 | enzyme | | -0.475 |
| FOXC2 | 0.289 | transcription regulator | | 0 |
| RAG1 | 0 | enzyme | | -0.555 |
| BCL6 | -0.093 | transcription regulator | Activated | 3.522 |
| GLI1 | -0.299 | transcription regulator | | -1 |
| C1QTNF1 | -0.188 | other | | -0.237 |
| INHBA | -0.029 | growth factor | | 0 |
| ACE2 | 0 | peptidase | Activated | 2.449 |
| DDIT4 | 0.577 | other | Inhibited | -2.156 |
| G protein alpha i | | group | Inhibited | -3.582 |
| IRF1 | -0.369 | transcription regulator | | -1.692 |
| Hbb-b2 | | other | Inhibited | -2.985 |
| RNA polymerase II | | complex | | |
| Ifnar | | group | Inhibited | -3.445 |
| HTR7 | 0.359 | G-protein coupled receptor | | |
| PRRX2 | -0.078 | transcription regulator | | |
| ATP6V0D2 | -0.064 | transporter | | |
| TCF15 | 0.148 | transcription regulator | | |
| B3GNT2 | -0.036 | enzyme | | |
| SPATA2 | -0.187 | other | | |
| RNF152 | 0.243 | enzyme | | |
| HDGF | -0.092 | growth factor | | |
| MCOLN2 | -0.905 | ion channel | | |
| FLT4 | 0.036 | transmembrane receptor | | |
| CASP6 | -0.171 | peptidase | | |
| HYOU1 | -0.204 | other | | |
| ACVR2A | -0.014 | kinase | | |
| FGA | 0 | other | | |

| | | | | |
|---|---|---|---|---|
| SKAP2 | -0.297 | other | | |
| GNAI3 | -0.185 | enzyme | | |
| PPM1L | -0.218 | phosphatase | | |
| TRADD | -0.294 | other | | |
| PBLD | 0.094 | enzyme | | |
| IFITM3 | -0.39 | other | | |
| IL17F | -1.092 | cytokine | | |
| LATS1 | -0.273 | kinase | | -1.89 |
| FASLG | -6.454 | cytokine | | -0.943 |
| IL1 | | group | | -1.997 |
| ANKS6 | 0.017 | other | | 1 |
| KCNN4 | -0.35 | ion channel | | -1.98 |
| NGF | -0.01 | growth factor | | |
| PTGDR | 1.326 | G-protein coupled receptor | | 1.03 |
| SDC1 | -0.08 | enzyme | | 1.987 |
| Havcr1 | 1.145 | other | | 1.153 |
| IL36B | | cytokine | | |
| SCAVENGER receptor CLASS A | | group | | -1.432 |
| HPGDS | -0.117 | enzyme | Activated | 2.177 |
| IDO1 | -0.578 | enzyme | | 1.45 |
| ACE | -0.124 | peptidase | | 0 |
| CRLF2 | -0.261 | transmembrane receptor | | -1.446 |
| PLA2G2E | -0.982 | enzyme | Activated | 2.219 |
| CDX2 | 0 | transcription regulator | Activated | 2.095 |
| MALAT1 | 0.096 | other | | 0.335 |
| PDCD1 | -1.246 | transmembrane receptor | | 1 |
| FHL2 | 0.065 | transcription regulator | | -0.456 |
| CEBPD | 0.706 | transcription regulator | | -0.702 |
| EIF4EBP2 | -0.371 | translation regulator | Activated | 2.733 |
| Jnk | | group | | -1.35 |
| GATA6 | -0.373 | transcription regulator | | -0.79 |
| CREB3L3 | -0.031 | transcription regulator | Inhibited | -2.429 |
| CSF1R | -0.383 | kinase | | -1.633 |
| DUSP5 | -0.357 | phosphatase | | 1.633 |
| IKBKG | -0.162 | kinase | | -1.348 |
| PRKCD | -0.388 | kinase | | -1.483 |
| KCNK9 | 0 | ion channel | | 0.333 |
| Ttc39aos1 | | other | Activated | 3.419 |
| CNR1 | -0.289 | G-protein coupled receptor | Inhibited | -2.261 |
| OSMR | -0.331 | transmembrane receptor | | -1 |
| ZMPSTE24 | -0.126 | peptidase | Activated | 2.464 |
| CD86 | -0.613 | transmembrane receptor | | -1.424 |
| RAC1 | -0.049 | enzyme | | -0.895 |
| EIF4EBP1 | 0.004 | translation regulator | Activated | 2.733 |
| RORC | -0.302 | ligand-dependent nuclear receptor | | -1.387 |
| AHSG | 0.884 | other | | 0.333 |
| JAK2 | -0.069 | kinase | | -1.3 |
| NRL | 0 | transcription regulator | | -0.175 |
| FABP4 | 0.117 | transporter | Inhibited | -2.425 |
| SPHK1 | -0.315 | kinase | | -1.497 |
| ILKAP | -0.13 | phosphatase | | 1.698 |
| CLCN5 | -0.766 | ion channel | | 0.271 |
| BMP4 | 0.013 | growth factor | | -1.067 |
| IL36G | | cytokine | | |
| TNPO3 | -0.125 | other | | -0.447 |
| CRH | 1.937 | cytokine | | 0.277 |
| RPS6KA5 | -0.202 | kinase | | 0.626 |
| GPR37 | 0.709 | G-protein coupled receptor | | 1.342 |
| IRAK4 | -0.353 | kinase | Inhibited | -2.112 |
| BMPR1A | -0.046 | kinase | | -0.03 |
| CGAS | | enzyme | Inhibited | -2.401 |
| LATS2 | -0.152 | kinase | | -1.89 |
| ALOX5 | -1.051 | enzyme | | -1.89 |
| CHRNA7 | -0.105 | transmembrane receptor | Activated | 2.615 |
| MAPK7 | -0.245 | kinase | Inhibited | -2.63 |
| CD40 | -0.309 | transmembrane receptor | | -0.179 |
| cyclooxygenase | | group | | -1 |
| LDL | | complex | | -1.98 |
| CAVIN1 | | transcription regulator | | 1 |
| EIF4EBP3 | -0.07 | other | | 1.977 |
| TRAF5 | -0.071 | transporter | | 1.995 |
| NPY | -0.065 | other | | -1 |
| MAP3K2 | -0.22 | kinase | | -1.993 |

| | | | | |
|---|---|---|---|---|
| NLRC4 | -0.438 | other | Inhibited | -2 |
| TLR4 | -0.409 | transmembrane receptor | Inhibited | -4.077 |
| IL2 | 0 | cytokine | | -1.855 |
| IFNGR1 | 0.019 | transmembrane receptor | | -1.803 |
| TRAF3 | -0.315 | enzyme | | 0.874 |
| CHUK | -0.091 | kinase | Inhibited | -3.283 |
| CFTR | -0.823 | ion channel | | 0.104 |
| SMAD4 | -0.157 | transcription regulator | | 0.832 |
| Alpha catenin | | group | Activated | 3.471 |
| HMOX1 | -0.431 | enzyme | | 0.81 |
| IKZF2 | -0.064 | transcription regulator | Activated | 2.828 |
| Scd2 | -0.553 | enzyme | Activated | 2.813 |
| RUNX2 | -0.308 | transcription regulator | | -0.132 |
| TCF7L1 | -0.049 | transcription regulator | Inhibited | -2.714 |
| DUSP11 | -0.244 | phosphatase | Activated | 2.433 |
| IL1RL1 | -0.641 | transmembrane receptor | | -0.762 |
| SCGB1A1 | -1.335 | cytokine | Activated | 2.433 |
| mir-142 | 0 | microRNA | | 1.597 |
| IL23A | -0.105 | cytokine | | -1.678 |
| PTGS1 | -0.003 | enzyme | | -1.987 |
| APC | -0.335 | enzyme | | |
| TAFAZZIN | | enzyme | | -1.455 |
| HIF1A | -0.188 | transcription regulator | Inhibited | -2.735 |
| PANX1 | -0.211 | transporter | Inhibited | -2.236 |
| UNC93B1 | -0.249 | other | Inhibited | -2.177 |
| CTSK | -0.079 | peptidase | Inhibited | -2.2 |
| CRHR2 | 0.092 | G-protein coupled receptor | | -1 |
| TRIM21 | -0.404 | enzyme | | 1.192 |
| ELF3 | 0.307 | transcription regulator | | |
| FIBRINOGEN (family) | | group | | |
| CIRBP | -0.062 | translation regulator | | |
| GATM | 0.16 | enzyme | | |
| ITGB5 | 0.031 | other | | |
| PRRX1 | -0.134 | transcription regulator | | |
| SELL | -1.175 | transmembrane receptor | | |
| ACVR1 | 0.042 | kinase | | |
| mir-214 | | microRNA | | |
| FGF1 | -0.179 | growth factor | | |
| FGG | 0 | other | | |
| MMP13 | -2.8 | peptidase | | |
| PRSS8 | 1.495 | peptidase | | |
| RASGRP4 | -0.479 | other | | |
| SERPINB4 | 0 | other | | |
| STK38 | -0.115 | kinase | | |
| RGS16 | -0.502 | enzyme | | |
| TRAF1 | -0.226 | other | | |
| POLB | -0.073 | enzyme | | |
| CAPN1 | -0.064 | peptidase | | |
| mir-30 | 0 | microRNA | | |
| Mek | | group | | -1.356 |
| SMAD7 | -0.325 | transcription regulator | | 1.731 |
| ZEB2 | -0.278 | transcription regulator | | -1.134 |
| TCR | | complex | Inhibited | -3.091 |
| RETNLB | 0 | other | Inhibited | -3.592 |
| IL12A | -0.53 | cytokine | | 0.221 |
| KDM1A | -0.082 | enzyme | | |
| CBFB | 0.214 | transcription regulator | | 0.027 |
| FFAR3 | 0 | G-protein coupled receptor | Inhibited | -2 |
| IL1RL2 | -0.436 | transmembrane receptor | | -1.964 |
| AGK | -0.159 | kinase | Activated | 2 |
| SLC47A1 | -0.416 | transporter | | 1 |
| MAPK13 | 1.714 | kinase | | -1.029 |
| CYP2E1 | 0.676 | enzyme | Inhibited | -2 |
| TRPM2 | -1.669 | ion channel | | -1.919 |
| mir-27 | 0 | microRNA | | -0.936 |
| PLTP | -0.123 | enzyme | | -0.762 |
| SLC10A1 | -1.115 | transporter | | 0 |
| SWAP70 | -0.121 | other | Activated | 2 |
| NCF4 | -0.475 | enzyme | | 1 |
| CD180 | -0.444 | other | | |
| FRS2 | -0.069 | other | | |
| IL9 | 0 | cytokine | | -0.655 |
| EREG | -1.835 | growth factor | | -1.981 |

| | | | | |
|---|---|---|---|---|
| TLR6 | -0.162 | transmembrane receptor | | |
| CCND1 | -0.328 | transcription regulator | | -1.897 |
| C3AR1 | -0.43 | G-protein coupled receptor | Activated | 2.008 |
| CD274 | -1.23 | enzyme | | -0.73 |
| LGALS3 | -0.184 | other | | 0.387 |
| IL17C | 5.575 | cytokine | Inhibited | -2.207 |
| Tcrd | | other | Inhibited | -2.219 |
| RARRES2 | -0.31 | transmembrane receptor | Activated | 2.236 |
| TREM1 | -1.868 | transmembrane receptor | | -1.748 |
| LGALS1 | -0.024 | other | | 0.573 |
| FECH | -0.075 | enzyme | | |
| MARK2 | -0.223 | kinase | | -1.89 |
| TRIM24 | -0.032 | transcription regulator | | 1.678 |
| WDR77 | 0.047 | transcription regulator | | -0.294 |
| Notch | | group | | -1.195 |
| ERK | | group | | -0.263 |
| ADAM10 | -0.052 | peptidase | | -0.632 |
| STK40 | -0.419 | kinase | | -0.707 |
| CISH | 0.216 | other | Activated | 2.887 |
| CD44 | -0.404 | other | Inhibited | -2.049 |
| IFN Beta | | group | | -1.641 |
| MAVS | -0.22 | other | | -1.234 |
| BTK | -0.411 | kinase | | 0.423 |
| C5AR1 | -0.553 | G-protein coupled receptor | Inhibited | -2.035 |
| TCF12 | -0.142 | transcription regulator | Activated | 2.53 |
| VAV | | group | | |
| Dynamin | | group | | |
| Vacuolar H+ ATPase | | complex | | |
| RIOX1 | | enzyme | | |
| UPK3A | 0 | other | | |
| TAFA4 | | other | | |
| SPHKAP | 0.268 | other | | |
| ZDHHC12 | -0.18 | enzyme | | |
| ARL11 | -0.203 | other | | |
| IKKA/B | | group | | |
| FGR | -1.116 | kinase | | |
| MMP10 | -7.418 | peptidase | | |
| TPSG1 | -0.981 | peptidase | | |
| CTSG | 0.237 | peptidase | | |
| BAG4 | -0.09 | other | | |
| CLIP1 | -0.132 | other | | |
| mir-25 | 0 | microRNA | | |
| HOXB2 | -0.233 | transcription regulator | | |
| PHOX2A | 0.614 | transcription regulator | | |
| Ighe | | other | | |
| SDC4 | -0.299 | other | | |
| LGALS7/LGALS7B | 0.74 | other | | |
| IL1RAPL1 | -2.939 | transmembrane receptor | | |
| PPP1R13L | -0.178 | transcription regulator | | |
| GSC | 0.247 | transcription regulator | | |
| NASP | -0.137 | other | | |
| SLC7A11 | -0.923 | transporter | | |
| PRMT2 | -0.29 | enzyme | | |
| GNPAT | -0.164 | enzyme | | |
| ANPEP | -0.413 | peptidase | | |
| CD300C | -1.243 | transmembrane receptor | | |
| RELB | -0.114 | transcription regulator | | 0.64 |
| GDF11 | 0.131 | growth factor | Activated | 2.619 |
| MAPK14 | 0.056 | kinase | Inhibited | -2.425 |
| NPM1 | 0.013 | transcription regulator | | |
| LRBA | -0.094 | other | Activated | 2.236 |
| GLIS3 | -0.997 | transcription regulator | | |
| NODAL | 0.259 | growth factor | | 0.447 |
| ANGPTL2 | -0.06 | other | | -1.551 |
| IL6R | -0.001 | transmembrane receptor | Inhibited | -2.219 |
| BID | -0.222 | other | | -1.172 |
| IL15RA | -0.082 | transmembrane receptor | | -0.468 |
| CSF1 | -0.763 | cytokine | | -1.355 |
| PTGER2 | 0.259 | G-protein coupled receptor | Inhibited | -3.643 |
| INPP5D | -0.322 | phosphatase | | 1.11 |
| OLFM4 | -1.734 | other | | 1.98 |
| Mcpt4 | -0.344 | peptidase | | 0 |
| IL17RD | -0.066 | other | | 1.969 |

| | | | | |
|---|---|---|---|---|
| MUC13 | -0.733 | other | | 0 |
| HIRA | -0.191 | transcription regulator | Inhibited | -2 |
| VASP | -0.089 | other | | 1.981 |
| MAPK12 | -0.094 | kinase | | -1.029 |
| HDAC7 | -0.066 | transcription regulator | | -0.927 |
| SGK1 | -0.07 | kinase | | 1 |
| NR4A3 | -1.26 | ligand-dependent nuclear receptor | | -0.132 |
| YBX1 | -0.121 | transcription regulator | | 0.577 |
| FABP5 | -0.165 | transporter | | -1.599 |
| Gm21596/Hmgb1 | -0.016 | transcription regulator | Inhibited | -2.406 |
| PI3K (complex) | | complex | | -0.311 |
| VEGFA | -0.112 | growth factor | | -0.672 |
| IL4 | -6.658 | cytokine | Inhibited | -3.135 |
| CISD2 | -0.184 | other | | |
| IgG | | complex | | |
| MARCHF2 | | enzyme | | |
| Tlr13 | -0.402 | other | | |
| RGMB | -0.072 | other | | |
| USP39 | -0.127 | peptidase | | |
| UCP3 | 0.098 | transporter | | |
| TYR | -0.823 | enzyme | | |
| ABCD2 | -0.02 | transporter | | |
| MUC2 | -0.696 | other | | |
| LBP | -0.065 | transporter | | |
| RALBP1 | -0.169 | enzyme | | |
| PIAS4 | -0.126 | transcription regulator | | |
| ATP6V0A2 | -0.115 | transporter | | |
| SOD3 | -0.114 | enzyme | | |
| ARF6 | -0.17 | transporter | | |
| OXTR | 0.411 | G-protein coupled receptor | | |
| TNC | -0.646 | other | | |
| KLRB1 | -0.985 | transmembrane receptor | | |
| ABCB1 | 0.107 | transporter | | |
| USP8 | -0.223 | peptidase | Activated | 2.975 |
| ADCYAP1 | 0 | other | | 1.437 |
| HNF1B | 0 | transcription regulator | | |
| TNIP1 | -0.151 | other | | 1.219 |
| LEP | -0.036 | growth factor | | -1.492 |
| PRKCI | -0.186 | kinase | | 0.816 |
| LTA | 0.116 | cytokine | | -1.478 |
| SPHK2 | -0.15 | kinase | | -0.747 |
| CYBB | -0.424 | enzyme | | -0.807 |
| CAMP | 1.028 | other | | 0.062 |
| DLX6 | -0.292 | transcription regulator | | 0.663 |
| Cyp2c23 | | enzyme | | 0.555 |
| CD200 | -0.007 | other | Activated | 2.208 |
| DUSP10 | -0.304 | phosphatase | Activated | 2.198 |
| AXL | -0.354 | kinase | | 0.842 |
| SOCS6 | -0.429 | other | | -1.342 |
| BECN1 | -0.205 | other | | -1.603 |
| MST1R | -0.172 | kinase | Inhibited | -2.219 |
| IRF6 | 0.193 | transcription regulator | Activated | 2.236 |
| TET2 | -0.344 | enzyme | | 0.78 |
| MAFA | -0.206 | transcription regulator | | |
| RPTOR | -0.098 | other | | -0.799 |
| GATA3 | -0.403 | transcription regulator | | -1.461 |
| Esrra | -0.042 | ligand-dependent nuclear receptor | | -1 |
| SOCS3 | -0.677 | phosphatase | Activated | 2.459 |
| ICOS | -1.198 | transmembrane receptor | | -0.716 |
| IKBKB | -0.061 | kinase | Inhibited | -3.25 |
| TNFSF4 | -0.865 | cytokine | | -1.991 |
| LAT | -0.572 | other | | -1.623 |
| CD19 | -2.027 | transmembrane receptor | | -1.982 |
| PTGIR | -0.329 | G-protein coupled receptor | | -1 |
| RPS6KA4 | -0.218 | kinase | | 1.029 |
| ODC1 | 0.031 | enzyme | Activated | 2 |
| PCSK2 | 0.217 | peptidase | | 0.577 |
| FLT3 | -0.907 | kinase | | |
| GAB1 | 0.051 | other | | -1.941 |
| TNFRSF1B | -0.512 | transmembrane receptor | Inhibited | -2.382 |
| IKZF1 | -0.533 | transcription regulator | | 0.721 |
| TCF3 | -0.149 | transcription regulator | | -0.077 |
| TNFAIP3 | -0.331 | enzyme | | 1.623 |

| | | | | |
|---|---|---|---|---|
| JUND | -0.122 | transcription regulator | | |
| PRKCQ | -0.159 | kinase | Inhibited | -2.19 |
| SLC9A3 | -0.592 | ion channel | | |
| HRH3 | 3.841 | G-protein coupled receptor | | |
| HES1 | 0.029 | transcription regulator | | 0.239 |
| OPA1 | -0.009 | enzyme | Activated | 2.213 |
| MOG | -4.605 | other | Inhibited | -2.236 |
| IFI16 | -4.143 | transcription regulator | | -1.067 |
| IL1RN | -0.301 | cytokine | | 1.558 |
| ABCA1 | -0.436 | transporter | Activated | 2.772 |
| LRP1 | -0.483 | transmembrane receptor | Activated | 2.397 |
| TCF4 | 0.102 | transcription regulator | | -0.721 |
| NR1I2 | 3.841 | ligand-dependent nuclear receptor | | 0 |
| RNF2 | -0.022 | transcription regulator | | 1.342 |
| YTHDC1 | -0.014 | other | | |
| Cd209b | -0.084 | other | | |
| 4732491K20Rik | -0.09 | other | | |
| SLC16A3 | 0.099 | transporter | | |
| CXCL17 | -0.164 | cytokine | | |
| PNPLA7 | 0.144 | enzyme | | |
| TNKS2 | -0.176 | enzyme | | |
| TCF21 | -4.143 | transcription regulator | | |
| EZH1 | -0.094 | enzyme | | |
| AVPR1A | -0.005 | G-protein coupled receptor | | |
| LY96 | 0.151 | transmembrane receptor | | |
| PROC | 0 | peptidase | | |
| TGFA | 0.275 | growth factor | | |
| CR1L | -0.018 | transmembrane receptor | | |
| TNFRSF11A | -0.428 | transmembrane receptor | | |
| C6 | 0.296 | other | | |
| SFTPA1 | 0.764 | transporter | | |
| HADH | 0.142 | enzyme | | |
| Ear2 (includes others) | -6.08 | enzyme | | |
| IL18BP | -0.323 | other | | |
| TNFAIP8 | -0.196 | other | | |
| GBX2 | -0.739 | transcription regulator | | |
| ST14 | 0.219 | peptidase | | |
| SP7 | 1.495 | transcription regulator | | |
| BMP2 | -0.008 | growth factor | | |
| Casp12 | -0.283 | peptidase | | |
| NEDD4L | -0.099 | enzyme | | |
| MYF6 | -0.063 | transcription regulator | | |
| MSX2 | 6.007 | transcription regulator | | -1.075 |
| BATF | -0.961 | transcription regulator | | |
| PLK4 | -0.535 | kinase | | 0 |
| RHO | -1.372 | G-protein coupled receptor | Inhibited | -2.333 |
| RUNX3 | -0.476 | transcription regulator | | |
| DLX2 | 3.841 | transcription regulator | | |
| ITGA1 | -0.141 | other | | -0.594 |
| DOCK2 | -0.531 | other | Inhibited | -2 |
| THRA | -0.22 | ligand-dependent nuclear receptor | | -0.165 |
| KCNJ10 | -0.872 | ion channel | Inhibited | -2.236 |
| GADD45A | 0.24 | other | Activated | 2.216 |
| MAP2K3 | 0.046 | kinase | | -1.96 |
| IFIT2 | -0.351 | other | | -1.091 |
| TRAF2 | -0.109 | enzyme | | 1.322 |
| HNF4A | 2.115 | transcription regulator | | 1.836 |
| CLIC4 | -0.219 | ion channel | | -1.404 |
| OGA | | enzyme | | -1.279 |
| MTTP | -0.305 | transporter | | |
| GTF2IRD1 | -0.048 | transcription regulator | | -0.398 |
| RUNX1 | -0.417 | transcription regulator | | 0.854 |
| TP63 | -0.384 | transcription regulator | | 1.777 |
| WLS | -0.131 | other | | 1.342 |
| CAMK4 | 0.158 | kinase | | -1.458 |
| HAVCR2 | -0.6 | other | | 0.447 |
| KEAP1 | -0.15 | other | | -1.091 |
| DLX5 | -0.42 | transcription regulator | | 0.648 |
| IRAK1 | -0.101 | kinase | | -1.983 |
| STAT5B | -0.06 | transcription regulator | | -1.454 |
| Yaf2 | -0.045 | transcription regulator | | -1.673 |
| JINK1/2 | | group | | -1.982 |
| Fc gamma receptor | | group | | -0.068 |

| | | | | |
|---|---|---|---|---|
| L2HGDH | 0.002 | enzyme | Activated | 2 |
| TNFRSF18 | -0.288 | transmembrane receptor | | -1.925 |
| ADRB3 | 0.444 | G-protein coupled receptor | | -1.989 |
| PIKFYVE | -0.186 | kinase | | -0.762 |
| HR | -0.337 | transcription regulator | | -1 |
| PTPRE | -0.386 | phosphatase | | -1 |
| WAS | -0.364 | other | | |
| BCL10 | -0.089 | transcription regulator | | -1.957 |
| ASIP | 0.829 | other | | |
| FCER1G | -0.331 | transmembrane receptor | | -1.934 |
| Klra7 (includes others) | -5.358 | transmembrane receptor | | |
| MAP1LC3 | | group | | |
| IKK (complex) | | complex | | |
| Pdgf Ab | | complex | | |
| PDGF BB | | complex | | |
| NEUROD4 | 0 | transcription regulator | | |
| GBP5 | -0.782 | enzyme | | |
| CERS5 | -0.113 | transcription regulator | | |
| HECTD3 | -0.099 | enzyme | | |
| INPP4B | -0.05 | phosphatase | | |
| SMCR8 | -0.183 | other | | |
| GJB6 | -0.204 | transporter | | |
| COQ7 | 0.11 | enzyme | | |
| SIRT4 | 0.113 | enzyme | | |
| WNT9B | -0.087 | other | | |
| NAAA | -0.316 | enzyme | | |
| AMH | -1.21 | growth factor | | |
| AZI2 | -0.111 | other | | |
| DHX15 | -0.118 | enzyme | | |
| STX11 | -0.638 | transporter | | |
| KHSRP | -0.136 | enzyme | | |
| KLK3 | 6.15 | peptidase | | |
| Cd24a | 0.161 | other | | |
| HCK | -0.441 | kinase | | |
| 2610528A11Rik | 0.554 | cytokine | | |
| MEP1A | 0 | peptidase | | |
| SEMA3E | -0.362 | other | | |
| P2RY12 | -0.798 | G-protein coupled receptor | | |
| CEBPG | -0.094 | transcription regulator | | |
| RAB10 | -0.052 | enzyme | | |
| SLC6A14 | 0 | transporter | | |
| CAMK2G | -0.054 | kinase | | |
| SENP6 | -0.083 | peptidase | | |
| C4A/C4B | -0.592 | other | | |
| GJC1 | -0.048 | ion channel | | |
| IRAK1BP1 | -0.164 | other | | |
| Trim30a/Trim30d | -0.446 | enzyme | | |
| ANK1 | -0.076 | other | | |
| mir-467 | 13.562 | microRNA | | |
| SORT1 | -0.075 | G-protein coupled receptor | | |
| DHCR7 | -0.361 | enzyme | | |
| ERO1A | | enzyme | | |
| CAMK2D | -0.037 | kinase | | |
| P2RX7 | -0.372 | ion channel | | |
| Ppp1cc | -0.147 | phosphatase | | |
| PTPN12 | -0.109 | phosphatase | | |
| CYP7A1 | -3.459 | enzyme | | |
| Tgf beta | | group | | 1.633 |
| SETDB1 | -0.155 | enzyme | | |
| mir-182 | 0 | microRNA | | -1.673 |
| TXNIP | 0.205 | other | | -0.946 |
| BCAP31 | -0.014 | transporter | | 1.141 |
| TRIM38 | -0.409 | enzyme | | |
| MTF1 | -0.237 | transcription regulator | | |
| PELI1 | -0.09 | enzyme | | |
| KLF9 | 0.103 | transcription regulator | | |
| EFNA5 | 0.023 | kinase | | |
| B4GALT1 | -0.169 | enzyme | | |
| MBIP | 0.137 | other | | |
| ELP3 | -0.016 | enzyme | | |
| mir-15 | 0 | microRNA | | |
| CSK | -0.259 | kinase | | |
| SCRIB | -0.143 | other | | |

| | | | | |
|---|---|---|---|---|
| PLAUR | -0.539 | transmembrane receptor | | |
| FOXD1 | 0.054 | transcription regulator | | |
| ADORA1 | 0.104 | G-protein coupled receptor | | |
| KLB | -0.069 | enzyme | | |
| MNX1 | 0 | transcription regulator | | |
| TNKS | -0.279 | enzyme | | |
| ITGAL | -1.296 | transmembrane receptor | | |
| ADAM12 | -0.702 | peptidase | | 1.315 |
| GFI1 | 0.038 | transcription regulator | Activated | 2.231 |
| GPD1 | -0.111 | enzyme | | |
| FIGLA | 0 | transcription regulator | | -0.308 |
| TBX21 | -0.38 | transcription regulator | | -0.453 |
| BMP10 | 0 | growth factor | Inhibited | -2.309 |
| PROP1 | -0.213 | transcription regulator | | 0.447 |
| SOX17 | 0.092 | transcription regulator | | |
| LILRB4 | | other | | 1.342 |
| MDM2 | -0.151 | transcription regulator | | |
| MALT1 | -0.179 | peptidase | Inhibited | -2.382 |
| EIF2AK2 | -0.335 | kinase | Inhibited | -2.207 |
| Histone h4 | | group | | |
| NAMPT | -0.062 | cytokine | | -0.125 |
| ITGB1 | -0.047 | transmembrane receptor | Inhibited | -2.308 |
| BTNL2 | -0.858 | transmembrane receptor | | -1.508 |
| SHARPIN | -0.06 | other | | 0.625 |
| TICAM2 | -0.484 | other | | -1.154 |
| CD70 | 4.605 | cytokine | | |
| MGAT5 | -0.497 | enzyme | | 0.762 |
| TNK1 | 0.536 | kinase | | -1.982 |
| SLC25A13 | -0.267 | transporter | | |
| CBFA2T3 | -0.154 | transcription regulator | | -1.195 |
| PAX6 | -0.147 | transcription regulator | | -1.019 |
| LTBR | -0.135 | transmembrane receptor | | -0.728 |
| ESR2 | 0.09 | ligand-dependent nuclear receptor | | 0.251 |
| DMRT1 | 0 | transcription regulator | | -0.97 |
| IHH | -0.91 | enzyme | | 1.035 |
| NR0B2 | 0.774 | ligand-dependent nuclear receptor | | 1.339 |
| IL27 | 0.368 | cytokine | | 0.573 |
| IRF2 | -0.127 | transcription regulator | | -1.97 |
| DIO2 | -0.456 | enzyme | | 1.257 |
| SSB | 0.045 | enzyme | | 1.342 |
| ZEB1 | -0.175 | transcription regulator | Inhibited | -2 |
| PARP2 | -0.157 | enzyme | Inhibited | -2.229 |
| GRN | -0.167 | growth factor | | 0.596 |
| EGFR | -0.304 | kinase | | -1.525 |
| CLEC4G | -0.895 | other | | 0.788 |
| NCOR1 | -0.128 | transcription regulator | Activated | 2 |
| PTPN6 | -0.692 | phosphatase | | 1.302 |
| RSPO3 | 0.027 | kinase | | |
| TMEM79 | 0.098 | other | | |
| ABCD1 | 0.003 | transporter | | |
| GPR132 | -0.889 | G-protein coupled receptor | | |
| CD81 | 0.046 | other | | |
| AGT | 0.186 | growth factor | | |
| mir-17 | 0 | microRNA | | |
| LGALS3BP | -0.309 | transmembrane receptor | | |
| MAP3K5 | -0.13 | kinase | | |
| SELP | -0.228 | transmembrane receptor | | |
| BMX | 0.105 | kinase | | |
| WWOX | -0.138 | enzyme | | |
| GAST | 0 | other | | |
| FOSL1 | -1.573 | transcription regulator | | |
| KLF13 | -0.188 | transcription regulator | | |
| Hotairm1 | | other | | |
| MRTFB | | transcription regulator | | -0.302 |
| NR4A1 | 0.221 | ligand-dependent nuclear receptor | | 0.592 |
| KCNIP3 | -0.099 | transcription regulator | | -1.414 |
| GRIN1 | 0 | ion channel | | 0 |
| EZR | -0.256 | other | | -0.728 |
| TGIF1 | -0.205 | transcription regulator | Inhibited | -2 |
| MAP3K1 | -0.296 | kinase | | -1.131 |
| IFNA4 | -6.508 | cytokine | | -1.982 |
| GNRH1 | -0.692 | other | | |
| SNAI2 | -0.025 | transcription regulator | Activated | 2 |

| | | | | |
|---|---|---|---|---|
| SRF | -0.099 | transcription regulator | | -0.715 |
| DIO3 | -0.549 | enzyme | | 0.686 |
| PLCG2 | -0.643 | enzyme | | 0.304 |
| XBP1 | -0.043 | transcription regulator | | -0.609 |
| CLDN6 | 0 | other | | |
| HSP90B1 | -0.09 | other | Inhibited | -2.168 |
| FGFR2 | 0.573 | kinase | | 0.64 |
| TNFRSF9 | -1.543 | transmembrane receptor | | 1.698 |
| STAT4 | -0.251 | transcription regulator | | -1.924 |
| EGF | -0.069 | growth factor | | |
| MRTFA | | transcription regulator | | -0.687 |
| SCD | -0.607 | enzyme | | 1.968 |
| NEDD9 | 0.031 | other | | -0.707 |
| KL | -0.348 | enzyme | | 1.39 |
| MLX | -0.158 | transcription regulator | | 0.816 |
| PILRB | -0.86 | other | | |
| Calcineurin B | | group | | |
| RAB7B | | peptidase | | |
| ENPP7 | 0 | enzyme | | |
| Carlr | | other | | |
| PHLDA2 | 0 | other | | |
| ATOH7 | -1.235 | transcription regulator | | |
| SLC22A3 | -0.188 | transporter | | |
| RNF122 | -0.52 | enzyme | | |
| PON3 | -0.025 | enzyme | | |
| KLHL22 | 0.071 | other | | |
| PARG | -0.089 | enzyme | | |
| PRKAB2 | 0.122 | kinase | | |
| TRAFD1 | -0.194 | other | | |
| CD6 | -1.04 | transmembrane receptor | | |
| CHST4 | -6.08 | enzyme | | |
| CPB2 | 0 | peptidase | | |
| PHF19 | 0.007 | other | | |
| mir-455 | 0 | microRNA | | |
| USP7 | -0.105 | peptidase | | |
| SLC12A3 | -2.939 | transporter | | |
| CYP8B1 | -1.865 | enzyme | | |
| PLCB3 | -0.093 | enzyme | | |
| ACSL4 | -0.179 | enzyme | | |
| A4GALT | -0.098 | enzyme | | |
| MMP7 | 0 | peptidase | | |
| TRU-TCA1-1 | | other | | |
| CCL4 | -1.506 | cytokine | | |
| SIGLEC1 | -0.761 | other | | |
| LHX6 | 0.086 | transcription regulator | | |
| FGF19 | 3.459 | growth factor | | |
| F2RL3 | 0.183 | G-protein coupled receptor | | |
| DLGAP3 | 0.136 | other | | |
| CHST2 | 0.337 | enzyme | | |
| NPY5R | 0 | G-protein coupled receptor | | |
| CAMK2A | -0.044 | kinase | | |
| ANXA6 | -0.115 | ion channel | | |
| DNM2 | -0.173 | enzyme | | |
| CX3CL1 | -0.199 | cytokine | | -0.333 |
| CTCF | -0.128 | transcription regulator | | 0.816 |
| BHLHE40 | -0.244 | transcription regulator | Inhibited | -3.623 |
| KDM4C | -0.092 | enzyme | | |
| ICAM1 | -0.383 | transmembrane receptor | | -1.96 |
| XDH | -0.37 | enzyme | | -0.577 |
| EPHX2 | 0.34 | enzyme | | -1.98 |
| CXCL12 | -0.01 | cytokine | | 0.762 |
| RETN | -0.117 | other | | -1.044 |
| APEX1 | -0.106 | enzyme | | 0 |
| ITCH | -0.081 | enzyme | Activated | 2.2 |
| NOS1 | -0.401 | enzyme | Inhibited | -2.207 |
| ADAMTS12 | -0.563 | peptidase | Activated | 2.213 |
| BIRC3 | -0.097 | enzyme | | |
| RC3H2 | -0.114 | enzyme | | |
| ELOVL2 | 0.185 | enzyme | | |
| SRC (family) | | group | | |
| NR2E3 | 0 | ligand-dependent nuclear receptor | | |
| NR0B1 | 0 | ligand-dependent nuclear receptor | | |
| TYRO3 | -0.384 | kinase | | |

| | | | | |
|---|---|---|---|---|
| mir-124 | 0 | microRNA | | |
| RAMP1 | -0.042 | G-protein coupled receptor | | |
| TACR1 | -0.285 | G-protein coupled receptor | | |
| FOSL2 | -0.324 | transcription regulator | | |
| NDUFS4 | 0.141 | enzyme | | |
| THRB | -0.028 | ligand-dependent nuclear receptor | | 0.555 |
| CD200R1 | -0.293 | transmembrane receptor | | 1 |
| HES3 | 0 | transcription regulator | | -0.378 |
| JAK1 | -0.077 | kinase | | |
| RAG2 | 0 | enzyme | | |
| CSF2 | -0.113 | cytokine | Inhibited | -3.925 |
| INSIG1 | -0.296 | other | Activated | 3.464 |
| LIF | -0.364 | cytokine | | -0.632 |
| CAV1 | -0.029 | transmembrane receptor | | 0.392 |
| LEF1 | -0.542 | transcription regulator | | 0.525 |
| FCGR2B | -0.682 | transmembrane receptor | | -1.172 |
| Trp53cor1 | -1.017 | other | | -1.477 |
| ARRB1 | -0.217 | transcription regulator | | -0.686 |
| SQSTM1 | 0.104 | transcription regulator | | 0.805 |
| BAX | -0.055 | transporter | | -1.238 |
| PRDM16 | -0.258 | transcription regulator | | 0.655 |
| CFB | -0.328 | peptidase | | -1.131 |
| CSF3 | 0 | cytokine | | |
| B2M | -0.317 | transmembrane receptor | | -0.545 |
| Cux1 | -0.082 | transcription regulator | | |
| CLU | 0.1 | other | | 1.746 |
| IL7 | -0.595 | cytokine | Inhibited | -2.449 |
| SOX2 | 0.313 | transcription regulator | | -1.24 |
| KDM4D | 1.196 | enzyme | | |
| PCK1 | 0.604 | kinase | | |
| MMP12 | -0.827 | peptidase | | |
| MBD2 | -0.145 | transcription regulator | | |
| CPEB1 | 0.043 | translation regulator | | |
| PAX2 | 0.521 | transcription regulator | | |
| NRP1 | 0.034 | transmembrane receptor | | |
| HAX1 | -0.039 | other | | |
| Flg | | other | | |
| NPC2 | -0.037 | transporter | | |
| TBXT | 3.459 | transcription regulator | | |
| NLRP1 | -0.311 | enzyme | | |
| ST8SIA1 | 0.369 | enzyme | | |
| Akt | | group | | -1.796 |
| UBR5 | -0.152 | enzyme | | -1.915 |
| SMARCA4 | -0.117 | transcription regulator | | -0.728 |
| PYCARD | -0.558 | transcription regulator | | -1.413 |
| CPE | 0.253 | peptidase | | |
| IL21R | -1.164 | transmembrane receptor | | -0.648 |
| JAK3 | -0.219 | kinase | Activated | 2.22 |
| MBTD1 | 0.077 | other | | -1.237 |
| IL21 | 2.939 | cytokine | | 0.465 |
| GNB2 | -0.039 | enzyme | | |
| PIN1 | -0.076 | enzyme | | 0 |
| GNB1 | -0.126 | enzyme | | |
| OGT | -0.12 | enzyme | Activated | 2.722 |
| BSCL2 | -0.167 | other | | |
| Hdac | | group | | 0.248 |
| Igm | | complex | | |
| IER3IP1 | 0.031 | other | | -1.067 |
| WWTR1 | -0.064 | transcription regulator | | -0.152 |
| PRKCA | -0.146 | kinase | | -1.977 |
| GJA1 | -0.01 | transporter | | -1 |
| SFRP4 | -0.301 | transmembrane receptor | | 1.131 |
| FGF2 | -0.232 | growth factor | | 0 |
| TYK2 | -0.394 | kinase | | -0.633 |
| CD3E | -0.978 | transmembrane receptor | | -1.953 |
| GNAQ | -0.03 | enzyme | | -1.134 |
| RXRB | -0.084 | ligand-dependent nuclear receptor | | 0.832 |
| Ck2 alpha | | group | | |
| Angiotensin II receptor type 1 | | group | | |
| ATP6AP2 | -0.021 | transporter | | |
| SLC5A1 | -2.939 | transporter | | |
| USP25 | -0.152 | peptidase | | |
| GALNT3 | -0.748 | enzyme | | |

| | | | | |
|---|---|---|---|---|
| RPE65 | 0 | enzyme | | |
| RAP1B | -0.196 | enzyme | | |
| CD244 | | transmembrane receptor | | |
| PPP4C | -0.109 | phosphatase | | |
| HOXB9 | -0.07 | transcription regulator | | |
| POMC | 0.056 | other | | |
| HOXC9 | -0.263 | transcription regulator | | |
| IL9R | -1.488 | transmembrane receptor | | |
| HSPA2 | -0.251 | other | | |
| DHX9 | -0.345 | enzyme | | |
| GNAI1 | -0.028 | enzyme | | |
| NCR1 | -1.073 | transmembrane receptor | | |
| KIDINS220 | -0.18 | transcription regulator | | |
| TRIM8 | -0.167 | enzyme | | |
| ETV6 | -0.264 | transcription regulator | | |
| RNF41 | -0.112 | enzyme | | |
| CD163 | -0.48 | transmembrane receptor | | |
| ADAM8 | -0.434 | peptidase | | |
| CCN3 | | growth factor | | |
| HOXD9 | -0.054 | transcription regulator | | |
| ACOD1 | | enzyme | | |
| ARL5B | -0.336 | enzyme | | |
| UBASH3A | -0.497 | enzyme | | |
| DSPP | 0 | other | | |
| CD247 | 0.468 | transmembrane receptor | | |
| PTGER3 | 0.084 | G-protein coupled receptor | | |
| TKT | -0.225 | enzyme | | |
| CREG1 | 0.014 | transcription regulator | | |
| NFKBIE | -0.429 | transcription regulator | | |
| CHRD | -0.112 | other | | |
| CCN1 | | other | | |
| PPP1R15A | -0.175 | other | | |
| Muc1 | -0.433 | transmembrane receptor | | |
| NDP | 1.402 | growth factor | | |
| Clec2d (includes others) | -0.598 | transmembrane receptor | | |
| MARCHF1 | | enzyme | | |
| ZBTB20 | -0.021 | transcription regulator | | -0.309 |
| MEF2A | -0.034 | transcription regulator | | -1.948 |
| CLEC4A | -0.211 | transmembrane receptor | | 1.219 |
| ATF3 | -0.372 | transcription regulator | Activated | 2.365 |
| IGF1 | -0.293 | growth factor | | 0.378 |
| IL6ST | -0.163 | transmembrane receptor | | 0.218 |
| SAMSN1 | -0.703 | other | Inhibited | -2.714 |
| RARG | -0.333 | ligand-dependent nuclear receptor | | -0.707 |
| TCIRG1 | -0.284 | enzyme | | |
| ARV1 | -0.218 | transporter | | |
| MFN2 | -0.191 | enzyme | | |
| IL13RA2 | 0.251 | transmembrane receptor | | |
| FERMT2 | -0.077 | other | | |
| PLA2G4A | -0.374 | enzyme | | |
| NR2F1 | -0.364 | ligand-dependent nuclear receptor | | |
| EPHA2 | -0.129 | kinase | | |
| GATA4 | 0 | transcription regulator | | -0.155 |
| MGLL | 0.085 | enzyme | | -1.981 |
| IL25 | 0.63 | cytokine | | -1.969 |
| USP19 | -0.083 | peptidase | | 1.988 |
| Nr1h | | group | Activated | 3.5 |
| PSMB11 | -2.115 | peptidase | Activated | 2.138 |
| PRKN | | enzyme | | 1.976 |
| PNPLA2 | 0.046 | enzyme | Activated | 2.964 |
| FOXA1 | 0 | transcription regulator | Activated | 2.433 |
| MAPK3 | 0.005 | kinase | Inhibited | -2.619 |
| ZC3H12A | -0.503 | enzyme | | 1.753 |
| NLRP12 | 0.684 | other | | 1.982 |
| BACH2 | 0.055 | transcription regulator | | 1.893 |
| DMP1 | 0.29 | other | Activated | 2.227 |
| RIGI | | enzyme | | -1.947 |
| C1QA | -0.61 | other | | -1.969 |
| CD36 | 0.075 | transmembrane receptor | | -1.981 |
| PRKAA1 | -0.172 | kinase | | 1.849 |
| CYLD | -0.255 | transcription regulator | Inhibited | -2.2 |
| S1PR4 | -0.141 | G-protein coupled receptor | | -1.89 |
| SGPP2 | -0.5 | phosphatase | Inhibited | -2 |

| | | | | |
|---|---|---|---|---|
| SPIB | -1.115 | transcription regulator | | -1.977 |
| GATA1 | 0.746 | transcription regulator | Inhibited | -2.578 |
| BCR (complex) | | complex | Inhibited | -2.299 |
| WNT4 | -0.37 | cytokine | | 1.98 |
| PTPN11 | -0.187 | phosphatase | Activated | 2.178 |
| KLF6 | 0.004 | transcription regulator | Inhibited | -3.361 |
| Hbb-b1 | | transporter | Inhibited | -2.985 |
| PTK2 | -0.104 | kinase | | -1.953 |
| HOXC8 | -0.252 | transcription regulator | | -1.964 |
| SNAI1 | -0.41 | transcription regulator | Activated | 2 |
| FKBP10 | -0.155 | enzyme | Activated | 2.646 |
| ADAM17 | -0.238 | peptidase | Inhibited | -2.917 |
| SOST | 0.143 | other | | -1.98 |
| FST | -0.109 | other | Inhibited | -2.219 |
| NLRX1 | 0.029 | other | Activated | 2.376 |
| ADAMTS18 | -0.542 | peptidase | Inhibited | -2 |
| KLF4 | -0.043 | transcription regulator | Activated | 2.366 |
| LHCGR | -0.521 | G-protein coupled receptor | | 1.932 |
| SMAD3 | -0.373 | transcription regulator | Inhibited | -2.051 |
| GPBAR1 | 1.887 | G-protein coupled receptor | | 1.97 |
| PRMT1 | -0.011 | enzyme | | -1.934 |
| CYP1A2 | 0 | enzyme | Inhibited | -2.236 |
| NRIP1 | -0.177 | transcription regulator | | -1.99 |
| PRKCB | -0.697 | kinase | | -1.981 |
| DRD2 | 0 | G-protein coupled receptor | Inhibited | -2.646 |
| NCOA5 | -0.161 | other | Activated | 2.219 |
| SIRT6 | -0.169 | enzyme | | 1.987 |
| TCF7L2 | 0.137 | transcription regulator | Inhibited | -3.147 |
| TYROBP | -0.181 | transmembrane receptor | | -1.91 |
| IPMK | -0.242 | kinase | Inhibited | -2.219 |
| MTORC1 | | complex | | -1.98 |
| METTL3 | -0.189 | enzyme | | 1.912 |
| CDKN1A | -0.148 | kinase | Activated | 2.4 |
| APOE | -0.022 | transporter | Activated | 2.366 |
| CREB1 | -0.051 | transcription regulator | | -1.949 |
| IL2RG | -0.528 | transmembrane receptor | Inhibited | -2.236 |
| CYP1A1 | 0.471 | enzyme | Inhibited | -2.236 |
| FOXO3 | -0.145 | transcription regulator | Activated | 2.352 |
| PDX1 | 0 | transcription regulator | | -1.96 |
| HMG20A | -0.132 | transcription regulator | Inhibited | -2.236 |
| EIF2AK3 | -0.26 | kinase | | -1.816 |
| CEBPB | 0.006 | transcription regulator | | -1.984 |
| INHA | 0.144 | growth factor | Activated | 2.219 |
| CD38 | -0.046 | enzyme | | -1.857 |
| ERN1 | -0.546 | kinase | Inhibited | -2.408 |
| RGS4 | -0.025 | enzyme | Activated | 2 |
| BMAL1 | -0.416 | transcription regulator | | 1.94 |
| MYB | 0.229 | transcription regulator | | -1.982 |
