## Supplementary material for "Scalable Generation of Universal hiPSC-Derived Vascular Progenitor Cells for Safe and Sustained Revascularization in Chronic Limb-Threatening Ischemia": Supplemental Table 3_pathways.pdf

**Supplemental Table 3. Ingenuity Pathway Analysis: Canoncial pathways**

| Ingenuity Canonical Pathways | -log(p-value) | Ratio | z-score |
|---|---|---|---|
| Granulocyte Adhesion and Diapedesis | 18.4 | 0.297 | #NUM! |
| Agranulocyte Adhesion and Diapedesis | 13.5 | 0.245 | #NUM! |
| S100 Family Signaling Pathway | 9.37 | 0.132 | -2.888 |
| Airway Pathology in Chronic Obstructive Pulmonary Disease | 8.89 | 0.257 | #NUM! |
| Pathogen Induced Cytokine Storm Signaling Pathway | 7.71 | 0.159 | -3.96 |
| Role of Hypercytokinemia/hyperchemokinemia in the Pathogenesis of Influenza | 6.45 | 0.269 | -3.3 |
| Phagosome Formation | 5.92 | 0.119 | -3.212 |
| Differential Regulation of Cytokine Production in Macrophages and T Helper Cells by IL-17A and IL-17F | 5.71 | 0.533 | -2.828 |
| CREB Signaling in Neurons | 5.55 | 0.12 | -2.425 |
| Neuroprotective Role of THOP1 in Alzheimer's Disease | 5.36 | 0.2 | -0.688 |
| G-Protein Coupled Receptor Signaling | 5.3 | 0.115 | -2.393 |
| Differential Regulation of Cytokine Production in Intestinal Epithelial Cells by IL-17A and IL-17F | 5.19 | 0.471 | -2.828 |
| FXR/RXR Activation | 5.03 | 0.191 | #NUM! |
| Atherosclerosis Signaling | 4.85 | 0.186 | #NUM! |
| LXR/RXR Activation | 4.69 | 0.188 | 0 |
| Phototransduction Pathway | 4.66 | 0.26 | #NUM! |
| Hepatic Cholestasis | 4.44 | 0.156 | #NUM! |
| Breast Cancer Regulation by Stathmin1 | 4.01 | 0.111 | -2.54 |
| Role of Pattern Recognition Receptors in Recognition of Bacteria and Viruses | 3.98 | 0.161 | -1.667 |
| Role of Cytokines in Mediating Communication between Immune Cells | 3.87 | 0.27 | #NUM! |
| TREM1 Signaling | 3.56 | 0.197 | -2.673 |
| Airway Inflammation in Asthma | 3.38 | 0.286 | #NUM! |
| Bile Acid Biosynthesis, Neutral Pathway | 3.19 | 0.353 | -1 |
| Inhibition of Matrix Metalloproteases | 3.16 | 0.243 | 1 |
| Tumor Microenvironment Pathway | 3.11 | 0.138 | -3.674 |
| SPINK1 Pancreatic Cancer Pathway | 3.04 | 0.204 | -0.905 |
| Role Of Chondrocytes In Rheumatoid Arthritis Signaling Pathway | 2.99 | 0.146 | -3.13 |
| IL-17 Signaling | 2.99 | 0.14 | -2.132 |
| Neuroinflammation Signaling Pathway | 2.94 | 0.119 | -2.268 |
| Glutamate Receptor Signaling | 2.87 | 0.185 | #NUM! |
| Eicosanoid Signaling | 2.87 | 0.185 | -1.134 |
| NOD1/2 Signaling Pathway | 2.77 | 0.135 | -2.558 |
| Histidine Degradation VI | 2.74 | 0.259 | #NUM! |
| Multiple Sclerosis Signaling Pathway | 2.72 | 0.128 | -1.8 |
| Coagulation System | 2.68 | 0.229 | -0.707 |
| Protein Citrullination | 2.54 | 0.6 | #NUM! |
| Embryonic Stem Cell Differentiation into Cardiac Lineages | 2.49 | 0.4 | #NUM! |
| Intrinsic Prothrombin Activation Pathway | 2.37 | 0.205 | -1.414 |
| Retinoate Biosynthesis I | 2.37 | 0.205 | -0.447 |
| LPS/IL-1 Mediated Inhibition of RXR Function | 2.35 | 0.117 | 0.277 |
| Gustation Pathway | 2.08 | 0.118 | -0.426 |
| Crosstalk between Dendritic Cells and Natural Killer Cells | 2.07 | 0.155 | -3.162 |
| Role Of Osteoblasts In Rheumatoid Arthritis Signaling Pathway | 2.03 | 0.112 | -2.353 |
| Immunogenic Cell Death Signaling Pathway | 2.02 | 0.146 | -1.732 |
| Bladder Cancer Signaling | 2 | 0.133 | -1.134 |
| Role of MAPK Signaling in Inhibiting the Pathogenesis of Influenza | 1.99 | 0.151 | -1.265 |
| Pyroptosis Signaling Pathway | 1.98 | 0.145 | -2.309 |
| MSP-RON Signaling Pathway | 1.94 | 0.164 | #NUM! |
| Glycine Biosynthesis III | 1.87 | 0.667 | #NUM! |
| Methionine Salvage II (Mammalian) | 1.87 | 0.667 | #NUM! |
| Neurovascular Coupling Signaling Pathway | 1.85 | 0.11 | -0.816 |
| FAK Signaling | 1.84 | 0.0883 | -3.628 |
| PXR/RXR Activation | 1.7 | 0.15 | 1.667 |
| cAMP-mediated signaling | 1.68 | 0.106 | -2.132 |
| Macrophage Classical Activation Signaling Pathway | 1.67 | 0.115 | -1.886 |
| CDX Gastrointestinal Cancer Signaling Pathway | 1.65 | 0.109 | 0.655 |
| Wound Healing Signaling Pathway | 1.65 | 0.105 | -2.2 |
| Role of IL-17F in Allergic Inflammatory Airway Diseases | 1.65 | 0.167 | -2.646 |
| Maturity Onset Diabetes of Young (MODY) Signaling | 1.62 | 0.139 | #NUM! |
| Retinol Biosynthesis | 1.6 | 0.163 | -0.816 |
| α-tocopherol Degradation | 1.58 | 0.3 | #NUM! |
| Primary Immunodeficiency Signaling | 1.55 | 0.159 | #NUM! |
| Pregnenolone Biosynthesis | 1.54 | 0.192 | #NUM! |
| Leukocyte Extravasation Signaling | 1.51 | 0.107 | -1.387 |
| Hepatic Fibrosis / Hepatic Stellate Cell Activation | 1.51 | 0.107 | #NUM! |
| Glucocorticoid Receptor Signaling | 1.49 | 0.09 | #NUM! |
| Role of Lipids/Lipid Rafts in the Pathogenesis of Influenza | 1.46 | 0.273 | #NUM! |
| Cardiac Hypertrophy Signaling (Enhanced) | 1.32 | 0.0878 | -3.244 |
| Acute Phase Response Signaling | 1.31 | 0.104 | -1.265 |

| | | | |
|---|---|---|---|
| Ubiquinol-10 Biosynthesis (Eukaryotic) | 1.3 | 0.167 | #NUM! |
| Trehalose Degradation II (Trehalase) | 1.24 | 0.333 | #NUM! |
| Glycine Cleavage Complex | 1.24 | 0.333 | #NUM! |
| Serotonin Receptor Signaling | 1.24 | 0.146 | #NUM! |
| Erythropoietin Signaling Pathway | 1.21 | 0.102 | 0.728 |
| Methylglyoxal Degradation III | 1.19 | 0.156 | #NUM! |
| Leukotriene Biosynthesis | 1.18 | 0.214 | #NUM! |
| Melatonin Degradation III | 1.17 | 1 | #NUM! |
| Triacylglycerol Degradation | 1.13 | 0.13 | -0.447 |
| Aspartate Degradation II | 1.11 | 0.286 | #NUM! |
| Role of Osteoblasts, Osteoclasts and Chondrocytes in Rheumatoid Arthritis | 1.11 | 0.095 | #NUM! |
| Role of JAK family kinases in IL-6-type Cytokine Signaling | 1.08 | 0.115 | -1 |
| Gαi Signaling | 1.05 | 0.101 | -1.732 |
| Role of MAPK Signaling in the Pathogenesis of Influenza | 1.05 | 0.114 | #NUM! |
| Extrinsic Prothrombin Activation Pathway | 1.04 | 0.188 | #NUM! |
| Oxidative Ethanol Degradation III | 1 | 0.128 | #NUM! |
| Histidine Degradation III | 1 | 0.25 | #NUM! |
| Synaptic Long Term Depression | 0.986 | 0.0942 | -1 |
| Stearate Biosynthesis I (Animals) | 0.984 | 0.114 | #NUM! |
| Macrophage Alternative Activation Signaling Pathway | 0.977 | 0.095 | -0.728 |
| Androgen Biosynthesis | 0.976 | 0.176 | #NUM! |
| Nicotine Degradation III | 0.971 | 0.125 | 0.816 |
| HMGB1 Signaling | 0.962 | 0.0968 | -1.732 |
| IL-13 Signaling Pathway | 0.945 | 0.105 | -2.53 |
| IL-33 Signaling Pathway | 0.936 | 0.0947 | -2.5 |
| Heparan Sulfate Biosynthesis (Late Stages) | 0.931 | 0.111 | -0.816 |
| Thyroid Hormone Metabolism II (via Conjugation and/or Degradation) | 0.919 | 0.143 | 2 |
| Role of IL-17A in Psoriasis | 0.914 | 0.222 | #NUM! |
| Role of Macrophages, Fibroblasts and Endothelial Cells in Rheumatoid Arthritis | 0.89 | 0.086 | #NUM! |
| Thyroid Hormone Biosynthesis | 0.88 | 0.5 | #NUM! |
| Threonine Degradation II | 0.88 | 0.5 | #NUM! |
| Phospholipases | 0.863 | 0.111 | -0.816 |
| Superpathway of Melatonin Degradation | 0.848 | 0.115 | 1.633 |
| Glycine Betaine Degradation | 0.836 | 0.2 | #NUM! |
| p38 MAPK Signaling | 0.808 | 0.0965 | -0.905 |
| Glucocorticoid Biosynthesis | 0.768 | 0.182 | #NUM! |
| Heparan Sulfate Biosynthesis | 0.768 | 0.101 | -0.816 |
| tRNA Splicing | 0.746 | 0.114 | -1.342 |
| Adrenomedullin signaling pathway | 0.745 | 0.0872 | -2.324 |
| Nicotine Degradation II | 0.741 | 0.107 | 0.816 |
| SPINK1 General Cancer Pathway | 0.741 | 0.107 | 0.816 |
| The Visual Cycle | 0.728 | 0.136 | #NUM! |
| MSP-RON Signaling In Macrophages Pathway | 0.727 | 0.0943 | 0.632 |
| Uracil Degradation II (Reductive) | 0.718 | 0.333 | #NUM! |
| Retinoate Biosynthesis II | 0.718 | 0.333 | #NUM! |
| Thymine Degradation | 0.718 | 0.333 | #NUM! |
| Glutamate Degradation II | 0.718 | 0.333 | #NUM! |
| Aspartate Biosynthesis | 0.718 | 0.333 | #NUM! |
| Antioxidant Action of Vitamin C | 0.71 | 0.0935 | 0.707 |
| γ-glutamyl Cycle | 0.708 | 0.167 | #NUM! |
| IL-23 Signaling Pathway | 0.691 | 0.109 | -1.342 |
| Oxytocin In Brain Signaling Pathway | 0.68 | 0.0856 | 0 |
| Osteoarthritis Pathway | 0.678 | 0.0837 | -1.807 |
| Corticotropin Releasing Hormone Signaling | 0.674 | 0.0878 | -1.265 |
| STAT3 Pathway | 0.674 | 0.0889 | -1.342 |
| Oxytocin In Spinal Neurons Signaling Pathway | 0.668 | 0.114 | -2 |
| Regulation Of The Epithelial Mesenchymal Transition By Growth Factors Pathway | 0.667 | 0.0851 | -1.667 |
| Melatonin Degradation I | 0.666 | 0.106 | 1.342 |
| Role of PKR in Interferon Induction and Antiviral Response | 0.659 | 0.0894 | -1.414 |
| GPCR-Mediated Integration of Enteroendocrine Signaling Exemplified by an L Cell | 0.657 | 0.0972 | -0.378 |
| Relaxin Signaling | 0.634 | 0.0861 | -2.236 |
| FGF Signaling | 0.632 | 0.093 | -0.707 |
| Catecholamine Biosynthesis | 0.608 | 0.25 | #NUM! |
| Eumelanin Biosynthesis | 0.608 | 0.25 | #NUM! |
| Arginine Degradation I (Arginase Pathway) | 0.608 | 0.25 | #NUM! |
| L-cysteine Degradation I | 0.608 | 0.25 | #NUM! |
| Acyl-CoA Hydrolysis | 0.606 | 0.143 | #NUM! |
| Phenylalanine Degradation IV (Mammalian, via Side Chain) | 0.606 | 0.143 | #NUM! |
| GABA Receptor Signaling | 0.588 | 0.0859 | #NUM! |
| Pulmonary Healing Signaling Pathway | 0.587 | 0.0821 | -1.5 |
| Gluconeogenesis I | 0.584 | 0.115 | #NUM! |

| | | | |
|---|---|---|---|
| Adenosine Nucleotides Degradation II | 0.563 | 0.133 | #NUM! |
| Transcriptional Regulatory Network in Embryonic Stem Cells | 0.551 | 0.0962 | #NUM! |
| Role of WNT/GSK-3β Signaling in the Pathogenesis of Influenza | 0.548 | 0.0923 | #NUM! |
| Acetone Degradation I (to Methylglyoxal) | 0.536 | 0.1 | #NUM! |
| Xenobiotic Metabolism AHR Signaling Pathway | 0.531 | 0.0886 | -1.134 |
| Role of IL-17A in Arthritis | 0.531 | 0.0943 | #NUM! |
| Serotonin Degradation | 0.531 | 0.0943 | 2.236 |
| Pentose Phosphate Pathway (Oxidative Branch) | 0.526 | 0.2 | #NUM! |
| Tetrapyrrole Biosynthesis II | 0.526 | 0.2 | #NUM! |
| Lysine Degradation II | 0.526 | 0.2 | #NUM! |
| Lysine Degradation V | 0.526 | 0.2 | #NUM! |
| Tyrosine Degradation I | 0.526 | 0.2 | #NUM! |
| Chondroitin Sulfate Degradation (Metazoa) | 0.524 | 0.125 | #NUM! |
| PD-1, PD-L1 cancer immunotherapy pathway | 0.52 | 0.086 | 0 |
| Dermatan Sulfate Biosynthesis (Late Stages) | 0.514 | 0.0976 | -1 |
| Activation of IRF by Cytosolic Pattern Recognition Receptors | 0.511 | 0.0926 | -2.236 |
| Endocannabinoid Neuronal Synapse Pathway | 0.506 | 0.0811 | -0.302 |
| Role of JAK1 and JAK3 in γc Cytokine Signaling | 0.495 | 0.0882 | #NUM! |
| Dermatan Sulfate Degradation (Metazoa) | 0.488 | 0.118 | #NUM! |
| Synaptogenesis Signaling Pathway | 0.474 | 0.0757 | -1.706 |
| Pathogenesis of Multiple Sclerosis | 0.461 | 0.167 | #NUM! |
| Urea Cycle | 0.461 | 0.167 | #NUM! |
| Arginine Degradation VI (Arginase 2 Pathway) | 0.461 | 0.167 | #NUM! |
| Adenine and Adenosine Salvage III | 0.461 | 0.167 | #NUM! |
| Tryptophan Degradation to 2-amino-3-carboxymuconate Semialdehyde | 0.461 | 0.167 | #NUM! |
| Ceramide Degradation | 0.461 | 0.167 | #NUM! |
| IL-6 Signaling | 0.456 | 0.08 | -1.265 |
| Purine Nucleotides Degradation II (Aerobic) | 0.455 | 0.111 | #NUM! |
| Retinoic acid Mediated Apoptosis Signaling | 0.451 | 0.0909 | #NUM! |
| Chondroitin Sulfate Biosynthesis (Late Stages) | 0.451 | 0.0909 | -1 |
| Role of Tissue Factor in Cancer | 0.447 | 0.0804 | #NUM! |
| Neutrophil Extracellular Trap Signaling Pathway | 0.441 | 0.074 | -0.962 |
| Th1 and Th2 Activation Pathway | 0.432 | 0.0774 | #NUM! |
| WNT/β-catenin Signaling | 0.432 | 0.0769 | 0.632 |
| Inflammasome pathway | 0.425 | 0.105 | #NUM! |
| Cellular Effects of Sildenafil (Viagra) | 0.422 | 0.0775 | #NUM! |
| Role Of Osteoclasts In Rheumatoid Arthritis Signaling Pathway | 0.415 | 0.0738 | -1.414 |
| Ceramide Biosynthesis | 0.408 | 0.143 | #NUM! |
| Purine Ribonucleosides Degradation to Ribose-1-phosphate | 0.408 | 0.143 | #NUM! |
| Salvage Pathways of Pyrimidine Deoxyribonucleotides | 0.408 | 0.143 | #NUM! |
| Sertoli Cell-Sertoli Cell Junction Signaling | 0.406 | 0.075 | #NUM! |
| Endothelin-1 Signaling | 0.405 | 0.0753 | 0.277 |
| Coronavirus Pathogenesis Pathway | 0.405 | 0.0753 | -0.535 |
| Toll-like Receptor Signaling | 0.404 | 0.0811 | -1 |
| Complement System | 0.404 | 0.0909 | #NUM! |
| GPCR-Mediated Nutrient Sensing in Enteroendocrine Cells | 0.402 | 0.0776 | -1 |
| MIF-mediated Glucocorticoid Regulation | 0.384 | 0.0882 | #NUM! |
| Chemokine Signaling | 0.378 | 0.0789 | -0.816 |
| IL-17A Signaling in Fibroblasts | 0.365 | 0.0857 | #NUM! |
| Sphingosine and Sphingosine-1-phosphate Metabolism | 0.364 | 0.125 | #NUM! |
| IL-12 Signaling and Production in Macrophages | 0.359 | 0.0728 | -0.258 |
| Gap Junction Signaling | 0.349 | 0.0725 | #NUM! |
| Tumoricidal Function of Hepatic Natural Killer Cells | 0.348 | 0.0909 | #NUM! |
| Th2 Pathway | 0.341 | 0.0738 | -2.121 |
| Th1 Pathway | 0.328 | 0.0734 | -1.89 |
| Role of MAPK Signaling in Promoting the Pathogenesis of Influenza | 0.328 | 0.0734 | -0.378 |
| IL-10 Signaling | 0.327 | 0.0725 | 1.265 |
| Citrulline Biosynthesis | 0.327 | 0.111 | #NUM! |
| Heme Biosynthesis II | 0.327 | 0.111 | #NUM! |
| Ketolysis | 0.327 | 0.111 | #NUM! |
| IL-17A Signaling in Gastric Cells | 0.326 | 0.087 | #NUM! |
| Chondroitin Sulfate Biosynthesis | 0.32 | 0.0769 | -1 |
| Docosahexaenoic Acid (DHA) Signaling | 0.313 | 0.0789 | #NUM! |
| IL-22 Signaling | 0.305 | 0.0833 | #NUM! |
| Cardiac β-adrenergic Signaling | 0.304 | 0.0706 | -1.633 |
| Sperm Motility | 0.299 | 0.0697 | -0.707 |
| Superpathway of Methionine Degradation | 0.298 | 0.0769 | #NUM! |
| Prostanoid Biosynthesis | 0.295 | 0.1 | #NUM! |
| Sucrose Degradation V (Mammalian) | 0.295 | 0.1 | #NUM! |
| Calcium Transport I | 0.295 | 0.1 | #NUM! |
| GDP-glucose Biosynthesis | 0.295 | 0.1 | #NUM! |

| | | | |
|---|---|---|---|
| Mineralocorticoid Biosynthesis | 0.295 | 0.1 | #NUM! |
| Necroptosis Signaling Pathway | 0.294 | 0.0704 | -0.632 |
| Role of PI3K/AKT Signaling in the Pathogenesis of Influenza | 0.293 | 0.0741 | 0 |
| Dermatan Sulfate Biosynthesis | 0.293 | 0.0741 | -1 |
| HOTAIR Regulatory Pathway | 0.292 | 0.0701 | -0.905 |
| CCR3 Signaling in Eosinophils | 0.289 | 0.0703 | 0.447 |
| Glycolysis I | 0.286 | 0.08 | #NUM! |
| Estrogen Biosynthesis | 0.283 | 0.075 | #NUM! |
| D-myo-inositol (1,4,5)-Trisphosphate Biosynthesis | 0.269 | 0.0769 | #NUM! |
| Apelin Liver Signaling Pathway | 0.269 | 0.0769 | #NUM! |
| Hematopoiesis from Multipotent Stem Cells | 0.267 | 0.0909 | #NUM! |
| Glucose and Glucose-1-phosphate Degradation | 0.267 | 0.0909 | #NUM! |
| NAD biosynthesis II (from tryptophan) | 0.267 | 0.0909 | #NUM! |
| Pentose Phosphate Pathway | 0.267 | 0.0909 | #NUM! |
| SNARE Signaling Pathway | 0.266 | 0.0687 | -1 |
| Apelin Adipocyte Signaling Pathway | 0.259 | 0.069 | 0.447 |
| MIF Regulation of Innate Immunity | 0.256 | 0.0714 | #NUM! |
| Oncostatin M Signaling | 0.256 | 0.0714 | #NUM! |
| UDP-N-acetyl-D-galactosamine Biosynthesis II | 0.242 | 0.0833 | #NUM! |
| Antigen Presentation Pathway | 0.237 | 0.0714 | #NUM! |
| Sonic Hedgehog Signaling | 0.223 | 0.069 | #NUM! |
| Guanosine Nucleotides Degradation III | 0.221 | 0.0769 | #NUM! |
| Urate Biosynthesis/Inosine 5'-phosphate Degradation | 0.201 | 0.0714 | #NUM! |
| Dilated Cardiomyopathy Signaling Pathway | 0 | 0.0411 | -1 |
| NAD Signaling Pathway | 0 | 0.0282 | -2 |
| CSDE1 Signaling Pathway | 0 | 0.0588 | #NUM! |
| Oxytocin Signaling Pathway | 0 | 0.0549 | -0.258 |
| Pulmonary Fibrosis Idiopathic Signaling Pathway | 0 | 0.0629 | -2.065 |
| CLEAR Signaling Pathway | 0 | 0.0284 | 1.414 |
| ID1 Signaling Pathway | 0 | 0.0663 | -0.832 |
| MicroRNA Biogenesis Signaling Pathway | 0 | 0.017 | #NUM! |
| Ribonucleotide Reductase Signaling Pathway | 0 | 0.0311 | 0.447 |
| Natural Killer Cell Signaling | 0 | 0.0471 | -2.121 |
| Neuregulin Signaling | 0 | 0.0431 | #NUM! |
| Circadian Rhythm Signaling | 0 | 0.0646 | #NUM! |
| Synaptic Long Term Potentiation | 0 | 0.0385 | -0.447 |
| Axonal Guidance Signaling | 0 | 0.063 | #NUM! |
| Amyotrophic Lateral Sclerosis Signaling | 0 | 0.0619 | 1 |
| Fc Epsilon RI Signaling | 0 | 0.0531 | 1 |
| Actin Cytoskeleton Signaling | 0 | 0.0593 | #NUM! |
| Huntington's Disease Signaling | 0 | 0.0221 | 0.447 |
| Chaperone Mediated Autophagy Signaling Pathway | 0 | 0.0297 | 2.496 |
| Myelination Signaling Pathway | 0 | 0.0379 | -1.155 |
| NRF2-mediated Oxidative Stress Response | 0 | 0.0548 | #NUM! |
| PPARα/RXRα Activation | 0 | 0.0484 | 0.707 |
| Aryl Hydrocarbon Receptor Signaling | 0 | 0.0608 | -1.134 |
| p53 Signaling | 0 | 0.0319 | #NUM! |
| Mitochondrial Dysfunction | 0 | 0.0185 | 0.816 |
| VDR/RXR Activation | 0 | 0.0641 | 1 |
| Ceramide Signaling | 0 | 0.0112 | #NUM! |
| Tight Junction Signaling | 0 | 0.0578 | #NUM! |
| TR/RXR Activation | 0 | 0.0625 | #NUM! |
| Regulation of Actin-based Motility by Rho | 0 | 0.0463 | #NUM! |
| Role of BRCA1 in DNA Damage Response | 0 | 0.025 | #NUM! |
| RAR Activation | 0 | 0.0653 | #NUM! |
| 14-3-3-mediated Signaling | 0 | 0.024 | #NUM! |
| α-Adrenergic Signaling | 0 | 0.0472 | #NUM! |
| Caveolar-mediated Endocytosis Signaling | 0 | 0.0548 | #NUM! |
| Clathrin-mediated Endocytosis Signaling | 0 | 0.0667 | #NUM! |
| Fcγ Receptor-mediated Phagocytosis in Macrophages and Monocytes | 0 | 0.0326 | #NUM! |
| IL-8 Signaling | 0 | 0.0398 | -0.816 |
| Role of RIG1-like Receptors in Antiviral Innate Immunity | 0 | 0.0286 | #NUM! |
| Role of NFAT in Regulation of the Immune Response | 0 | 0.00775 | -1 |
| FcγRIIB Signaling in B Lymphocytes | 0 | 0.0058 | #NUM! |
| LPS-stimulated MAPK Signaling | 0 | 0.012 | #NUM! |
| CCR5 Signaling in Macrophages | 0 | 0.0291 | 0 |
| CD40 Signaling | 0 | 0.0152 | #NUM! |
| Calcium-induced T Lymphocyte Apoptosis | 0 | 0.00772 | #NUM! |
| Cytotoxic T Lymphocyte-mediated Apoptosis of Target Cells | 0 | 0.00855 | #NUM! |
| CD27 Signaling in Lymphocytes | 0 | 0.0357 | #NUM! |
| Lymphotoxin β Receptor Signaling | 0 | 0.0185 | #NUM! |

| | | | |
|---|---|---|---|
| fMLP Signaling in Neutrophils | 0 | 0.0469 | #NUM! |
| CXCR4 Signaling | 0 | 0.0366 | 0 |
| 4-1BB Signaling in T Lymphocytes | 0 | 0.0606 | #NUM! |
| Thrombopoietin Signaling | 0 | 0.0159 | #NUM! |
| CTLA4 Signaling in Cytotoxic T Lymphocytes | 0 | 0.0229 | 0.333 |
| Induction of Apoptosis by HIV1 | 0 | 0.0469 | #NUM! |
| B Cell Activating Factor Signaling | 0 | 0.0465 | #NUM! |
| IL-15 Production | 0 | 0.0513 | -1.633 |
| T Helper Cell Differentiation | 0 | 0.0368 | #NUM! |
| IL-9 Signaling | 0 | 0.0571 | #NUM! |
| CD28 Signaling in T Helper Cells | 0 | 0.00633 | #NUM! |
| IL-15 Signaling | 0 | 0.0118 | -1 |
| Virus Entry via Endocytic Pathways | 0 | 0.00885 | #NUM! |
| Dendritic Cell Maturation | 0 | 0.0348 | -3.051 |
| Mechanisms of Viral Exit from Host Cells | 0 | 0.0256 | #NUM! |
| Reelin Signaling in Neurons | 0 | 0.0296 | #NUM! |
| HIF1α Signaling | 0 | 0.0398 | -2.121 |
| Melatonin Signaling | 0 | 0.0435 | #NUM! |
| Neuropathic Pain Signaling In Dorsal Horn Neurons | 0 | 0.0505 | -1.342 |
| Factors Promoting Cardiogenesis in Vertebrates | 0 | 0.0473 | -0.378 |
| Agrin Interactions at Neuromuscular Junction | 0 | 0.0448 | #NUM! |
| Renin-Angiotensin Signaling | 0 | 0.0417 | -1 |
| Thrombin Signaling | 0 | 0.0459 | -0.816 |
| Cardiac Hypertrophy Signaling | 0 | 0.0438 | 0.378 |
| CDK5 Signaling | 0 | 0.0446 | -2.236 |
| ICOS-ICOSL Signaling in T Helper Cells | 0 | 0.013 | -1 |
| Molecular Mechanisms of Cancer | 0 | 0.0342 | #NUM! |
| Mitotic Roles of Polo-Like Kinase | 0 | 0.0308 | #NUM! |
| HGF Signaling | 0 | 0.031 | #NUM! |
| Role of CHK Proteins in Cell Cycle Checkpoint Control | 0 | 0.0357 | #NUM! |
| FLT3 Signaling in Hematopoietic Progenitor Cells | 0 | 0.0123 | #NUM! |
| Polyamine Regulation in Colon Cancer | 0 | 0.0345 | #NUM! |
| GNRH Signaling | 0 | 0.043 | -1 |
| Cholecystokinin/Gastrin-mediated Signaling | 0 | 0.0354 | -1 |
| Human Embryonic Stem Cell Pluripotency | 0 | 0.0352 | 0.378 |
| Melanocyte Development and Pigmentation Signaling | 0 | 0.0206 | #NUM! |
| DNA Methylation and Transcriptional Repression Signaling | 0 | 0.0278 | #NUM! |
| ATM Signaling | 0 | 0.0306 | #NUM! |
| Antiproliferative Role of Somatostatin Receptor 2 | 0 | 0.0519 | #NUM! |
| Androgen Signaling | 0 | 0.0368 | 0 |
| Role of OCT4 in Mammalian Embryonic Stem Cell Pluripotency | 0 | 0.0444 | #NUM! |
| Germ Cell-Sertoli Cell Junction Signaling | 0 | 0.0247 | #NUM! |
| Aldosterone Signaling in Epithelial Cells | 0 | 0.0595 | -1 |
| Role of NANOG in Mammalian Embryonic Stem Cell Pluripotency | 0 | 0.041 | #NUM! |
| Growth Hormone Signaling | 0 | 0.0147 | #NUM! |
| Prolactin Signaling | 0 | 0.0106 | #NUM! |
| Melanoma Signaling | 0 | 0.02 | #NUM! |
| Prostate Cancer Signaling | 0 | 0.0177 | #NUM! |
| Type I Diabetes Mellitus Signaling | 0 | 0.0261 | -1 |
| Small Cell Lung Cancer Signaling | 0 | 0.0105 | #NUM! |
| Basal Cell Carcinoma Signaling | 0 | 0.058 | #NUM! |
| Allograft Rejection Signaling | 0 | 0.0301 | #NUM! |
| Glioma Signaling | 0 | 0.0163 | #NUM! |
| Autoimmune Thyroid Disease Signaling | 0 | 0.0202 | #NUM! |
| Acute Myeloid Leukemia Signaling | 0 | 0.0111 | #NUM! |
| Thyroid Cancer Signaling | 0 | 0.039 | #NUM! |
| Graft-versus-Host Disease Signaling | 0 | 0.0247 | #NUM! |
| Type II Diabetes Mellitus Signaling | 0 | 0.0397 | #NUM! |
| Chronic Myeloid Leukemia Signaling | 0 | 0.0513 | 0 |
| Non-Small Cell Lung Cancer Signaling | 0 | 0.0108 | #NUM! |
| Production of Nitric Oxide and Reactive Oxygen Species in Macrophages | 0 | 0.0393 | -0.447 |
| Gα12/13 Signaling | 0 | 0.0538 | 1.134 |
| p70S6K Signaling | 0 | 0.0128 | -2.236 |
| ERK5 Signaling | 0 | 0.0411 | #NUM! |
| Colorectal Cancer Metastasis Signaling | 0 | 0.0682 | -2 |
| mTOR Signaling | 0 | 0.00495 | #NUM! |
| MYC Mediated Apoptosis Signaling | 0 | 0.0625 | #NUM! |
| Pancreatic Adenocarcinoma Signaling | 0 | 0.0163 | #NUM! |
| G Beta Gamma Signaling | 0 | 0.0472 | -1.633 |
| G Protein Signaling Mediated by Tubby | 0 | 0.0143 | -1 |
| Communication between Innate and Adaptive Immune Cells | 0 | 0.0258 | #NUM! |

| | | | |
|---|---|---|---|
| Sphingosine-1-phosphate Signaling | 0 | 0.0351 | #NUM! |
| Systemic Lupus Erythematosus Signaling | 0 | 0.0185 | #NUM! |
| CDC42 Signaling | 0 | 0.0223 | #NUM! |
| ILK Signaling | 0 | 0.0314 | 0 |
| EIF2 Signaling | 0 | 0.0144 | #NUM! |
| April Mediated Signaling | 0 | 0.0238 | #NUM! |
| AMPK Signaling | 0 | 0.0338 | #NUM! |
| PAK Signaling | 0 | 0.0439 | #NUM! |
| Hereditary Breast Cancer Signaling | 0 | 0.0216 | #NUM! |
| RAC Signaling | 0 | 0.0221 | #NUM! |
| RHOA Signaling | 0 | 0.0336 | 2 |
| Phospholipase C Signaling | 0 | 0.0123 | 1.342 |
| Ovarian Cancer Signaling | 0 | 0.0323 | #NUM! |
| HER-2 Signaling in Breast Cancer | 0 | 0.0228 | -1.342 |
| Altered T Cell and B Cell Signaling in Rheumatoid Arthritis | 0 | 0.0254 | #NUM! |
| Protein Kinase A Signaling | 0 | 0.0543 | -1.291 |
| Regulation of eIF4 and p70S6K Signaling | 0 | 0.0174 | #NUM! |
| Estrogen-Dependent Breast Cancer Signaling | 0 | 0.0253 | #NUM! |
| Leptin Signaling in Obesity | 0 | 0.0533 | #NUM! |
| Role of NFAT in Cardiac Hypertrophy | 0 | 0.0411 | 0 |
| Glioma Invasiveness Signaling | 0 | 0.0286 | #NUM! |
| B Cell Development | 0 | 0.00692 | #NUM! |
| IL-1 Signaling | 0 | 0.0532 | #NUM! |
| RANK Signaling in Osteoclasts | 0 | 0.0225 | #NUM! |
| Glioblastoma Multiforme Signaling | 0 | 0.0298 | 0.447 |
| Regulation of IL-2 Expression in Activated and Anergic T Lymphocytes | 0 | 0.00339 | #NUM! |
| Granzyme A Signaling | 0 | 0.0299 | #NUM! |
| NUR77 Signaling in T Lymphocytes | 0 | 0.0201 | #NUM! |
| PKCθ Signaling in T Lymphocytes | 0 | 0.00838 | #NUM! |
| TNFR1 Signaling | 0 | 0.02 | #NUM! |
| TNFR2 Signaling | 0 | 0.0312 | #NUM! |
| Antiproliferative Role of TOB in T Cell Signaling | 0 | 0.0042 | #NUM! |
| OX40 Signaling Pathway | 0 | 0.0149 | #NUM! |
| PI3K Signaling in B Lymphocytes | 0 | 0.015 | -1.342 |
| Inhibition of Angiogenesis by TSP1 | 0 | 0.0606 | #NUM! |
| P2Y Purigenic Receptor Signaling Pathway | 0 | 0.0305 | #NUM! |
| Cyclins and Cell Cycle Regulation | 0 | 0.0366 | #NUM! |
| Mismatch Repair in Eukaryotes | 0 | 0.0556 | #NUM! |
| IL-17A Signaling in Airway Cells | 0 | 0.0635 | -1 |
| Role of JAK2 in Hormone-like Cytokine Signaling | 0 | 0.0536 | #NUM! |
| Actin Nucleation by ARP-WASP Complex | 0 | 0.0227 | #NUM! |
| Dopamine-DARPP32 Feedback in cAMP Signaling | 0 | 0.0608 | -0.447 |
| NGF Signaling | 0 | 0.00847 | #NUM! |
| Paxillin Signaling | 0 | 0.0381 | #NUM! |
| Signaling by Rho Family GTPases | 0 | 0.0462 | 0 |
| RHOGDI Signaling | 0 | 0.0607 | 0 |
| Telomerase Signaling | 0 | 0.00962 | #NUM! |
| Mouse Embryonic Stem Cell Pluripotency | 0 | 0.0294 | #NUM! |
| Hematopoiesis from Pluripotent Stem Cells | 0 | 0.0249 | #NUM! |
| eNOS Signaling | 0 | 0.0537 | 0 |
| iNOS Signaling | 0 | 0.0435 | #NUM! |
| nNOS Signaling in Neurons | 0 | 0.0667 | #NUM! |
| nNOS Signaling in Skeletal Muscle Cells | 0 | 0.0638 | #NUM! |
| VEGF Family Ligand-Receptor Interactions | 0 | 0.0366 | #NUM! |
| Ephrin A Signaling | 0 | 0.0652 | #NUM! |
| Ephrin B Signaling | 0 | 0.0282 | #NUM! |
| ERBB Signaling | 0 | 0.044 | -1 |
| ERB2-ERBB3 Signaling | 0 | 0.0312 | #NUM! |
| ERBB4 Signaling | 0 | 0.0299 | #NUM! |
| Estrogen-mediated S-phase Entry | 0 | 0.0385 | #NUM! |
| GADD45 Signaling | 0 | 0.0678 | 1 |
| GDNF Family Ligand-Receptor Interactions | 0 | 0.0133 | #NUM! |
| Netrin Signaling | 0 | 0.0423 | #NUM! |
| Fatty Acid Activation | 0 | 0.0667 | #NUM! |
| Bupropion Degradation | 0 | 0.0455 | #NUM! |
| Glutathione Redox Reactions I | 0 | 0.04 | #NUM! |
| CDP-diacylglycerol Biosynthesis I | 0 | 0.04 | #NUM! |
| Superpathway of Citrulline Metabolism | 0 | 0.0667 | #NUM! |
| D-myo-inositol-5-phosphate Metabolism | 0 | 0.0269 | -2.236 |
| γ-linolenate Biosynthesis II (Animals) | 0 | 0.0556 | #NUM! |
| NAD Salvage Pathway II | 0 | 0.0435 | #NUM! |

| | | | |
|---|---|---|---|
| Pyridoxal 5'-phosphate Salvage Pathway | 0 | 0.0154 | #NUM! |
| Tryptophan Degradation X (Mammalian, via Tryptamine) | 0 | 0.04 | #NUM! |
| Glutathione-mediated Detoxification | 0 | 0.0312 | #NUM! |
| Phosphatidylglycerol Biosynthesis II (Non-plastidic) | 0 | 0.037 | #NUM! |
| D-myo-inositol (1,4,5,6)-Tetrakisphosphate Biosynthesis | 0 | 0.0235 | -2 |
| Superpathway of Inositol Phosphate Compounds | 0 | 0.0268 | -1.633 |
| Mitochondrial L-carnitine Shuttle Pathway | 0 | 0.0526 | #NUM! |
| D-myo-inositol (3,4,5,6)-tetrakisphosphate Biosynthesis | 0 | 0.0235 | -2 |
| TCA Cycle II (Eukaryotic) | 0 | 0.0455 | #NUM! |
| 3-phosphoinositide Degradation | 0 | 0.022 | -2 |
| 3-phosphoinositide Biosynthesis | 0 | 0.0255 | -1.342 |
| Tryptophan Degradation III (Eukaryotic) | 0 | 0.0385 | #NUM! |
| Triacylglycerol Biosynthesis | 0 | 0.0588 | #NUM! |
| Salvage Pathways of Pyrimidine Ribonucleotides | 0 | 0.0217 | #NUM! |
| Noradrenaline and Adrenaline Degradation | 0 | 0.0323 | #NUM! |
| Ethanol Degradation II | 0 | 0.0345 | #NUM! |
| Fatty Acid β-oxidation I | 0 | 0.0286 | #NUM! |
| DNA damage-induced 14-3-3σ Signaling | 0 | 0.0455 | #NUM! |
| Epithelial Adherens Junction Signaling | 0 | 0.00654 | #NUM! |
| Gαq Signaling | 0 | 0.0303 | #NUM! |
| Gαs Signaling | 0 | 0.065 | -1.342 |
| PEDF Signaling | 0 | 0.0241 | #NUM! |
| Regulation of Cellular Mechanics by Calpain Protease | 0 | 0.0568 | #NUM! |
| Remodeling of Epithelial Adherens Junctions | 0 | 0.0312 | #NUM! |
| Regulation of the Epithelial-Mesenchymal Transition Pathway | 0 | 0.0674 | #NUM! |
| Role of p14/p19ARF in Tumor Suppression | 0 | 0.037 | #NUM! |
| TEC Kinase Signaling | 0 | 0.0234 | #NUM! |
| UVA-Induced MAPK Signaling | 0 | 0.0206 | #NUM! |
| UVB-Induced MAPK Signaling | 0 | 0.0196 | #NUM! |
| UVC-Induced MAPK Signaling | 0 | 0.0196 | #NUM! |
| Oxidative Phosphorylation | 0 | 0.0098 | #NUM! |
| Adipogenesis pathway | 0 | 0.00741 | #NUM! |
| BER (Base Excision Repair) Pathway | 0 | 0.0233 | #NUM! |
| HIPPO signaling | 0 | 0.0119 | #NUM! |
| PCP (Planar Cell Polarity) Pathway | 0 | 0.0667 | #NUM! |
| Unfolded protein response | 0 | 0.0449 | #NUM! |
| WNT/Ca+ pathway | 0 | 0.0606 | -1 |
| Parkinson's Signaling | 0 | 0.0625 | #NUM! |
| Estrogen Receptor Signaling | 0 | 0.0375 | -1.732 |
| EGF Signaling | 0 | 0.0182 | #NUM! |
| Cell Cycle: G1/S Checkpoint Regulation | 0 | 0.0154 | #NUM! |
| ERK/MAPK Signaling | 0 | 0.024 | #NUM! |
| SAPK/JNK Signaling | 0 | 0.00641 | #NUM! |
| PI3K/AKT Signaling | 0 | 0.0556 | #NUM! |
| PTEN Signaling | 0 | 0.0268 | #NUM! |
| Nitric Oxide Signaling in the Cardiovascular System | 0 | 0.0435 | #NUM! |
| Protein Ubiquitination Pathway | 0 | 0.0265 | #NUM! |
| IL-2 Signaling | 0 | 0.0164 | #NUM! |
| Amyloid Processing | 0 | 0.04 | #NUM! |
| JAK/STAT Signaling | 0 | 0.0244 | #NUM! |
| Cell Cycle: G2/M DNA Damage Checkpoint Regulation | 0 | 0.06 | #NUM! |
| Xenobiotic Metabolism Signaling | 0 | 0.0597 | #NUM! |
| IL-4 Signaling | 0 | 0.0187 | -1.89 |
| B Cell Receptor Signaling | 0 | 0.0136 | -0.447 |
| Insulin Receptor Signaling | 0 | 0.029 | -2 |
| Neurotrophin/TRK Signaling | 0 | 0.026 | #NUM! |
| Integrin Signaling | 0 | 0.0294 | 0 |
| Death Receptor Signaling | 0 | 0.0426 | #NUM! |
| PPAR Signaling | 0 | 0.0667 | 1.89 |
| IGF-1 Signaling | 0 | 0.0098 | #NUM! |
| Dopamine Receptor Signaling | 0 | 0.026 | #NUM! |
| TGF-β Signaling | 0 | 0.0426 | #NUM! |
| Apoptosis Signaling | 0 | 0.0485 | 0 |
| Notch Signaling | 0 | 0.0541 | #NUM! |
| NF-κB Signaling | 0 | 0.016 | -2.449 |
| VEGF Signaling | 0 | 0.0208 | #NUM! |
| Hypoxia Signaling in the Cardiovascular System | 0 | 0.0132 | #NUM! |
| T Cell Receptor Signaling | 0 | 0.0297 | -1.508 |
| BMP signaling pathway | 0 | 0.023 | #NUM! |
| Phagosome Maturation | 0 | 0.0338 | #NUM! |
| Autophagy | 0 | 0.0144 | #NUM! |

| | | | |
|---|---|---|---|
| Macropinocytosis Signaling | 0 | 0.0133 | #NUM! |
| Sumoylation Pathway | 0 | 0.0103 | #NUM! |
| GP6 Signaling Pathway | 0 | 0.0242 | #NUM! |
| IL-7 Signaling Pathway | 0 | 0.026 | #NUM! |
| Sirtuin Signaling Pathway | 0 | 0.00743 | #NUM! |
| Opioid Signaling Pathway | 0 | 0.0593 | -0.277 |
| Iron homeostasis signaling pathway | 0 | 0.053 | #NUM! |
| Th17 Activation Pathway | 0 | 0.0327 | -1 |
| Endocannabinoid Developing Neuron Pathway | 0 | 0.0317 | #NUM! |
| Endocannabinoid Cancer Inhibition Pathway | 0 | 0.014 | #NUM! |
| Apelin Pancreas Signaling Pathway | 0 | 0.0222 | #NUM! |
| Apelin Cardiac Fibroblast Signaling Pathway | 0 | 0.0435 | #NUM! |
| Apelin Cardiomyocyte Signaling Pathway | 0 | 0.0515 | 0.447 |
| Apelin Endothelial Signaling Pathway | 0 | 0.036 | #NUM! |
| Apelin Muscle Signaling Pathway | 0 | 0.0417 | #NUM! |
| BAG2 Signaling Pathway | 0 | 0.0123 | #NUM! |
| FAT10 Signaling Pathway | 0 | 0.037 | #NUM! |
| FAT10 Cancer Signaling Pathway | 0 | 0.04 | #NUM! |
| T Cell Exhaustion Signaling Pathway | 0 | 0.0201 | -2.449 |
| Systemic Lupus Erythematosus In T Cell Signaling Pathway | 0 | 0.0305 | -1.387 |
| Systemic Lupus Erythematosus In B Cell Signaling Pathway | 0 | 0.0471 | -1.633 |
| Senescence Pathway | 0 | 0.0493 | 0 |
| White Adipose Tissue Browning Pathway | 0 | 0.0515 | 0.378 |
| Inhibition of ARE-Mediated mRNA Degradation Pathway | 0 | 0.0519 | -0.707 |
| Hepatic Fibrosis Signaling Pathway | 0 | 0.0565 | -1.147 |
| BEX2 Signaling Pathway | 0 | 0.013 | #NUM! |
| Xenobiotic Metabolism General Signaling Pathway | 0 | 0.0455 | 2.449 |
| Xenobiotic Metabolism CAR Signaling Pathway | 0 | 0.0592 | 0.632 |
| Xenobiotic Metabolism PXR Signaling Pathway | 0 | 0.0643 | 0.905 |
| Insulin Secretion Signaling Pathway | 0 | 0.0418 | -1.414 |
| Semaphorin Neuronal Repulsive Signaling Pathway | 0 | 0.0278 | #NUM! |
| Regulation Of The Epithelial Mesenchymal Transition In Development Pathway | 0 | 0.0471 | 0 |
| Kinetochore Metaphase Signaling Pathway | 0 | 0.0275 | #NUM! |
| Coronavirus Replication Pathway | 0 | 0.0526 | #NUM! |
| MSP-RON Signaling In Cancer Cells Pathway | 0 | 0.0294 | 0 |
| Calcium Signaling | 0 | 0.0574 | 0.707 |
| GM-CSF Signaling | 0 | 0.0143 | #NUM! |
| Ephrin Receptor Signaling | 0 | 0.04 | #NUM! |
| Ferroptosis Signaling Pathway | 0 | 0.0312 | -1 |
