## Supplementary material for "Scalable Generation of Universal hiPSC-Derived Vascular Progenitor Cells for Safe and Sustained Revascularization in Chronic Limb-Threatening Ischemia": Supplemental Table 4.pdf

| Antibody | Clone | Fluorochrome | Dilution | Company | Cat Number | Assay |
|---|---|---|---|---|---|---|
| Live/Dead | NA | Near IR | 1:1000 | ThermoFisher | L10119 | Flow cytometry |
| CD31 | WM59 | eFluor450 | 1:100 | eBiosciences | 48-0319-42 | Flow cytometry |
| CD43 | 1G10 | FITC | 1:50 | BD Biosciences | 555475 | Flow cytometry |
| CD144 | REA199 | APC | 1:50 | Miltenyi | 130-125-985 | Flow cytometry |
| CD140b | REA363 | PE | 1:50 | Miltenyi | 130-123-772 | Flow cytometry |
| CXCR4 | REA649 | FITC | 1:50 | Miltenyi | 130-117-370 | Flow cytometry |
| CD271 | REA844 | APC | 1:50 | Miltenyi | 130-112-602 | Flow cytometry |
| IgG1 | NA | eFluor450 | 1:50 | eBiosciences | 48-4714-82 | Flow cytometry |
| IgG1 | NA | FITC | 1:50 | BioLegend | 400110 | Flow cytometry |
| IgG1 | NA | PE | 1:50 | BioLegend | 400114 | Flow cytometry |
| IgG1 | NA | APC | 1:50 | Miltenyi | 130-113-446 | Flow cytometry |
| Isolectin-B4 | NA | AF647 | 1:200 | Invitrogen | I32450 | IF |
| aSMA | 1A4 | FITC | 1:500 | Sigma | F3777 | IF |
| dystrophin | NA | unconjugated | 1:250 | Proteintech | 12715-1-AP | IF |
| Ku80 | C48E7 | unconjugated | 1:400 | Cell Signaling | 2180 | IF |
| Fc Block | NA | unconjugated | 1:20 | Miltenyi | 130-059-901 | Flow cytometry |
| HLA-ABC | G46-2.6 | APC | 1:100 | BD Biosciences | BDB562006 | Flow cytometry |
| HLA-E | REA1031 | APC | 1:100 | Miltenyi | 130-117-402 | Flow cytometry |
| HLA-DP,DQ,DR | Tu39 | APC / Fire 750 | 1:100 | Biolegend | 361711 | Flow cytometry |
| CD40 | HB14 | APC | 1:100 | Biolegend | 313008 | Flow cytometry |
| CD86 | IT2.2 | PE | 1:100 | Biolegend | 305405 | Flow cytometry |
| CD80 | L307.4 | PE-Cy7 | 1:100 | BD Biosciences | 561135 | Flow cytometry |
| CD274 | MIH1 | PE | 1:100 | BD Biosciences | 1557924 | Flow cytometry |
| CD273 | MIH18 | APC-Cy7 | 1:100 | Biolegend | 345515 | Flow cytometry |
| FasL | NOK-I | PE-Cy7 | 1:100 | Biolegend | 306417 | Flow cytometry |
| VCAM1 | 429 (MVCAM.A) | APC-Vio770 | 1:100 | Miltenyi | 130-104-128 | Flow cytometry |
| ICAM1 | HA58 | PE-Cy7 | 1:100 | Biolegend | 353115 | Flow cytometry |
| CD142 | NY2 | PE | 1:100 | Biolegend | 356203 | Flow cytometry |
| CD46 | TRA-2-10 | APC-Cy7 | 1:100 | Biolegend | 352409 | Flow cytometry |
| CD59 | H19 | PE | 1:100 | Biolegend | 304707 | Flow cytometry |
| CD55 | JS11 | PE-Cy7 | 1:100 | Biolegend | 311314 | Flow cytometry |
| E-selectin | HAE-1f | PE-Cy7 | 1:100 | Biolegend | 336015 | Flow cytometry |
| P-selectin | AK4 | APC-Cy7 | 1:100 | Biolegend | 304943 | Flow cytometry |
| CD45 | 2D1 | PE-Cy7 | 1:100 | Fisher | 25-9459-42 | Flow cytometry |
| CD4 | RPA-T4 | PE | 1:100 | Fisher | 12-0049-42 | Flow cytometry |
| CD8 | RFT8 | APC-Cy7 | 1:100 | Fisher | A15448 | Flow cytometry |
